# Native Hydrogen/Deuterium Exchange Mass Spectrometry of Structured DNA Oligonucleotides

**DOI:** 10.1101/848598

**Authors:** Eric Largy, Valérie Gabelica

## Abstract

Although solution hydrogen-deuterium exchange mass spectrometry (HDX/MS) is well-established for the analysis of the structure and dynamics of proteins, it is currently not exploited for nucleic acids. Here we used DNA G-quadruplex structures as model systems to demonstrate that DNA oligonucleotides are amenable to in-solution HDX/MS in native conditions. In trimethylammonium acetate solutions and in soft source conditions, the protonated phosphate groups are fully back-exchanged in the source, while the exchanged nucleobases remain labeled without detectable back-exchange. As a result, the exchange rates depend strongly on the secondary structure (hydrogen bonding status) of the oligonucleotides, but neither on their charge state nor on the presence of non-specific adducts. We show that native mass spectrometry methods can measure these exchange rates on the second to the day time scale with high precision. Such combination of HDX with native MS opens promising avenues for the analysis of the structural and biophysical properties of oligonucleotides and their complexes.

## INTRODUCTION

In-solution HDX/MS is increasingly used to study the structural dynamics and interactions of proteins, in particular since the advent of automated systems. It relies on measuring exchange rates of backbone amide hydrogens in a deuterium-rich buffer, because these rates are heavily dependent on hydrogen-bonding and solvent accessibility.^1–9^ Since all amino-acid (except prolines) possess an amide hydrogen, HDX/MS can effectively cover the entire sequence. HDX/MS is particularly valuable for proteins that are not amenable to X-Ray diffraction or NMR experiments, because of *e.g.* their large size, high dynamics, or limited availability. In this manuscript, we show that HDX/MS is also particularly well suited to study structured oligonucleotides (Figure 1).

**Figure 1.**
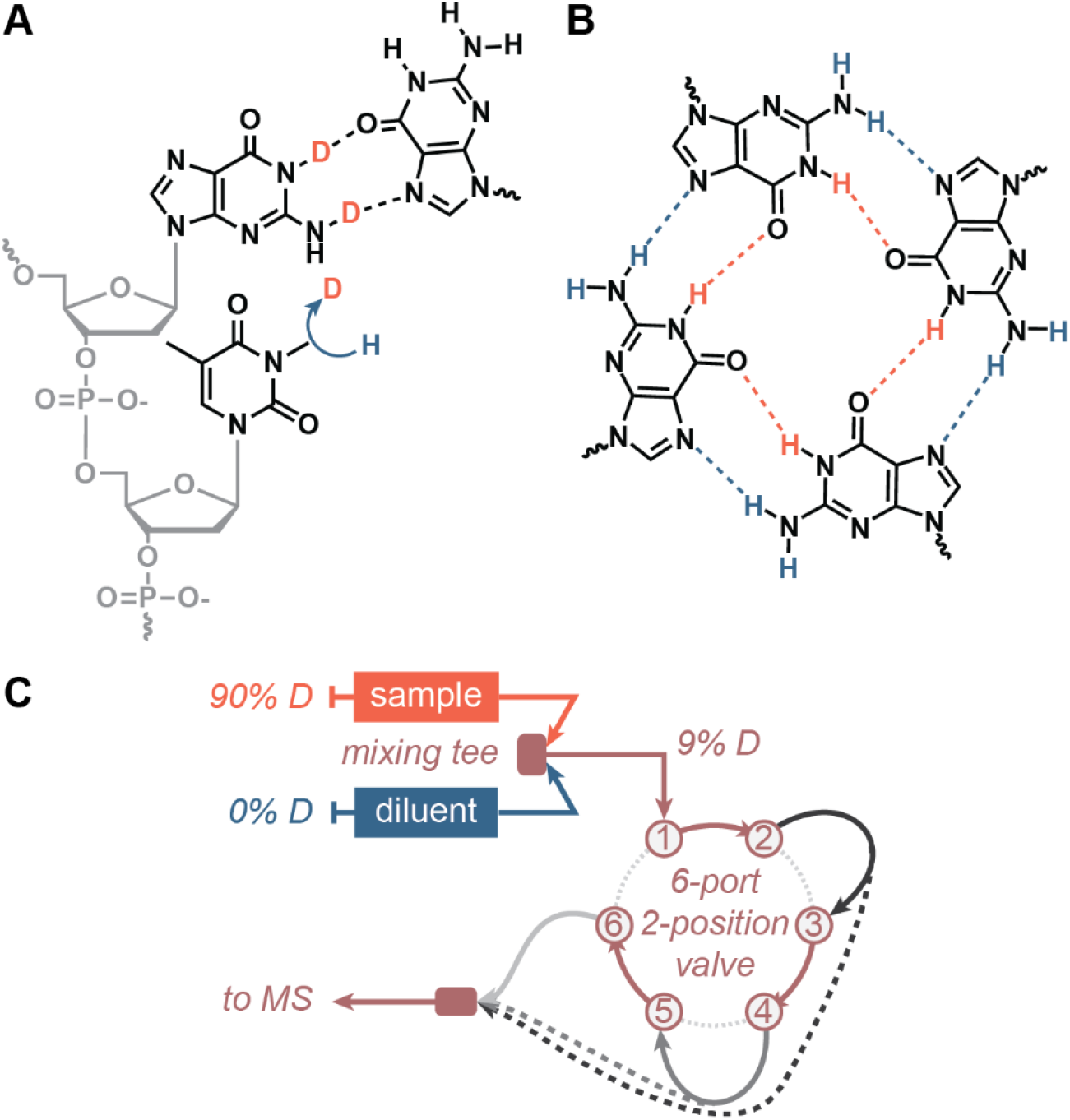
(A) HDX relies on the differential exchange of hydrogen and deuterium as a function of the hydrogen bonding status (herein a D to H exchange of nucleobases). (B) G-quadruplexes were used as model samples because the G-quartets they contain are formed by a network of eight imino (orange) and AMINO (blue) hydrogen bonds that can be exchanged over a large time range. (C) To monitor fast-exchanging sites (1—300 s), a continuous flow setup was used, making use of a 6-port, 2-position valve to easily cycle through capillary sizes (shades of grey; dashed lines illustrate direct coupling of a capillary to the MS source to shorten the mixing time).

Oligonucleotides contain several exchangeable sites: the 5’- and 3’-OH termini, the 2’-OH of RNA riboses (not explored herein), the phosphates, and the amino and imino hydrogens of bases. The -OH termini obviously do not cover the whole sequence, and are not good markers of the structure. The phosphate groups do cover the full sequence, but given their very low p*K*_a_ (~ 1), all phosphate groups are fully deprotonated at physiological pH.^10^ Some are neutralized during the electrospray process, but in such case may not provide clear information on the oligonucleotide’s structure and dynamics in the bulk solution. The nucleobases are involved in secondary structure formation via hydrogen bonds (Figure 1A), and given that each base contains at least one exchangeable site (dG: 3, dA and dC: 2, dT: 1), they theoretically suffice to cover the whole sequence.

Hydrogen exchange of nucleic acids with deuterium or tritium was in fact studied early on. Some of the earliest attempts at characterizing dsDNA breathing fluctuation on the minute time scale were made in the 1960s by H-T exchange quantified by scintillation counting.^11–13^ The exchange was however too fast to resolve the surface sites accessible in the grooves. From the 1970s, NMR was used to obtain exchange rates at the nucleotide level. However, as nucleic acids contain only four distinct nucleotides, determining the exchange rate of a particular nucleotide is only possible for short analytes (e.g. 10 bp), or those selected for their particularly well-resolved NMR peaks.^14^

DNA secondary structures are not limited to the Watson-Crick double helix. Nucleic acids can form a variety of non-canonical structures such as G-quadruplexes, triplexes or i-motifs,^15,16^ which either have biological roles^17,18^ or can be exploited in artificial nucleic acid constructs, for example aptamers.^19–22^ G-quadruplexes are formed in guanine-rich sequences by the stacking of a minimum of two G-quartets. A G-quartet (or tetrad) is a quasi-coplanar association of four guanines linked by a network of eight Hoogsteen hydrogen bonds through their imino and amino hydrogens (Figure 1B). A cation, typically potassium, is coordinated between each pair of G-quartets, and strongly contributes to the stability of the structure. In a G-quadruplex with *n* G-quartets, *n*-1 cations are generally specifically coordinated.^23^ G-quadruplexes are extremely polymorphic: their topology is defined by the strand stoichiometry, the relative orientation of their strands (relative to their 5’-to 3’polarity), number of G-quartets, and the length and geometry of the loops bridging tracts of guanines.^16,24^ The fact that not every guanine is necessarily part of a G-quartet makes structures difficult to predict, and is often a cause of the polymorphism.^24,25^

HDX/^1^H-NMR is frequently applied on G-quadruplexes.^26^ Usually, oligonucleotide samples are folded in a non-deuterated buffer, freeze-dried, then re-dissolved with D_2_O just prior to ^1^H-NMR acquisition. As the imino protons are exchanged with deuterium, the intensity of the corresponding peak decreases.^26^ However, this approach is almost always applied in a binary fashion, differentiating fast vs. slow exchanging sites to identify the protons protected in G-quartets and stacking interfaces.^26–29^ Moreover, it is limited to structures that give well-resolved NMR peaks, and the signal integration may require the use of a high-magnetic-field spectrometer.^30^ The ^1^H-NMR exchange kinetics of some imino sites of a c-kit2 G-quadruplex variant were reported, with a relatively large experimental dead time (about 4 minutes, during which one quartet was fully exchanged).^30^

More generally, both NMR and X-ray crystallography can provide high-resolution structures of oligonucleotides. Unfortunately, they are limited in their ability to deal with several conformers in equilibrium, may promote the formation of higher-order structures that might not be biologically relevant, and require expensive equipment. We and others have shown that native MS helps studying individual oligonucleotide species from mixtures of conformers.^10,31–33^ It relies on the direct stoichiometry observation and quantification of each peak by m/z separation. In particular, the measurement of cation stoichiometry provides information on structures and folding pathways of G-quadruplexes.^24,34,35^

Herein, we show that coupling HDX with native MS has the potential to combine the benefits of the two approaches. We validated that the bases, but not the phosphates, are isotopically exchanged proportionally to the solution deuterium content without in-source back-exchange. We developed analytical tools to precisely monitor the exchange over several orders of magnitude of time scales (seconds to days) (Figure 1C). Finally, we demonstrate on DNA G-quadruplexes that the measured exchange rates depend on the hydrogen bonding status of nucleobases, thus allowing one to probe nucleic acid folding.

## EXPERIMENTAL SECTION

### Materials

Oligonucleotides were purchased from Eurogentec (Seraing, Belgium) or IDT (Leuven, Belgium), in desalted and lyophilized form. Their sequence and melting temperatures are gathered in Table 1 and Table S1. Trimethyl ammonium acetate (TMAA) (1 M aqueous solution, HPLC grade), ammonium acetate (5 M aqueous solution, molecular biology grade), KCl (99.999% trace metal basis), and D_2_O (99.9% D atom) were obtained from Sigma-Aldrich (Saint-Quentin Fallavier, France). Samples were prepared in nuclease-free water (Ambion, Life technologies SAS, Saint-Aubin, France). Data analysis was performed with OriginPro 2018 (OriginLab, Northampton, MA, USA) and RStudio 1.2.1335 (RStudio Team, Boston, MA, USA).

**Table 1.** Selected oligodeoxynucleotide sequences.^a^

| Name | Sequence (5' to 3') | Structure<sup>b</sup> |
| --- | --- | --- |
| **T12** | T<sub>12</sub> | unfolded |
| **T24** | T<sub>24</sub> | unfolded |
| **23nonG4** | TG<sub>3</sub>ATGCGACA(GA)<sub>2</sub>G<sub>2</sub>ACG<sub>3</sub> | unfolded<sup>25</sup> |
| **24nonG4** | TG<sub>3</sub>ATGCGACA(GA)<sub>2</sub>G<sub>2</sub>ACG<sub>3</sub>A | unfolded<sup>25</sup> |
| **N-myc** | TAG<sub>3</sub>CG<sub>3</sub>(AG<sub>3</sub>)<sub>2</sub>A<sub>2</sub> | G-quadruplex<sup>36</sup> |
| **26CEB** | A<sub>2</sub>G<sub>3</sub>TG<sub>3</sub>TGTA<sub>2</sub>GTGTG<sub>3</sub>TG<sub>3</sub>T | G-quadruplex<sup>37</sup> |
| **T30177-TT** | T<sub>2</sub>GTG<sub>2</sub>(TG<sub>3</sub>)<sub>3</sub>T | G-quadruplex<sup>38</sup> |
| **TBA** | G<sub>2</sub>T<sub>2</sub>G<sub>2</sub>TGTG<sub>2</sub>T<sub>2</sub>G<sub>2</sub> | G-quadruplex<sup>39</sup> |
| **24TTG** | TTG<sub>3</sub>(TTAG<sub>3</sub>)<sub>3</sub>A | G-quadruplex<sup>40</sup> |
| **222T** | T(G<sub>3</sub>T)<sub>4</sub> | G-quadruplex<sup>24</sup> |
| **ds26** | CA<sub>2</sub>TCG<sub>2</sub>ATCGA<sub>2</sub>T<sub>2</sub>CGATC<sub>2</sub>GAT<sub>2</sub>G | dsDNA |

<sup>a</sup> All sequences are reported in Table S1
<sup>b</sup> Guanines involved in G-quartet formation are bolded

### Sample preparation

The deuterated (pre-exchange) samples contained 50 µM DNA (from 1 mM stock solutions in D_2_O) and 90% D_2_O, 1 mM KCl, 100 mM TMAA (from 10 mM KCl and 1 M TMAA stock solution in water, pH = 7.0 (see Supporting Information). Melting temperatures were determined by UV-melting (see Supporting Information; Figure S1). The abundance of deuterium was adjusted by adding the appropriate amounts of D_2_O or H_2_O. The samples were annealed at 85°C for 5 min (ensuring proper folding while all sites are exchanged to 90% D), and stored at 4°C for at least a night before use. The diluent solution used to trigger the D-to-H exchange was composed of 1 mM KCl and 100 mM TMAA in H_2_O. The 9%-D references were prepared from samples incubated at 85°C for 5 min to ensure that they reflect the exact same mixing ratio than in the D-to-H exchange measurements.

### HDX/MS experiments

Electrospray ionization mass spectrometry (ESI-MS) experiments were performed on a Thermo Orbitrap Exactive mass spectrometer calibrated daily, and operated in negative mode, on a 400 – 3000 m/z scan range, using the 50 000-resolution setting, with the following tuning parameters: 2.95 kV capillary, 200°C capillary temperature, sheath gas: 55, aux gas: 5, heater temperature: 25°C, tube lens voltage: −200V, skimmer voltage: −14.5 V (see Supporting Information). For denaturing conditions, the capillary temperature was set to 240°C. For softer conditions, the capillary temperature was set at 170°C, and the tube lens voltage at −190 V. The relative softness of all three settings was assessed by infusing a 10-µM solution of the [G_4_T_4_G_4_]_2_ G-quadruplex in 100 mM ammonium acetate (Figure S2).^41^ It is best practice to perform this control when using a different instrument and/or a different tune. All data was acquired with Thermo Xcalibur 2.2. The HDX summary and data tables, adapted from the recommendations by Masson *et al.*,^42^ are available as Supporting Information.

Performing the experiments in the D-to-H direction allows working in a mostly non-deuterated solvent (9% deuterium), which is closer to physiological conditions and reduces the consumption of D_2_O. The exchange was triggered by mixing the deuterated sample (90% D) with nine volumes of the same buffer, albeit non-deuterated (0% D), *i.e.* 100 mM TMAA, 1 mM KCl in pure H_2_O. In manual mixing experiments, the sample and diluent were thoroughly mixed with a vortex and the resulting solution (9% D) was transferred in a 500-µL glass syringe, which was then used to flush the tubing dead volume, before being installed on a syringe pump operated at 5 µL/min. The signal was integrated for a minute for each time point, and the mean labelling time was reported. In continuous-flow experiments (Fig. 1C), the deuterated sample and the diluent were pumped at a 1:9 ratio towards a PEEK high pressure static mixing tee, equipped with a UHMWPE frit (IDEX Health & Science, Oak Harbor, WA, USA). The resulting mixed solution was brought to the MS source through PEEK tubings of different volumes (by changing both the length and internal diameter) to generate different mixing times. To vary the mixing time in a robust and simple way, an MX Series II 2 Position/6 Port UltraLife Switching Valve (IDEX Health & Science, Oak Harbor, WA, USA) was used (see Figure S3). The accuracy of the flow rates was verified by measuring the volumes delivered in two minutes. All experiments were performed at room temperature, i.e 22±1°C. The signal was integrated for a minute for each time point. The labelling time *t*_*l*_ = *V*/*F*, corresponds to the time spent by the sample in the mixing fluidics (from the mixing tee to the MS source) of volume *V*, at a flow rate *F*. The integration time can be increased to enhance the signal-to-noise ratio where necessary.

Raw HDX/MS data were analyzed manually, by visually selecting the isotope distributions for all exchange time points then calculating the centroid mass (Figure S4). Exchange curves can be plotted as the *m/z* centroid of the isotopic distributions of interest against time *t* (Figure S5). Centroids were converted to a number of unexchanged sites (*NUS*) using Equation 1.

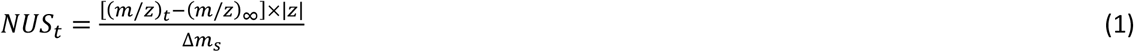

It is defined as the measured mass shift between the sample centroid mass (m/z)_t_ at a time t and that of the fully-exchanged reference centroid (m/z)_∞_, converted to mass units by multiplying by |z|, and divided by the apparent isotopic mass shift for a single site (Δm_s_). The value of (m/z)_∞_ will be discussed in the text. The value of Δm_s_ depends on the pre- and post-mixing isotopic compositions of the solvent, because the apparent isotopic mass of each site is the sum of the isotopic masses of hydrogen and deuterium (m_H_ and m_D_, respectively), multiplied by their respective molar fractions in the medium (H or D),

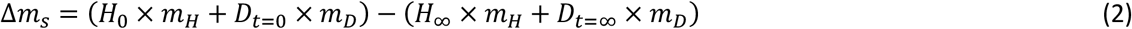

Which simplifies into Equation 3 because *H* = 1 − *D*:

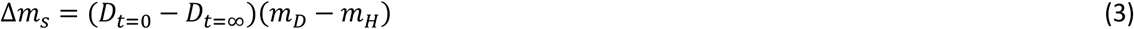

Unless otherwise mentioned, all experiments were performed here with D_t=0_ = 0.9 and D_t=∞_= 0.09, yielding Δm_s_ ≈ 0.815.

The exchange of a single site can typically be fitted with an exponential decay model. All sites are independent from one another and, unless there is evidence of cooperative unfolding, exchange at their respective rates.^4,6^ Exchange curves acquired at the intact level therefore consists of the combination of the simultaneous exponential decay of all n sites,^5^ and thus might not be fitted with a single exponential decay model. Some sites nevertheless display similar exchange rates, which reduces the total number of terms necessary to fit the data. Non-linear fitting of the NUS = f(t) exchange curves was therefore carried out with Equation 4, where NUS_0_ is the offset, k is the rate constant, N is the number of sites in each group, and j the number of exponentials.

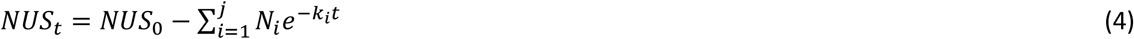

Over large time ranges, non-linear fitting was typically performed with j = 3, and was reduced to j = 2 or 1 for fast exchanging species and/or when the exchange time range was restricted to either 1 s—5 min or 2 min—90 min, as is also usually observed for peptides in protein bottom-up experiments.^4^

## RESULTS AND DISCUSSION

### The exchangeable sites are located on the nucleobases

In DNA, the acquisition of the final charge is a partial neutralization of the phosphate groups by protons. The total number of sites nX that are potentially exchanged in a DNA oligonucleotide detected by ESI-MS in negative mode is therefore equal to the sum of all bases imino and amino groups, neutralized phosphate groups nP, and the -OH termini T. In an n-mer oligonucleotide capped by two OH termini [dX_n_]^z-^, there are n-1 phosphates and therefore nP = n – 1 – z. Furthermore, in MS of oligonucleotides, it is frequent to observe non-specific cation binding to the phosphates. We will suppose that the charge balance is ensured by phosphate neutralization. The number of neutralized phosphates of an n-mer oligonucleotide binding m cations C of absolute charge y is given by Equation 5.

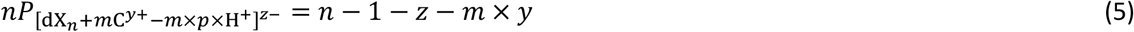

To determine the number of actually exchanged sites, we acquired the mass spectra of several oligonucleotides (dT_12_, dT_24_, 23nonG4, and N-myc) across a large range of deuterium content (10 to 90%; Figures S6 to S9). These sequences differ by their length (12 to 24 nucleotides), composition (polydT vs. mixed-base nucleotides), and only the latter is folded into a specific structure at room temperature, in 100-mM TMAA, 1-mM KCl solutions, specifically a G-quadruplex of T_m_ = 55.3 ± 0.3°C (n = 5; Figure S1). To ensure a quantitative isotopic exchange, all oligonucleotides were incubated 5 min at 85°C prior to MS measurements (N-myc is entirely unfolded at equilibrium above 72°C). These measurements were performed on different days by two distinct operators, using the MS tune used in all HDX experiments presented herein, as well as both a softer tune and a denaturing tune (see the experimental section) to check for possible gas-phase back-exchange. The centroid masses were plotted as a function of solution deuterium content and overlaid with the theoretical centroid values calculated for all possible number of exchangeable sites (0 to nX) and deuterium content (0—100%) to readily visualize the number of exchanging sites (Figure 2A and S10).

**Figure 2.**
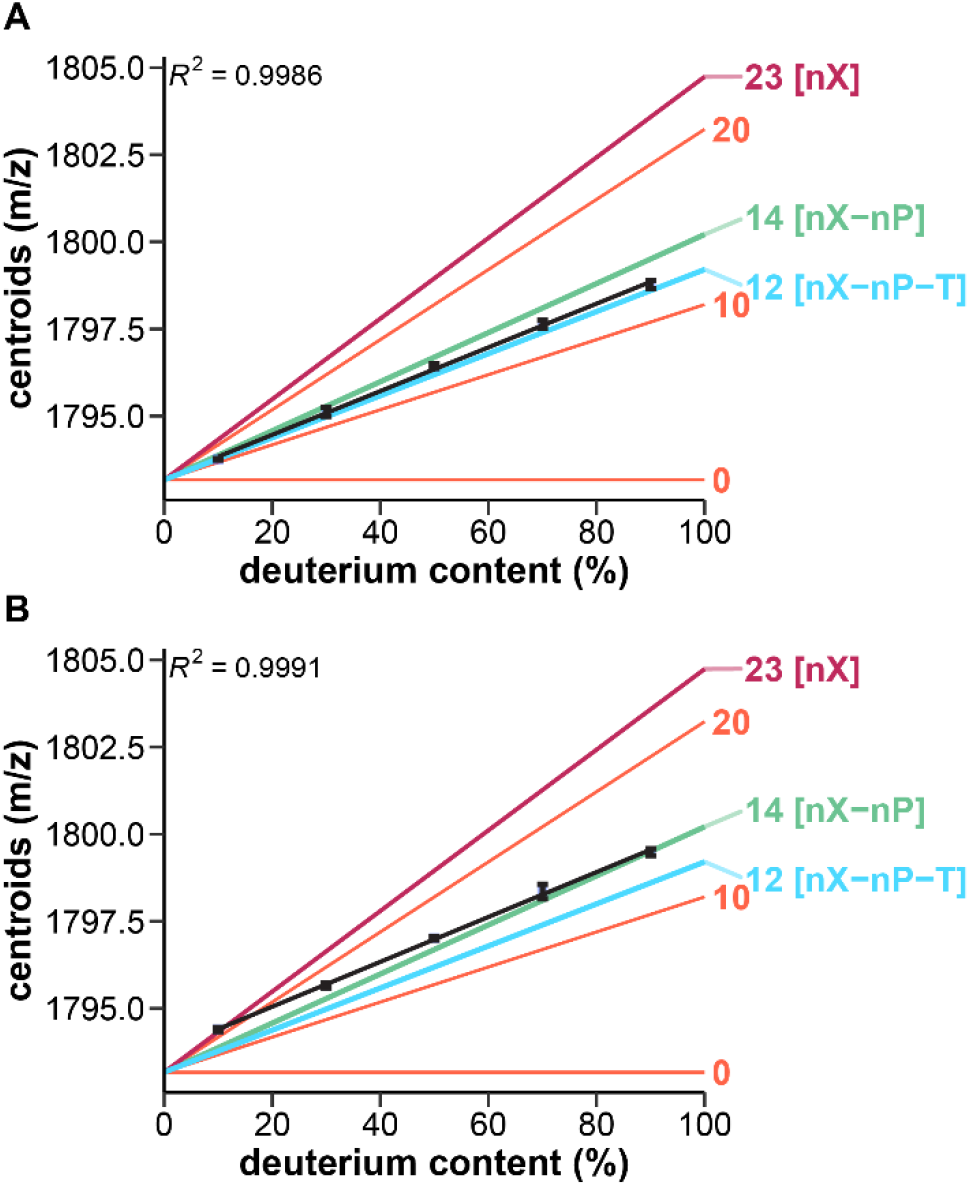
Exchangeable sites in T12 for z = 2-, in 100 mM TMAA, 1 mM KCl, using the HDX tune in A) ambient conditions and B) in a D_2_O-doped source atmosphere: Measured centroid masses against the solution deuterium content (n = 2, standard deviation as error bars, linear fit as black lines), compared to theoretical m/z: maximum number of potentially exchangeable sites (nX; purple), no phosphate exchanged (nX–nP; green), no phosphate and termini exchanged (nX–nP–T; cyan), and every ten sites (coral).

A linear relationship between the solution deuterium abundance and the centroid mass is observed for all oligonucleotides, for data replicated with the standard and softer tunes. The slopes are consistent with a statistical exchange of a defined number of unprotected sites, in solution, which follows the binomial law assuming the exchange of different sites is uncorrelated.^5,43,44^ However, the number of actually exchanging sites is lower than the total number of theoretically exchangeable sites listed above. It is closest to the number of bases’ exchangeable sites (*nX* – *nP* – *T*), although a few more sites are exchanged for T12 (one) and T24 (four).

These results suggest that the phosphate groups are not (or only weakly) deuterated, regardless of the solution D_2_O content. Katta and Chait demonstrated that labile hydrogens (from protein side chains) can be back-exchanged in electrosprayed solutions from water vapor of the ambient atmosphere.^45^ To test this hypothesis, we monitored the exchange in ambient air, with additional H_2_O vapor in the source (Figures S11 and S12), and with additional D_2_O vapor in the source (Figures S13 and S14), coming from droplets lying at the bottom of the source during the experiment. Enriching the source with D_2_O leads to increased deuteration (Figure 2B and Figures S15 to S18). Moreover, although there is still a linear relationship between the centroid mass and the solution D_2_O content, the data is not consistent anymore with a defined number of solution-exchanging sites, highlighting the presence of in-ESI exchange.^46^ However, the number of exchanged sites does not change when increasing the source H_2_O partial vapor pressure. We also noted that using ammonium acetate instead of trimethylammonium acetate yields 15% more exchanged sites for both T12 and T24 (Figures S19 to S21). The buffer choice thus influences back-exchange. Trimethylammonium is a protic, tertiary ammonium ionic liquid (IL), less volatile and acidic than ammonium (boiling point ≈ 3 vs. −33°C, p*K*_a_ = 9.25 vs. 9.80).^47^ The cation and anion of an IL can form an ion pair via hydrogen bonding, and some (including ammonium-based ILs such as TMAA) can H-bond with water as well, leading to altered viscosities and acidities.^47–50^ These differences in properties might alter the droplets’ composition during the electrospray mechanism, leading to changes in the neutralization of phosphates. The phosphate sites are therefore deuterated to an extent that depends on both the atmosphere isotope composition and buffer cation nature.

A possible explanation to the virtual absence of phosphate deuteration is that, in ambient conditions, the humidity in the source transfers protons (or deuterons in presence of D_2_O vapors) to remaining TMA (or ammonium) cations capping DNA, and then to the DNA phosphates, provided that the de-clustering finishes well after the ions escape the droplets (Figure 3B). Phosphate back-exchange could also originate from H_2_O/D_2_O vapors entrainment in droplets. It has been observed in ESI, although it is mostly suppressed when using a dry nebulizing gas.^51,52^ Change in the droplet deuterium content could lead to solution-phase exchange in the droplet, as evidenced by Gallagher and coworkers.^46^ This could explain the increased deuteration in presence of D_2_O vapors, particularly at low solution deuterium content where the alteration of the droplet composition would be more significant.

**Figure 3.**
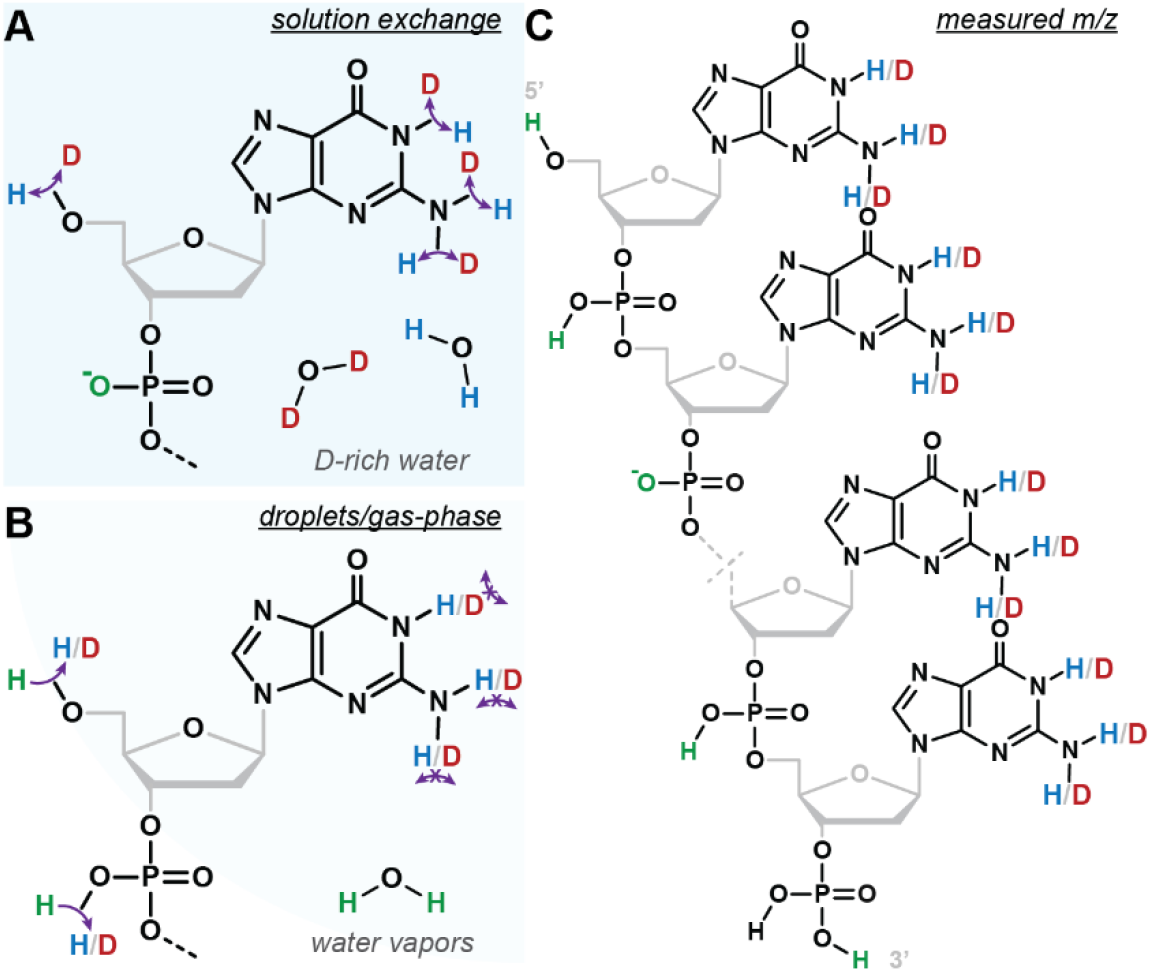
Exchangeable sites in native HDX/MS of TMAA-solutions of DNA oligonucleotides. A) Unpaired nucleobases exchange to reflect the isotopic solvent composition, whereas phosphates remain ionized until B) the electrospray process during which some (depending on the charge) are neutralized by either H or D. Ultimately they are back-exchanged by the protons from the ambient H_2_O vapors (unless the source was doped with D_2_O vapors as in Figure 2B) in the gas phase and/or droplets. Nucleobases do not back-exchange, leading to C) the final analyte in which the nucleobases reflect the solution exchange and the phosphate are either ionized or protonated.

Phosphate back-exchange is consistent with the fact that phosphates are the most acidic sites within nucleotides, leading to faster exchange than the base sites, both in solution and the gas-phase (RO_3_PO**H** > R_2_N**H** > RN**H**_2_).^53,54^ It is further corroborated by the fact that, although the number of neutralized (and potentially exchangeable) phosphates increases with decreasing charges (Equation 5), the number of effectively exchanged sites remains constant across charge states for all oligonucleotides. Phosphates within oligonucleotides anions were shown to exchange rapidly in the gas-phase.^55^ The exchange of a rigid G-quadruplex (in which base pairing and stacking is conserved) is faster than the corresponding single strand.^55^ Additionally, phosphates of 6-mer DNA polyanions can form H-bonds in the gas-phase,^56^ which could lead to decreased exchange rates. This may account for the differences between the G-quadruplex N-myc (base stacking and pairing in the gas-phase; all phosphates back-exchanged) and the unfolded T12 and T24 (amenable to phosphate H-bonding; some deuterated phosphates).

When infusing the samples in harsher source conditions, the number of exchanged sites decreases for all oligonucleotides, below the number of bases’s sites in some cases (N-myc and 23nonG4; Figures S22 to S26). Back-exchanging the bases is therefore possible as well, although it remains very limited, and is totally avoided when using sufficiently soft source conditions. Base exchange in the gas phase with H_2_O or D_2_O is slow (hour-scale at 5.0 × 10^−8^ Torr of D_2_O),^54^ because their gas-phase acidity difference is quite large (> 40 kcal/mol).^53,57,58^ The impact of organic co-solvents in samples on back-exchange was not explored herein, as it can alter the conformational equilibrium,^59^ and is not necessary to obtain adequate signal-to-noise ratio. Methanol has already been shown to have little effect on oligonucleotide gas-phase exchange,^60^ and has been proposed to decrease in-droplet exchange.^46^

In summary, we have set up a robust approach and robust conditions to quantify the number of exchanging sites by MS, and demonstrated that it is consistent with the exchangeable sites being located on the nucleobases (Figure 3A), without gas-phase back-exchange if using relatively soft ionization conditions (Figure 3B). The virtual absence of phosphate deuteration on the final analyte using TMAA is not detrimental, and may in fact be considered as an advantage, as it streamlines the data analysis without information loss (Figure 3C).

### Implications for converting the *m*/*z* distribution into unexchanged sites equivalent and interpreting *NUS* values

In HDX/MS experiments, monitoring *m*/*z* shifts as a function of time is sufficient to obtain relative trends, but does not provide an absolute number-of-site information, and complicates the comparison across species of different masses. In protein HDX/MS studies, the “deuterium level” is frequently employed, *i.e.* the absolute number of *deuterium* incorporated, calculated using 0%- and 100%-D references.^4–6^ Here, we opted to convert the Δ*m/z* to a number of unexchanged sites (*NUS*) using Equation (1). It reports on the absolute number of non-exchanged sites, and provides a direct readout of the apparent number of sites involved in secondary structure formation, because these H-bonded sites should be protected from exchange. This greatly simplifies structural interpretations, because different structural motifs (e.g. base-pairs, quartets) have different contributions. In the case of G-quadruplexes, the imino groups involved in internal G-quartets are more protected than those from external quartets.^30^ This is clearly reflected in kinetic plots expressed in *NUS* values: TBA (2 quartets) does not have any long-lived sites (t_1/2_ < 1 min; Figure 4), 3-quartet structures have around four long-lived sites (Figure 4), and TG_5_T (5 quartets) contains twelve highly protected sites (Figure S5). Note that a *NUS* of 1 does not necessarily mean that one specific site is unexchanged, but rather that the overall, apparent amount of unexchanged is *equivalent* to one site (e.g. five 20%-unexchanged sites would give a *NUS* of 1).

**Figure 4.**
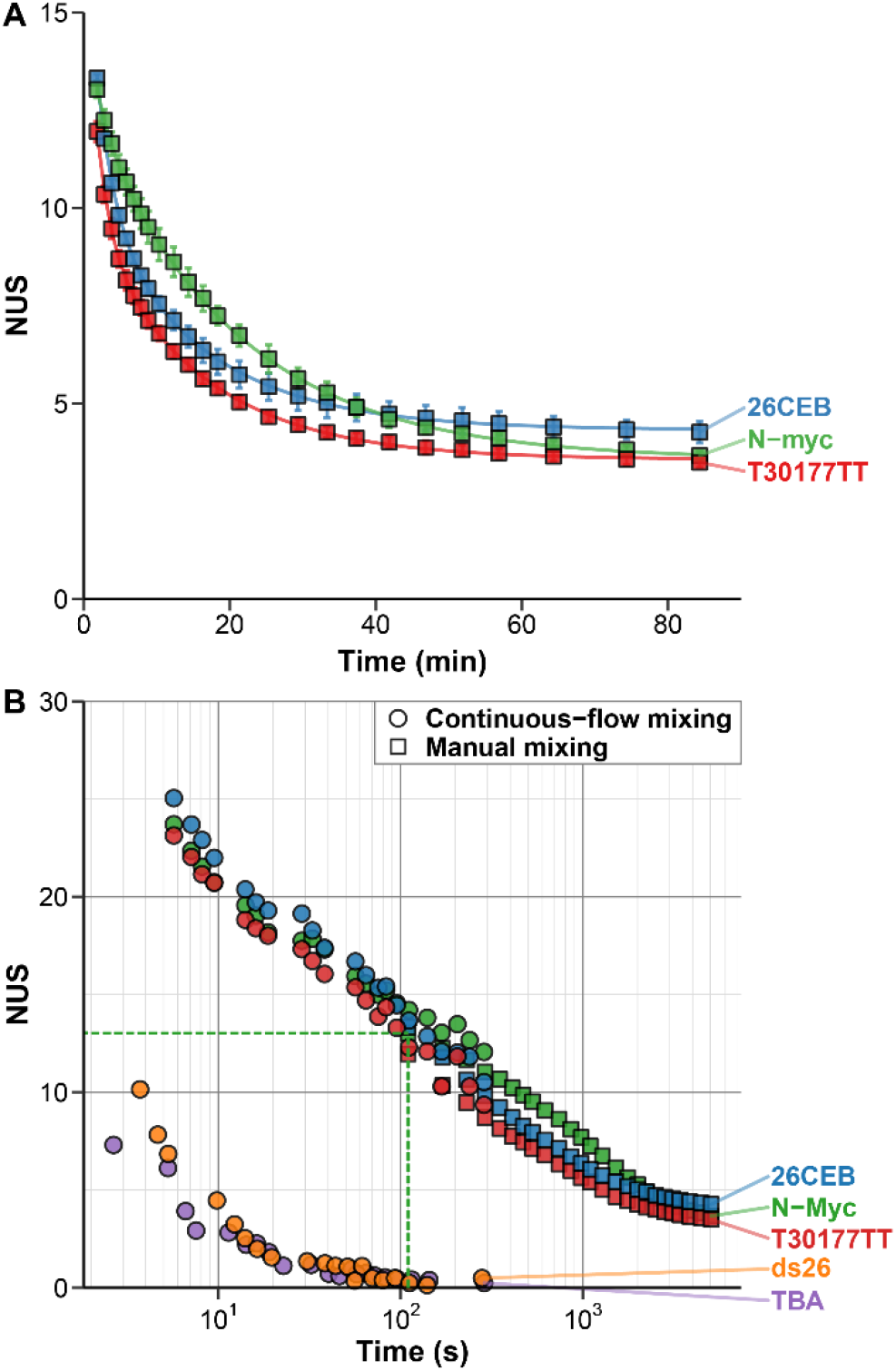
Exchange kinetics of oligonucleotides acquired by manual mixing (A and B; squares) and continuous-flow (B; circles) native HDX/MS (90% to 9% D) for five oligonucleotides. A) The error bars show the standard deviation across all species (z = 3 to 5-for N-myc and T30177TT, and 4 to 6-for 26CEB, K^+^ = 2 to 4) for three replicates. The lines are the results of the non-linear fitting using Equation 4 with j = 2. B) The dashed green lines indicate the position of the earliest N-Myc point acquired by manual mixing (110 s).

The *NUS* is independent from the number of sites, thus allowing to compare different analytes. The conversion to *NUS* values only requires the acquisition of a mock end-point (*m/z*_∞_ in Equation (1)), which mimics the state of the oligonucleotides upon exchange completion. This was achieved by incubating the exchanging samples for 5 minutes at 85°C, to ensure that all sites are available for rapid exchange. Following our findings on phosphate and terminal-OH back-exchange, we compared the experimentally acquired mock-end point spectra of a dozen oligonucleotides to the corresponding theoretical isotopic distributions, following the postulate that only the exchange of nucleobases is monitored (Figures S27 and S28). Mass errors down to the 10-ppm range were obtained, which highlights the satisfactory accuracy in deuterium content in the HDX/MS experiments described below, and further validates the localization of the exchanged sites on the bases.

### Exchange rates can be measured precisely on a large time scale, regardless of the charge state and adduct stoichiometry

To monitor the isotopic distributions upon exchange, the sample and diluent solutions can be manually mixed to trigger the exchange, then directly infused in the mass spectrometer (Figure 4A - squares). This gives access to time scales from a few minutes to several hours (Figure S5) or even days.

Relying on HDX/MS to study oligonucleotide structures and biophysics requires sufficient precision in measuring the exchange rates, and ideally no dependence on artifacts related to ionization such as the charge state or the formation of non-specific adducts. We identified several potential sources of errors on both the time and mass dimensions. Errors in masses may arise from the mass spectrometry measurement itself (mass spectrometer accuracy, inaccurate peak selection and integration, particularly in the case of partially overlapped isotopic distributions, weak signal and poor signal-to-noise ratio) as well as from inaccurate deuterium content before (sample preparation errors) and during exchange (inaccurate mixing ratio). The measurement of the mixing dead time (for manual-mixing experiments), and temperature variations during the exchange may also be sources of errors on the HDX rate.

To evaluate the precision of the method, two operators ran triplicate experiments on three G-quadruplex-forming oligonucleotides, namely N-myc, T30177-TT, and 26CEB. These are three-tetrad G-quadruplexes of different lengths (19—26 nucleotides) and number of exchanging sites (45—62). However, they have similar thermal stabilities (*T*_m_ = 52—55°C; Table S1 and Figure S1), and apparent exchange rates. For each replicate, the data of the three most abundant potassium stoichiometries from the three most intense charge states were examined (Figures S29 to S38). The mean *NUS* values, and their corresponding standard deviations and RSDs across replicates were calculated for each time point (Figure 4A, Figure S39A,B). Overall, the method has an excellent precision, with low mean RSD, whether all species (3.7%, 3.1%, and 4.4% for N-Myc, T30177-TT, and 26CEB, respectively) or only the most abundant ones (2.7%, 3.0%, and 4.6%) are considered (Table S2). The corresponding mass errors are inferior or equal to 0.2 Da (Table S3), which is on par with well-controlled protein HDX experiments (precise sample preparation, stable mass spectrometer calibration).^6,61–63^

Despite being quite similar (three-tetrad parallel quadruplexes of close stability), all three oligonucleotides can therefore be discriminated from each other using the exchange data. This can be achieved by visual inspection (Figure 4A), pairwise statistical comparison of individual time points (Table S4) or fitting parameters (Table S5). Comparing apparent exchange rates may not be reliable because it reduces numerous sites exchanging at their own rates to a few apparent rates, which masks part of the information and can lead to nonconclusive results (the 95% confidence intervals can overlap). The two former approaches are straightforward and the large number of data points acquired (compared to classical bottom-up approaches) allows to confidently conclude that these are three different exchange behaviors. The results on this set of oligonucleotides suggest that HDX/MS is well-suited for the quality control analysis of oligonucleotide folding, for instance during the production of structured oligonucleotides such as aptamers (batch-to-batch consistency, stability and stress testing), or when introducing modifications that may alter the structure (e.g. modified nucleotides, biotinylation, fluorophores).

In a typical native mass spectrum of oligonucleotide, a few charge states are detected (here, 3-to 6-depending on the sequence; Figure S29). We did not observe any significant differences in exchange rates between different charge states (Figure S39C). Hence, any or all charge states can be selected for data treatment; those of higher intensity and/or with fewer non-specific cation binding should thus typically be favored. Additionally, all four oligonucleotides bind specifically two K^+^ to form 3-tetrad quadruplexes, whereas all higher stoichiometries (i.e. 3 and 4 K^+^) arise from additional non-specific binding. No significant differences in the exchange rates between the 2, 3, and 4 K^+^ species were observed, across all replicates and charge states (Figure S39D), and therefore any or all can be used for data treatment. In absence of knowledge on cation specificity, it is however necessary to process each stoichiometry separately to assess which can be used, and possibly detect the coexistence of several conformers in solution.

In manual-mixing experiments, fast exchanging phenomena can be missed within the dead time. For instance, the N-myc sequence has 50 exchanging sites, but at the earliest continuous flow time point (110 s; signal integrated from 50 s to 170 s) only the equivalent of 13 sites remain unexchanged (Figure 4, dashed green line). To access faster exchange events, a continuous flow setup was employed, wherein the sample is continuously mixed with the exchanging buffer for a fixed duration. Continuous-flow mass spectrometry is used to study time-resolved events,^7,64^ and has already been employed in HDX/MS experiments by Wilson and Konermann.^65–67^ Another advantage is to allow recording a given mixing time point for as long as necessary. This is particularly interesting for analyzing low-concentration samples, as well as low-abundance and/or fast-exchanging species, for which real time analysis would require the extraction of mass spectra over a narrow time range, which could lead to poor S/N. Here, continuous flow was used to access exchange events ranging from around 1 second to 5 minutes (Figure 4B - circles). The exchange of 10 additional sites could be monitored for N-myc. Importantly, this enables the analysis of fast-exchanging species such as 2-tetrad G-quadruplexes (e.g. TBA; Figure S40) and dsDNA (e.g. ds26; Figure S41) that is not possible by manual-mixing. The results of the manual and continuous flow mixing techniques can be combined to obtain exchange curves over labeling times spanning several orders of magnitude (Figure 4B), as is recommended in protein HDX/MS,^42^ giving access to both weakly and strongly protected sites.

### Exchange rates on the second-to-day time scale are sensitive to oligonucleotide folding

To be used in structural and biophysical studies, HDX/MS of oligonucleotides must depend on site exchange availability, or *hydrogen bonding status*. A simple comparison can be made with 24TTG and 24nonG4, two sequences of close composition and same length but of different structures. The unfolded 24nonG4 oligonucleotide, whose sites are all available for exchange, is fully exchanged within a second, likely at the chemical exchange rates of the individual sites (Figure 5A). Conversely, the G-quadruplex 24TTG is still partially unexchanged after 5 minutes of incubation, most probably because some sites are engaged in hydrogen bonding and therefore protected from exchange.

**Figure 5.**
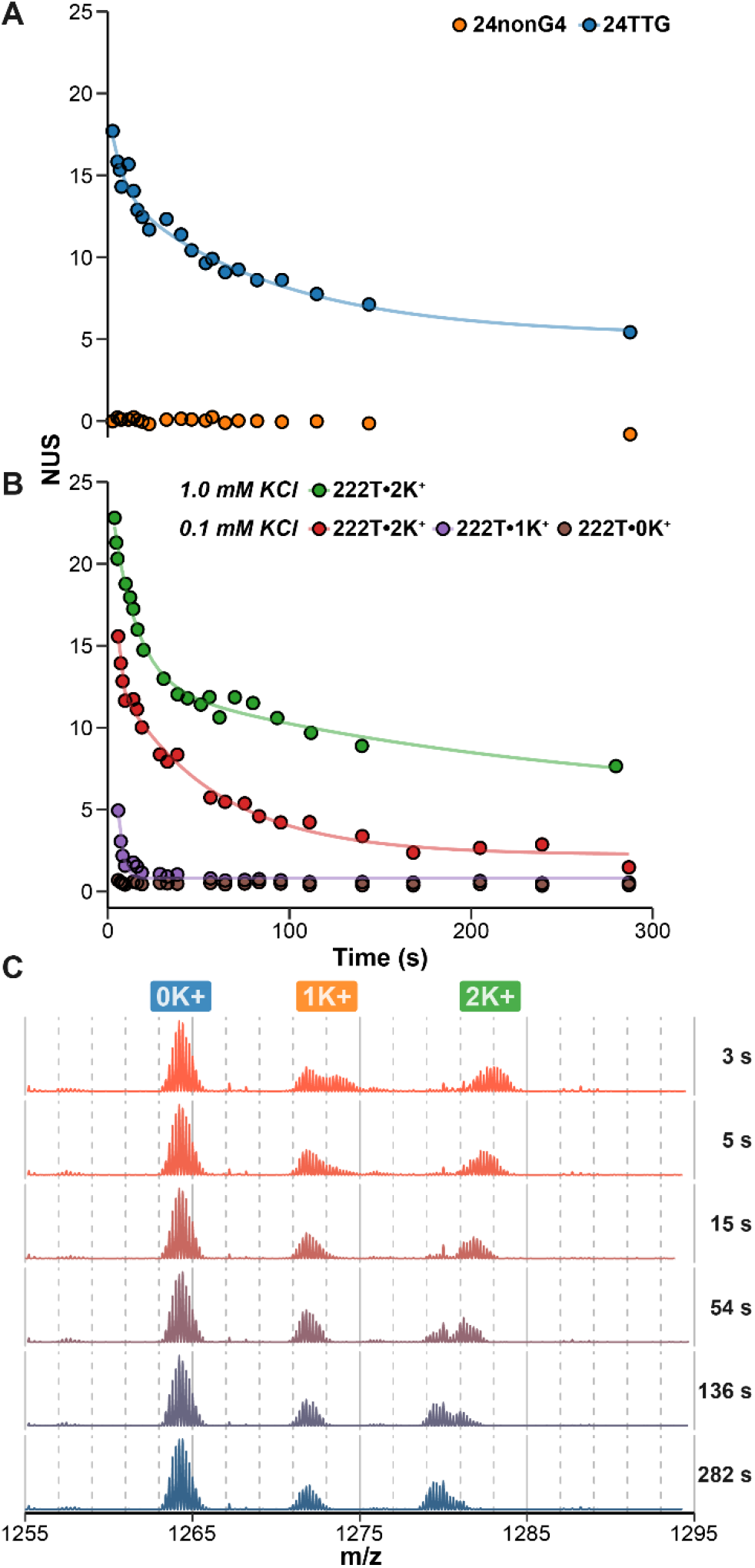
Hydrogen bonding status dependency of oligonucleotide HDX/MS. Exchange kinetics acquired by continuous-flow HDX (90% to 9% D) of A) unstructured (24nonG4) and structured (24TTG•2K^+^) oligonucleotides, and B) the same oligonucleotide (222T) in presence of 1.0 or 0.1 mM KCl, which leads to the formation of three distinct species (the G-quadruplexes 222T•2K^+^ and 222T•1K^+^, and the unfolded conformer 222T•0K^+^). The lines are the results of nonlinear fitting using Equation 4 with j = 1 or 2 for 2- and 3-tetrad species, respectively. C) Corresponding spectra for 222T in 0.1 mM KCl solution, zoomed on the 5-charge state. The K^+^-binding stoichiometry is indicated with labels. No exchange occurs in the explored time range for unfolded species (0 K^+^), unlike the G-quadruplexes species (≥ 1K^+^).

In the next example, we tested the sequence 222T, which forms different G-quadruplex conformers depending on the K^+^ concentration.^24^ First, we observed clearly different exchange profiles for 222T•2K^+^, a three-tetrad quadruplex, in 0.1 or 1.0 mM KCl solutions (Figure 5B). Second, such differences can be easily measured for different species in equilibrium by operating in native MS conditions. Here, a two-tetrad quadruplex 222T•1K^+^ and an unfolded conformer 222T•0K^+^ are also detected (Figure 5C): 222T•1K^+^ exchanges quicker than 222T•2K^+^, and 222T•0K^+^ exchanges within a second owing to the absence of base pairing (Figure 5B). The result also means that the three complexes do not interconvert faster than the time scale of the experiment. This showcases the advantage of coupling native MS to HDX to characterize several conformers in a mixture, which can be challenging with conventional methods. More generally, all the unfolded oligonucleotides we essayed systematically exchanged fully in less than a second (and likely close to their intrinsic exchange rates owing to the lack of hydrogen bonding). Conversely, structured oligonucleotides were all found to be protected from exchange to an extent that depends on their precise conformations. G-quartets are remarkably protected (min-to-days exchange), in particular those shielded from the solvent, whereas base-pairs are exchanged within seconds to minutes (see ds26 in Figure 4B), likely owing to the breathing fluctuations that increase their exchange availability.^13^ Among G-quadruplexes, significant differences in exchange rates were observed in the examples above (Figures 4 and 5), as well as in a larger panel of G-quadruplex structures, even across structures of similar numbers of protected sites (Figures S42), further demonstrating that the exchange rates are strongly dependent on the precise conformers at hand. This opens promising avenues for the use of native HDX/MS nucleic acids biophysics by native HDX/MS.

## CONCLUSION

We demonstrated that oligonucleotides are perfectly amenable to HDX/native MS, by using DNA G-quadruplexes as models. Most notably, we have established that:

i. Nucleobases are isotopically exchanged proportionally to the solution deuterium content without in-source back-exchange, whereas phosphates are virtually entirely back exchanged, using TMAA as a buffer and relatively soft ionization conditions.
ii. The apparent exchange rates depend strongly on the specific hydrogen bonding status of DNA bases, and therefore on oligonucleotides’ secondary structure dynamics.
iii. Mass spectrometry can precisely measure these exchange rates over a wide time range (seconds to minutes with continuous flow setups, minutes to days with manual mixing), without influence from the charge state or the presence of non-specific adducts.

The technique will find numerous applications in nucleic acids biophysics. The relatively low concentrations of oligonucleotides (here 5 µM) allows avoiding the formation of non-physiologically-relevant multimers.^16^ It does not require digestion, quenching and chromatographic separation steps, found routinely in bottom-up HDX/MS analysis of proteins. Native HDX/MS also does not require expensive or highly specialized equipment other than the mass spectrometer itself. The use of native MS gives concomitant access to other biophysical parameters (*e.g.* binding constants, stoichiometries), opening up promising avenues for the analysis of oligonucleotide structures and interactions. In particular, coupling to ion mobility mass spectrometry to further disentangle complex conformational ensembles and small-molecule ligand binding analysis that cannot be straightforwardly deciphered by NMR or X-ray crystallography, will be presented in further communications. The use of ^1^H-NMR will also be explored as it can potentially give access to fast exchange rates at the atomic level (by e.g. phase-modulated CLEAN chemical exchange (CLEANEX-PM) sequence),^68,69^ which could provide insights complementary to those of native MS.

The method can also find applications in the quality control of structured oligonucleotides, such as aptamers. Thanks to its simple and high precision acquisition of a large number of data points, native HDX/MS of oligonucleotide is well suited for typical quality control procedures such as structure batch-to-batch consistency, stability over time, and stress testing (e.g. from oxidant, high temperature, non-physiological pH). Modified sequences are often used in oligonucleotide therapeutics (modifications on e.g. the nucleobase to increase binding affinities, phosphodiester backbone for resisting nuclease degradation, PEGylation for lowering renal clearance),^70^ probes (e.g. fluorophore coupling or fluorescent base analogues),^71^ PCR primers^72^ and affinity tags^73^ (biotinylation). Another possible application of native HDX/MS is therefore the assessment of the influence of modifications on oligonucleotide structural dynamics.

## Supporting information

Figures S1-S42, Tables S1-S5, pH measurement protocol, UV-melting protocol, theoretical isotopic distribution calculation script

HDX data and summary tables

## ASSOCIATED CONTENT

### SUPPORTING INFORMATION

Figures S1—S42, Tables S1—S5, pH measurement protocol, UV-melting protocol, theoretical isotopic distribution calculation script (PDF)

HDX data and summary tables (Excel)

### Author Contributions

The manuscript was written through contributions of all authors. All authors have given approval to the final version of the manuscript.

## ACKNOWLEDGMENT

This work was supported by a grant from the FR TECSAN. We thank Frédéric Rosu at the structural biophysico-chemistry platform of the Institut Européen de Chimie et Biologie for the access to mass spectrometers and his invaluable support. We gratefully acknowledge Laura Fricot, Anaïs Ferrer, Emile Feugas, and Benjamin Lienard for their technical assistance.

## Notes

#### Summary of Updates

Expanded discussion on in-source back-exchange, HDX-NMR, NUS calculation rationale. Minor corrections.

