## Supplementary material for "Native Hydrogen/Deuterium Exchange Mass Spectrometry of Structured DNA Oligonucleotides": Figures S1-S42, Tables S1-S5, pH measurement protocol, UV-melting protocol, theoretical isotopic distribution calculation script

### Contents

Experimental section ........ S3
Oligonucleotides ........ S3
pH of the exchange buffer ........ S4
UV-melting ........ S5
Tune softness ........ S7
Reference tune ........ S7
Soft tune ........ S9
Denaturing tune ........ S11
Continuous-flow setup ........ S14
Data treatment ........ S15
Results and discussion ........ S17
The exchangeable sites are located on the nucleobases. ........ S17
Implications for converting the *m*/*z* distribution into unexchanged sites equivalent and interpreting NUS values. ........ S38
Calculation of theoretical isotopic distribution of oligonucleotides ........ S38
Exchange rates can be measured precisely on a large time scale, regardless of the charge state and adduct stoichiometry ........ S45
References ........ S60

### Experimental section

#### Oligonucleotides

| Name | Sequence (5' to 3') | Description | Structure (literature) | PDB ID | $T_m$±SD (°C)[a] |
|---|---|---|---|---|---|
| **T12** | $T_{12}$ | Poly-thymine | Unfolded | | |
| **T24** | $T_{24}$ | Poly-thymine | Unfolded | | |
| **23nonG4** | $TG_3ATGCGACA(GA)_2G_2ACG_3$ | Non-folding G-rich | Unfolded[1] | | |
| **24nonG4** | $TG_3ATGCGACA(GA)_2G_2ACG_3A$ | Non-folding G-rich | Unfolded[1] | | |
| **c-kit87up** | $(AG_3)_2CGCTG_3AG_2AG_3$ | c-kit promoter | G4: parallel with a snapback feature[2] | 2O3M | 33.1 ± 1.6 |
| **Bcl-2** | $G_3CGCG_3AG_2A_2T_2G_3CG_3$ | BCL-2 promoter | G4: hybrid 3+1[3] | 2F8U | 40.5 ± 0.5 |
| **VEGF** | $CG_4CG_3C_2T_2G_3CG_4T$ | VEGF promoter | G4: parallel[4] | 2M27 | 49.1 ± 0.7 |
| **N-myc** | $TAG_3CG_3(AG_3)_2A_2$ | N-myc intron | G4: parallel[5] | 2LEE | 55.3 ± 0.3 |
| **26CEB** | $A_2G_3TG_3TGTA_2GTGTG_3TG_3T$ | 25CEB human minisatellite locus | G4: parallel[6] | 2LPW | 51.5 ± 0.5 |
| **T30177-TT** | $T_2GTG_2(TG_3)_3T$ | Aptamer (HIV-1 integrase inhibitor) | G4: parallel, with bulge[7] | 2M4P | 55.3 ± 0.3 |
| **T95-2T** | $T_2(G_3T)_4$ | Aptamer (HIV-1 integrase inhibitor) | G4: parallel[8] | 2LK7 | 66.9 ± 0.5 |
| **TBA** | $G_2T_2G_2TGTG_2T_2G_2$ | Aptamer (thrombin) | G4: antiparallel, 2-quartets[9] | 148D | 35.6 ± 0.1 |
| **26TTA** | $(T_2AG_3)_4T_2$ | Human telomere | G4: hybrid 3+1[10] | 2JPZ | 28.4 ± 0.4 |
| **24TTG** | $TTG_3(TTAG_3)_3A$ | Modified human telomere | G4: hybrid 3+1[11] | 2GKU | 40.3 ± 0.4 |
| **23TAG** | $TAG_3(T_2AG_3)_3$ | Human telomere | G4: hybrid 3+1[12] | 2JSM | 37 ± 0.4 |
| **22GT** | $(G_3T_2A)_3G_3T$ | Human telomere | G4: two-tetrad basket antiparallel[13] | 2KF8 | 40.7 ± 0.3 |
| **22CTA** | $AG_3(CTAG_3)_3$ | Modified human telomere | G4: antiparallel, 2-quartets, 1 triad[14] | 2KM3 | 36.8 ± 1.1 |
| **TG4T** | $TG_4T$ | Synthetic construct | G4: tetramolecular parallel[15] | 2O4F | |
| **AG4T** | $AG_4T$ | Synthetic construct | G4: tetramolecular parallel | | |
| **TG5T** | $TG_5T$ | Synthetic construct | G4: tetramolecular parallel[16] | | |
| **222T** | $T(G_3T)_4$ | Synthetic construct | G4: parallel[17] | | 41.8 ± 0.3 |
| **ds26** | $CA_2TCG_2ATCGA_2T_2CGATC_2GAT_2G$ | Palindromic sequence | dsDNA | | 69.6 ± 0.7 |

[a]Average and standard deviation on a minimum of 3 independent $T_m$ determinations.

*Table S1. Oligonucleotides used in this study.*

#### pH of the exchange buffer

pH measurements were performed at 22°C with a WTW inoLab Multi 9420 IDS pH-meter.

The calibration was performed daily on 3 points, and accepted for calibration slopes within 98—102% of the theoretical value, i.e. -58.2 mV $pH^{-1}$ at 20°C, and a pH 7.0 offset within 6 mV.

The pH of a given buffer batch was very consistent on different days of a week (pH = 7.04 ± 0.005, $n$ = 4 with independent calibrations) and remained stable over several months at 4°C ($\Delta$pH < 0.1 over 9 months).

The pH was also consistent across different independent preparations (pH = 7.04 ± 0.004, $n$ = 4 measured with independent calibrations on different days).

#### UV-melting

The samples contained 10 µM DNA (from 1 mM stock solutions) in 1 mM KCl/100 mM TMAA. They were annealed at 85°C for 5 min, and stored at 4°C for at least a night before use.

Melting temperatures were determined by measuring the changes in absorbance as a function of the temperature, using a Secomam Uvikon XL spectrophotometer equipped with two thermostatable 6-cell holders and a Julabo F25 bath. Temperature control and programming were performed using the Julabo EsayTemp professional and Thermalys (Secomam) software.

The samples were first heated to a target temperature of 96.5°C for 8 minutes, and then the absorbance was monitored at 260, 295, and 350 nm on a cycle composed of a cooling down to 4°C at a rate of 0.2°C $min^{-1}$, followed by 15 minutes at 4°C, heating back up to 96.5°C at 0.2°C $min^{-1}$, and a final cooling down to 25°C in one hour.

For G4- and ds-DNA-forming sequences, the absorbance data points at 295 nm and 260 nm, respectively, were used. The blank absorbance at the same wavelengths and the sample absorbance at 335 nm were subtracted from the raw data.[18,19] UV-melting data were analyzed by nonlinear fitting of the corrected absorbance data, based on Equation S1, where $A_T^F$ and $A_T^U$ are respectively the contributions of the folded (F) and unfolded (U) forms to the absorbance (called associated and dissociated baselines in the framework of Mergny and Lacroix[18]), and $\theta_T$ is the folded fraction of oligonucleotide.

$$A_T = A_T^F \times \theta_T + A_T^U \times (1 - \theta_T) \qquad \text{Equation S1}$$

The baselines are given by:

$$A_T^F = (a_F T + b_F) \qquad \text{Equation S2}$$

$$A_T^U = (a_U T + b_U) \qquad \text{Equation S3}$$

With *a* and *b* the slopes and intercepts of the baselines. For an equilibrium between a folded and unfolded form, characterized by an equilibrium constant *K*, the folded fraction can be defined by Equation S4, assuming that the heat capacity does not significantly change between the two forms.[20]

$$\theta = \frac{[F]}{[F]+[U]} = \frac{1}{1+K} = \frac{1}{1+exp\left(-\frac{\Delta H_{Tm}^0\left(1-\frac{T}{Tm}\right)}{RT}\right)} \qquad \text{Equation S4}$$

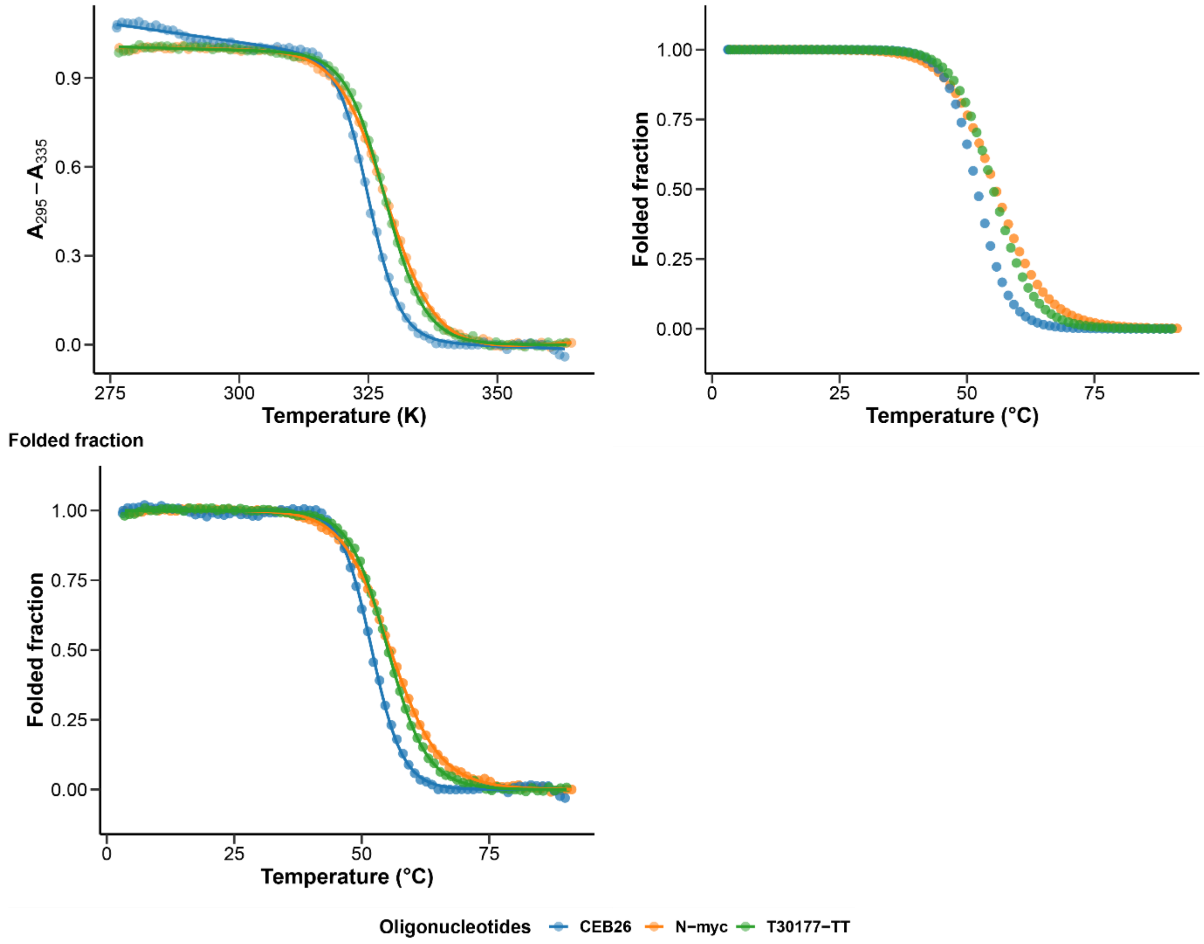


*Figure S1. UV-melting of 26CEB, N-myc, and T30177-TT: raw data fitted with Equation S4 (top left), data modeled from the fitting results (top right), and folded fraction (bottom left).*

#### Tune softness

##### Reference tune

=== Tune Data: ===

| | |
|---|---|
| Spray Voltage (+) | 3500.00 |
| Spray Voltage (-) | 2950.00 |
| Capillary Temperature (+ or +-) | 165.00 |
| Capillary Temperature (-) | 200.00 |
| Sheath Gas (+ or +-) | 60.00 |
| Sheath Gas (-) | 55.00 |
| Aux Gas (+ or +-) | 2.00 |
| Aux Gas (-) | 5.00 |
| Spare Gas (+ or +-) | 0.00 |
| Spare Gas (-) | 0.00 |
| Max Spray Current (+) | 100.00 |
| Max Spray Current (-) | 100.00 |
| Heater Temperature (+ or +-) | 0.00 |
| Heater Temperature (-) | 25.00 |

Thermo Exactive Orbitrap Data:

==============================

| | |
|---|---|
| AGC | On |
| Micro Scan Count | 2 |
| Scan Segment | 0 |
| Scan Event | 92 |
| Ion Injection Time (ms) | 250.000 |
| Max. Ion Time (ms) | 250.00 |
| FT Resolution | 50000 |
| AGC Target | 3000000 |
| Analyzer Temperature | 29.97 |

=== Mass Calibration: ===

| | |
|---|---|
| Conversion Parameter B | 67833113.0979 |
| Conversion Parameter C | -24702974.7587 |
| Temperature Comp. (ppm) | -4.50 |
| RF Comp. (ppm) | 0.29 |
| Space Charge Comp. (ppm) | -0.05 |
| Resolution Comp. (ppm) | 0.27 |
| Number of Lock Masses | 0 |
| Lock Mass #1 (m/z) | 0.0000 |
| Lock Mass #2 (m/z) | 0.0000 |
| Lock Mass #3 (m/z) | 0.0000 |
| LM Search Window (ppm) | 0.0 |
| Number of LM Found | 0 |
| Last Locking (sec) | 0.0 |
| LM Correction (ppm) | 0.00 |

=== Ion Optics Settings: ===

| | |
|---|---|
| Capillary Voltage (V) | -15.00 |
| Tube lens Voltage (V) | -200.00 |
| Skimmer Voltage (V) | -14.00 |

| | |
|---|---|
| RF0 DC Offset (V) | -6.00 |
| Lens 1 Voltage (V) | 0.00 |
| MP0 and MP1 RF (V) | 700.00 |
| RF1 DC Offset (V) | -7.00 |
| MP2 and MP3 RF (V) | 700.00 |
| Gate lens Voltage (V) | -7.10 |
| C-trap RF (V) | 2400.0 |
| ==== Extra Settings: ==== | |
| HCD stepped energy | 0.00 |
| ==== Diagnostic Data: ==== | |
| Intens Comp Factor | 0.5785 |
| PS Inj. Time (ms) | 125.000 |
| AGC PS Mode 1 | |
| Analog Input 1 (V) | -0.002 |
| Analog Input 2 (V) | 0.001 |

#### Soft tune

| | |
|---|---|
| === Tune Data: === | |
| Spray Voltage (+) | 3500.00 |
| Spray Voltage (-) | 2950.00 |
| Capillary Temperature (+ or +-) | 165.00 |
| Capillary Temperature (-) | 170.00 |
| Sheath Gas (+ or +-) | 60.00 |
| Sheath Gas (-) | 55.00 |
| Aux Gas (+ or +-) | 2.00 |
| Aux Gas (-) | 5.00 |
| Spare Gas (+ or +-) | 0.00 |
| Spare Gas (-) | 0.00 |
| Max Spray Current (+) | 100.00 |
| Max Spray Current (-) | 100.00 |
| Heater Temperature (+ or +-) | 0.00 |
| Heater Temperature (-) | 25.00 |
| Thermo Exactive Orbitrap Data: | |
| ============================== | |
| AGC | On |
| Micro Scan Count | 2 |
| Scan Segment | 0 |
| Scan Event | 92 |
| Ion Injection Time (ms) | 250.000 |
| Max. Ion Time (ms) | 250.00 |
| FT Resolution | 50000 |
| AGC Target | 3000000 |
| Analyzer Temperature | 29.92 |
| === Mass Calibration: === | |
| Conversion Parameter B | 67833643.6361 |
| Conversion Parameter C | -23070061.7380 |
| Temperature Comp. (ppm) | -4.45 |
| RF Comp. (ppm) | 0.24 |
| Space Charge Comp. (ppm) | -0.15 |
| Resolution Comp. (ppm) | -0.03 |
| Number of Lock Masses | 0 |
| Lock Mass #1 (m/z) | 0.0000 |
| Lock Mass #2 (m/z) | 0.0000 |
| Lock Mass #3 (m/z) | 0.0000 |
| LM Search Window (ppm) | 0.0 |
| Number of LM Found | 0 |
| Last Locking (sec) | 0.0 |
| LM Correction (ppm) | 0.00 |
| === Ion Optics Settings: === | |
| Capillary Voltage (V) | -20.00 |
| Tube lens Voltage (V) | -192.00 |
| Skimmer Voltage (V) | -15.00 |
| RF0 DC Offset (V) | -6.00 |
| Lens 1 Voltage (V) | 0.00 |

| | |
|---|---|
| MP0 and MP1 RF (V) | 700.00 |
| RF1 DC Offset (V) | -7.00 |
| MP2 and MP3 RF (V) | 700.00 |
| Gate lens Voltage (V) | -7.10 |
| C-trap RF (V) | 2400.0 |
| ==== Extra Settings: ==== | |
| HCD stepped energy | 0.00 |
| ==== Diagnostic Data: ==== | |
| Intens Comp Factor | 0.5785 |
| PS Inj. Time (ms) | 125.000 |
| AGC PS Mode 1 | |
| Analog Input 1 (V) | -0.002 |
| Analog Input 2 (V) | 0.001 |

#### Denaturing tune

=== Tune Data: ===

| | |
|---|---|
| Spray Voltage (+) | 3500.00 |
| Spray Voltage (-) | 2950.00 |
| Capillary Temperature (+ or +-) | 165.00 |
| Capillary Temperature (-) | 240.00 |
| Sheath Gas (+ or +-) | 60.00 |
| Sheath Gas (-) | 55.00 |
| Aux Gas (+ or +-) | 2.00 |
| Aux Gas (-) | 5.00 |
| Spare Gas (+ or +-) | 0.00 |
| Spare Gas (-) | 0.00 |
| Max Spray Current (+) | 100.00 |
| Max Spray Current (-) | 100.00 |
| Heater Temperature (+ or +-) | 0.00 |
| Heater Temperature (-) | 25.00 |
| Thermo Exactive Orbitrap Data: | |
| ============================== | |
| AGC | On |
| Micro Scan Count | 2 |
| Scan Segment | 0 |
| Scan Event | 92 |
| Ion Injection Time (ms) | 250.000 |
| Max. Ion Time (ms) | 250.00 |
| FT Resolution | 50000 |
| AGC Target | 3000000 |
| Analyzer Temperature | 31.37 |
| === Mass Calibration: === | |
| Conversion Parameter B | 67834646.2145 |
| Conversion Parameter C | -24957445.1995 |
| Temperature Comp. (ppm) | -5.76 |
| RF Comp. (ppm) | 0.00 |
| Space Charge Comp. (ppm) | -0.11 |
| Resolution Comp. (ppm) | -1.23 |
| Number of Lock Masses | 0 |
| Lock Mass #1 (m/z) | 0.0000 |
| Lock Mass #2 (m/z) | 0.0000 |
| Lock Mass #3 (m/z) | 0.0000 |
| LM Search Window (ppm) | 0.0 |
| Number of LM Found | 0 |
| Last Locking (sec) | 0.0 |
| LM Correction (ppm) | 0.00 |
| === Ion Optics Settings: === | |
| Capillary Voltage (V) | -15.00 |
| Tube lens Voltage (V) | -200.00 |
| Skimmer Voltage (V) | -14.00 |
| RF0 DC Offset (V) | -6.00 |
| Lens 1 Voltage (V) | 0.00 |

| | |
|---|---|
| MP0 and MP1 RF (V) | 700.00 |
| RF1 DC Offset (V) | -7.00 |
| MP2 and MP3 RF (V) | 700.00 |
| Gate lens Voltage (V) | -7.10 |
| C-trap RF (V) | 2400.0 |
| ==== Extra Settings: ==== | |
| HCD stepped energy | 0.00 |
| ==== Diagnostic Data: ==== | |
| Intens Comp Factor | 0.6964 |
| PS Inj. Time (ms) | 125.000 |
| AGC PS Mode | 1 |
| Analog Input 1 (V) | -0.007 |
| Analog Input 2 (V) | 0.000 |

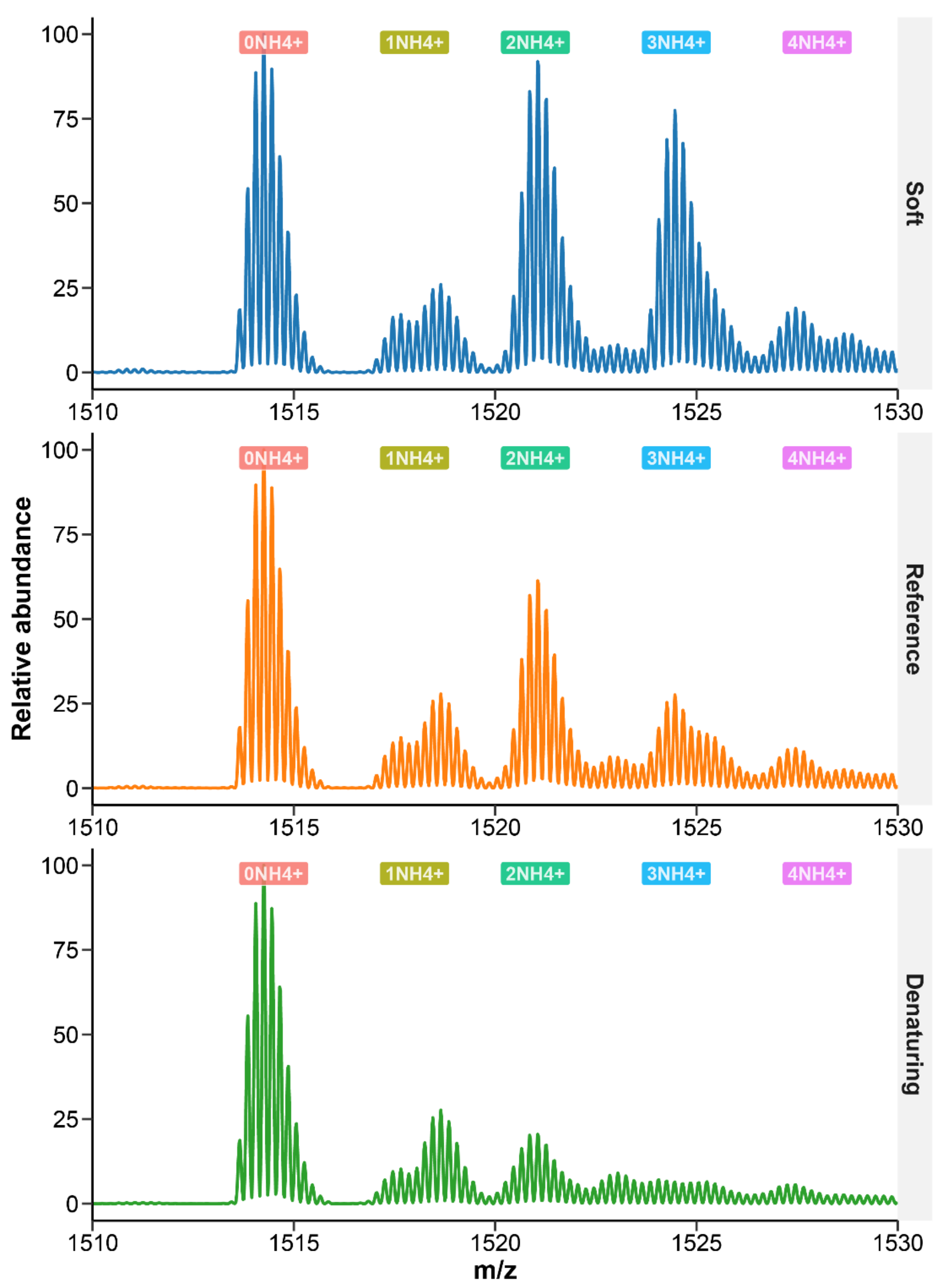


*Figure S2. Mass spectra of a 10-µM solution of $[G_4T_4G_4]_2$ G-quadruplex in 100 mM ammonium acetate, zoomed on the 5- charge state, using three tunes of varying softness. Decreasing the softness of the tunes yields smaller proportions of the fragile 3-ammonium binding quadruplex.*

#### Continuous-flow setup

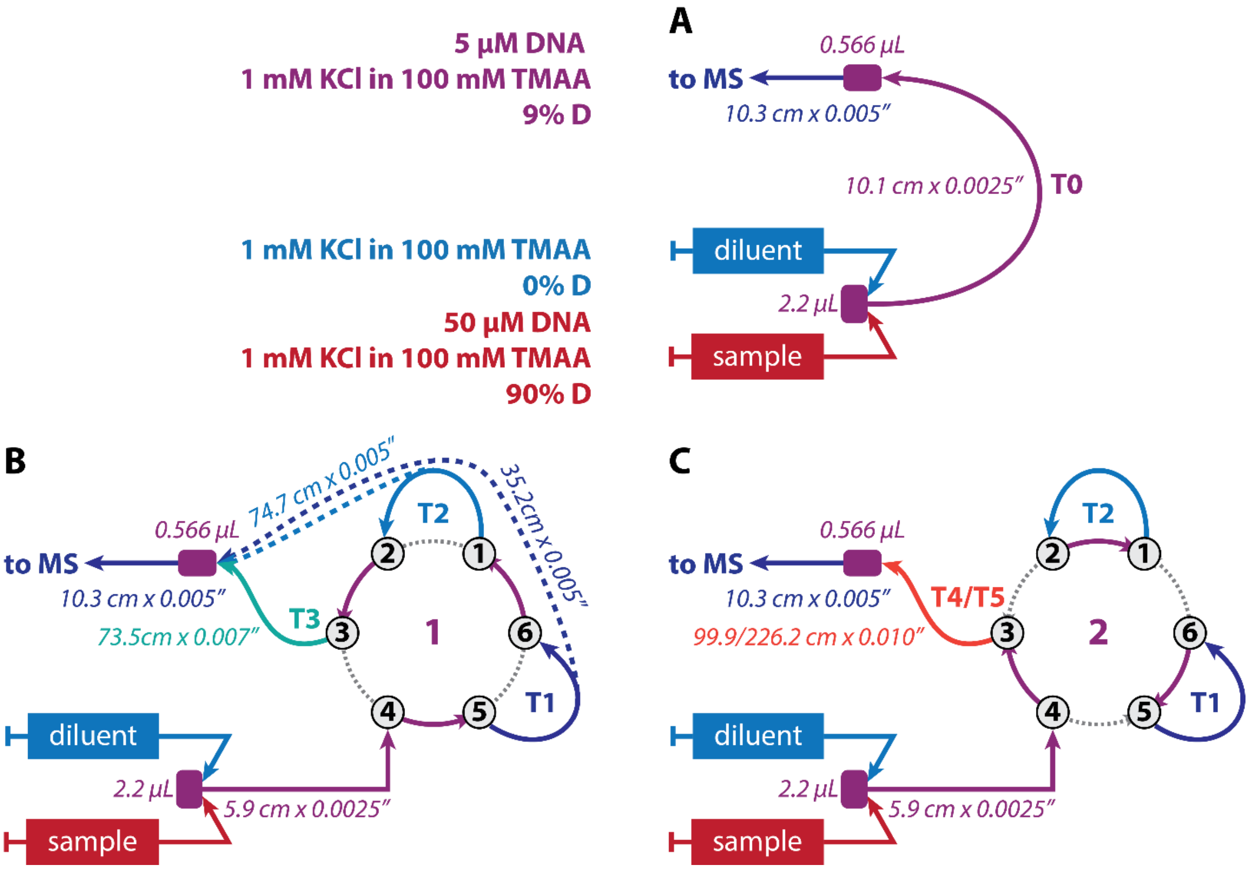


*Figure S3. Continuous-flow setup allowing a variety of mixing times. Configuration A bypasses the valve and gives access to the shortest mixing times (< 10s). Configuration B involves 1 to 3 tubes of increasing volumes linked in series in the position 1 of the valve to reach times in the 10 s—75 s. In configuration C, the valve is switched to either position to reach mixing times up to 5 minutes.*

#### Data treatment

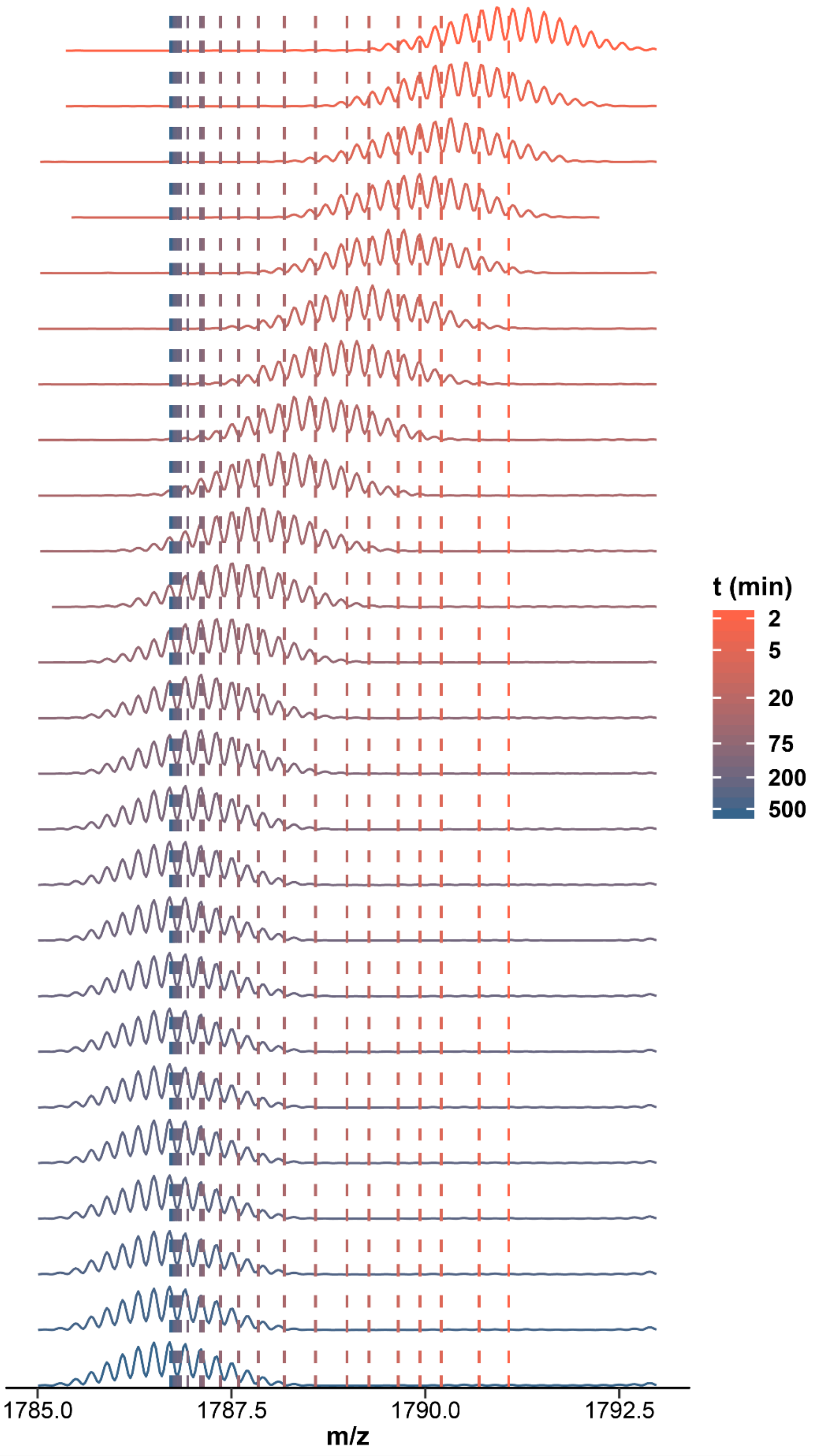


*Figure S4. HDX/native MS spectra of [TG$_5$T]$_4$, zoomed on the 5- charge state, 4-K$^+$ species. 25 exchange time points are presented from top to bottom. The centroids of each time points are shown with a dashed line of the corresponding color. The exchange occurs from D to H, that is from high m/z to low m/z*

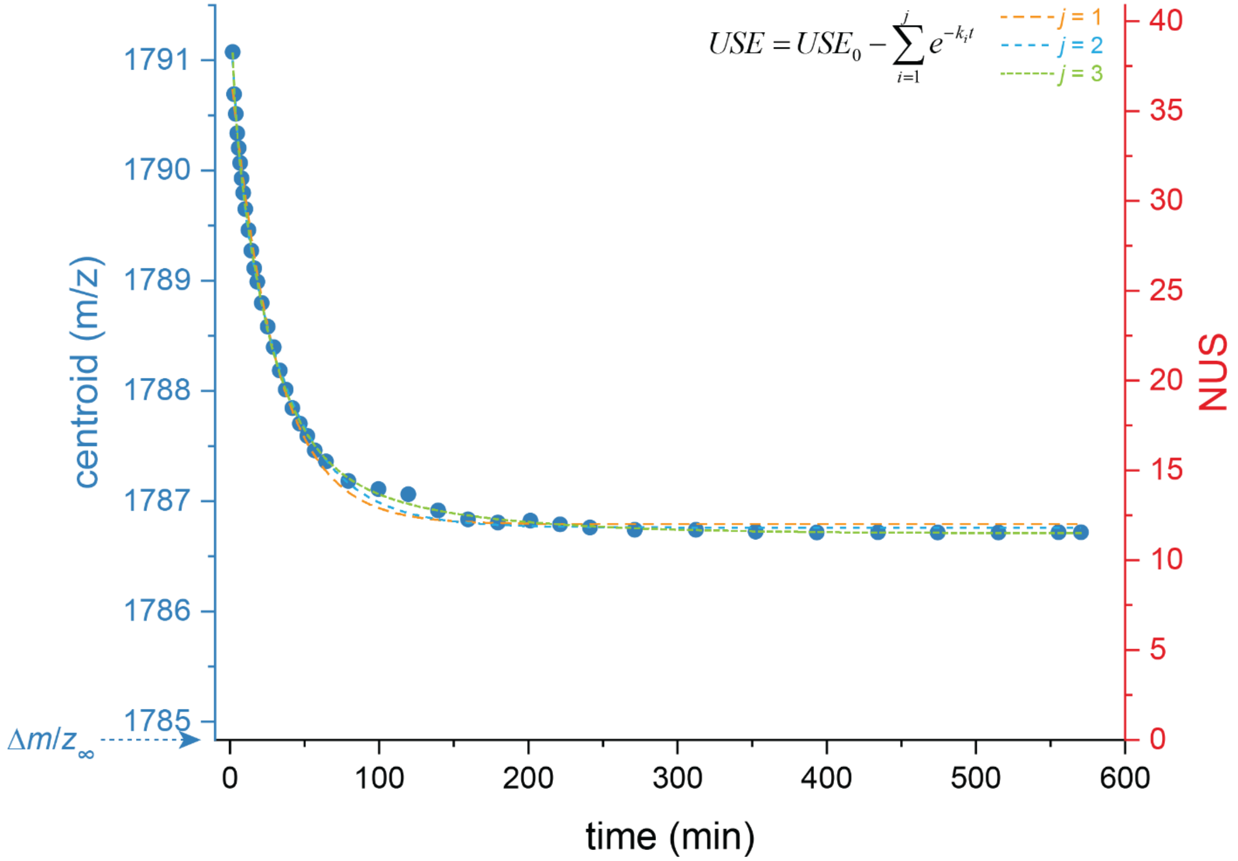


*Figure S5. Exchange plot expressed as the centroid (in m/z units; blue, left y-axis) or the NUS (red, right y-axis) against time, of $[TG_5T]_4$ acquired by continuous-flow and manual mixing experiments. NUS = f(t) was fitted with equation 4 with j = 1, 2, or 3.*

#### Results and discussion

##### The exchangeable sites are located on the nucleobases.

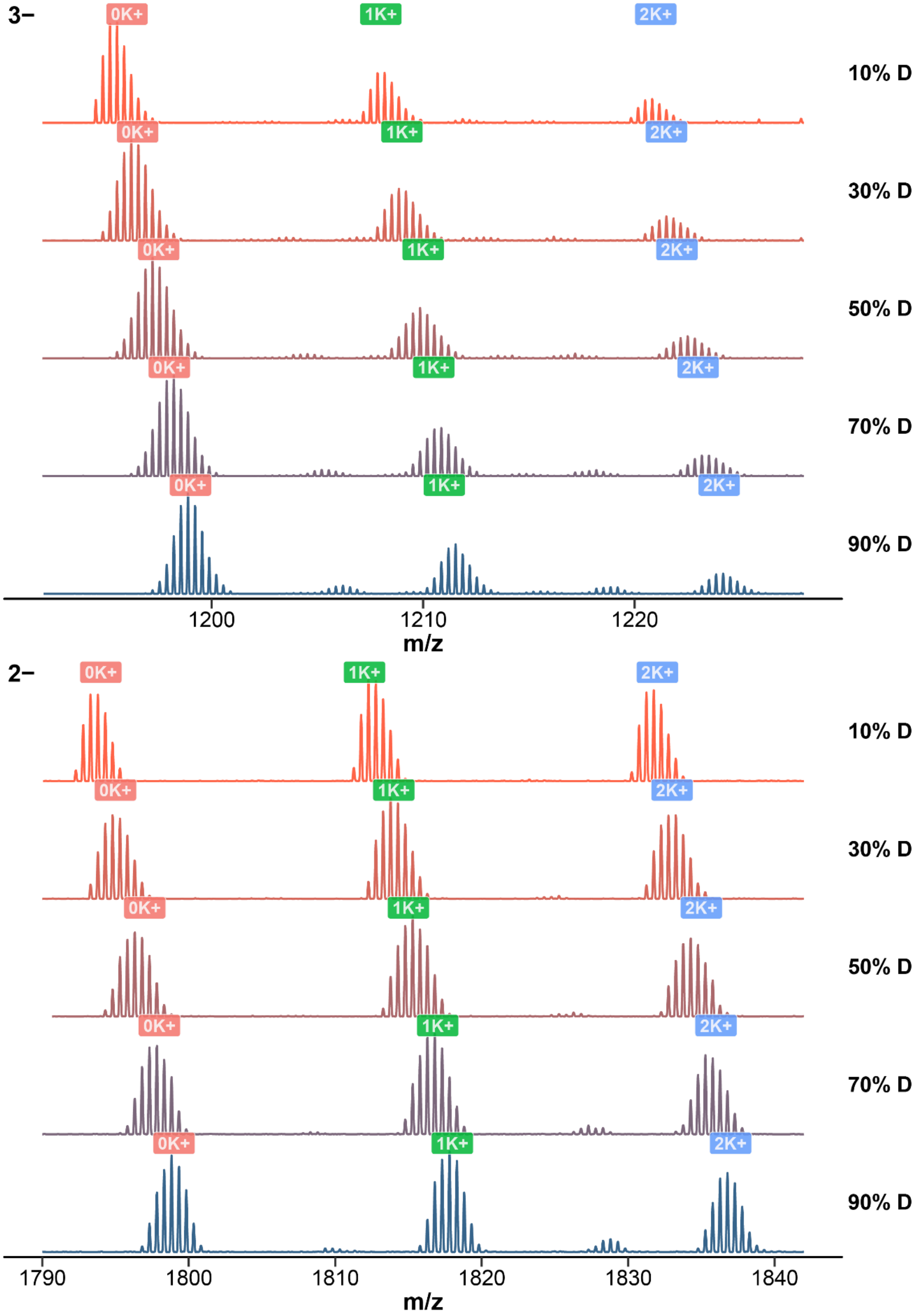


*Figure S6. Native mass spectra of T12 zoomed at the 3- (top) and 2- (bottom) charge states, for solutions containing 10 to 90% deuterium in 100 mM TMAA, 1 mM KCl, using the reference tune. The number of bound potassium is labeled over the peaks.*

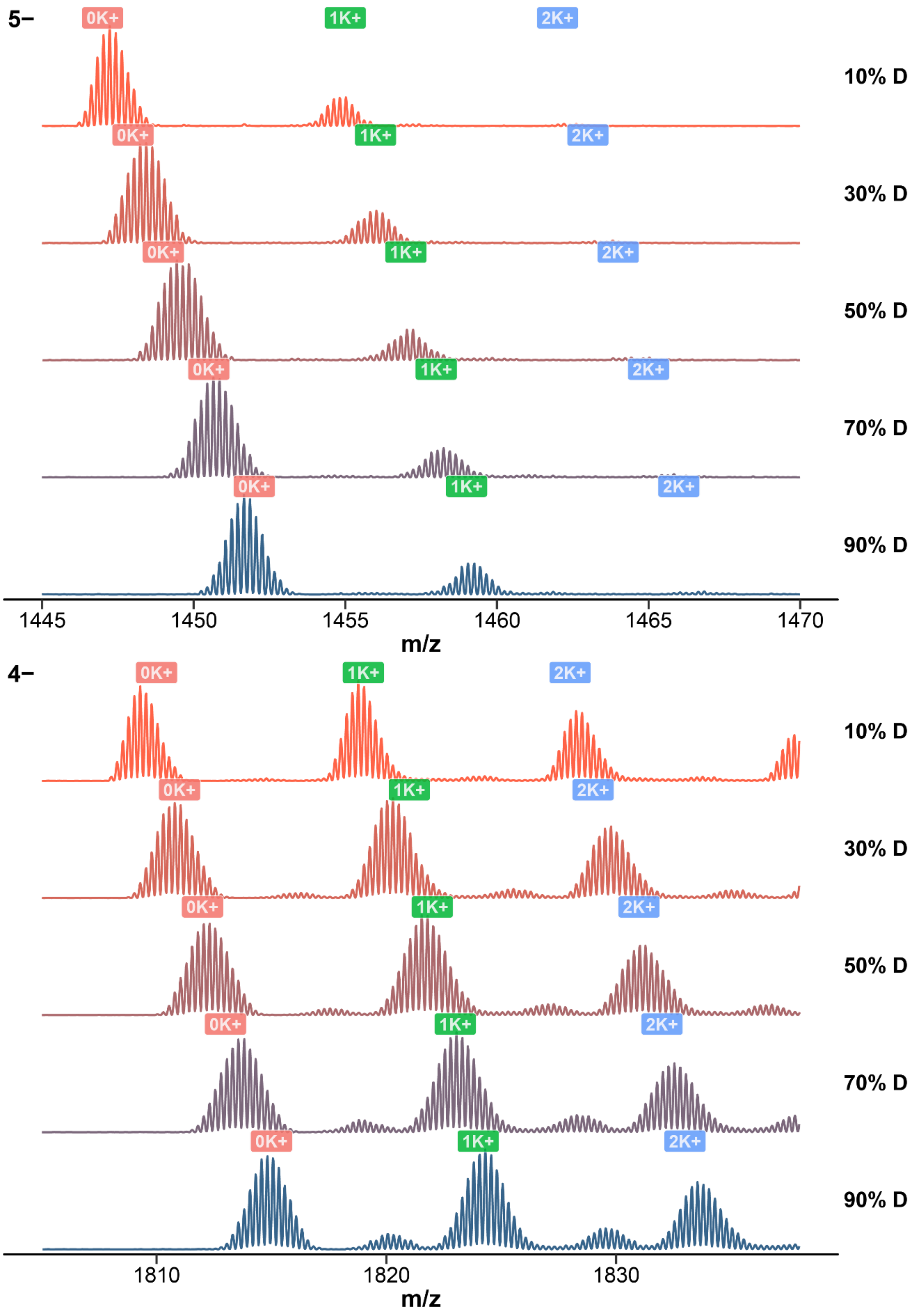


*Figure S7. Native mass spectra of T24 zoomed at the 5- (top) and 4- (bottom) charge states, for solutions containing 10 to 90% deuterium in 100 mM TMAA, 1 mM KCl, using the reference tune. The number of bound potassium is labeled over the peaks.*

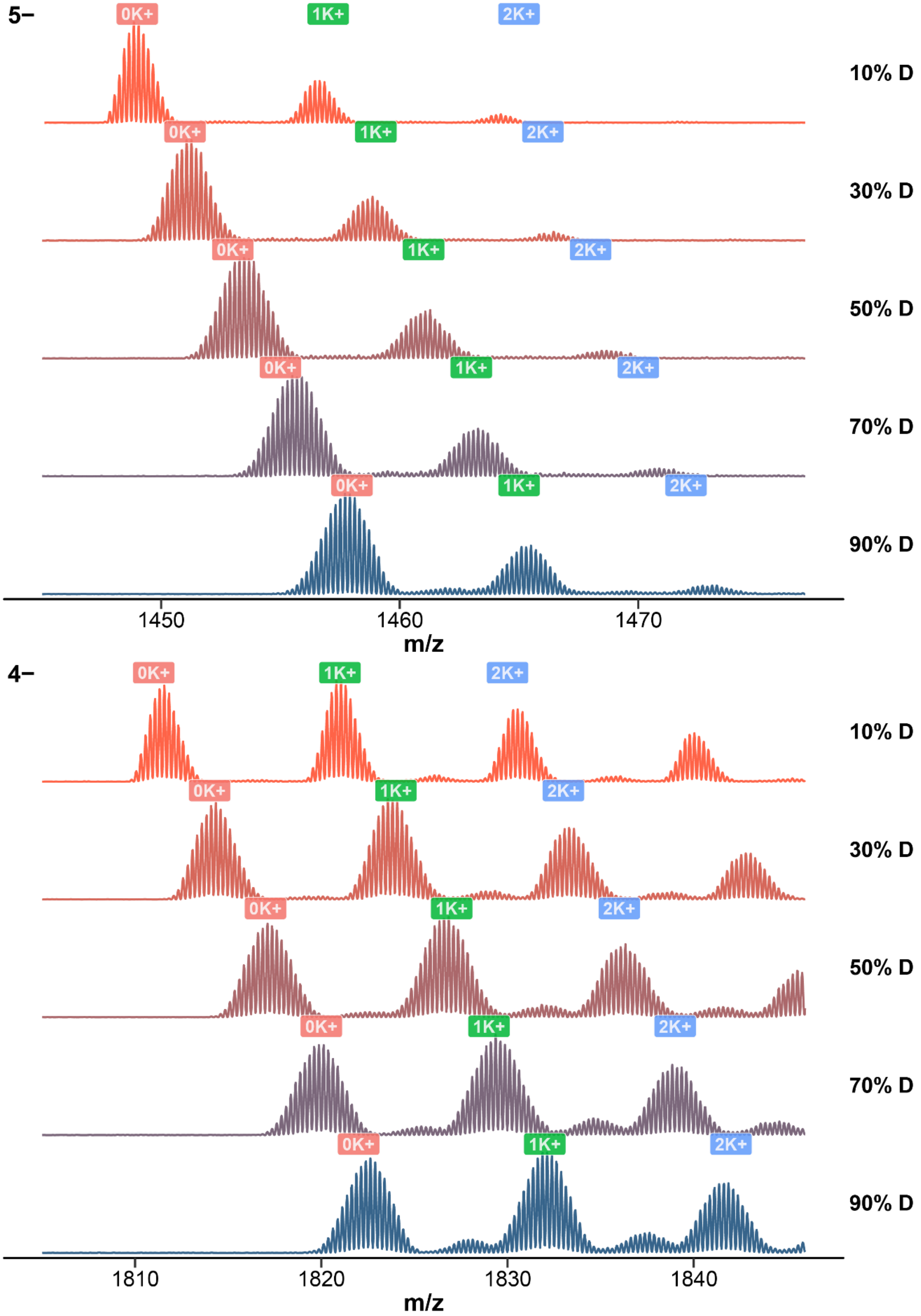


*Figure S8. Native mass spectra of 23nonG4 zoomed at the 5- (top) and 4- (bottom) charge states, for solutions containing 10 to 90% deuterium in 100 mM TMAA, 1 mM KCl, using the reference tune. The number of bound potassium is labeled over the peaks.*

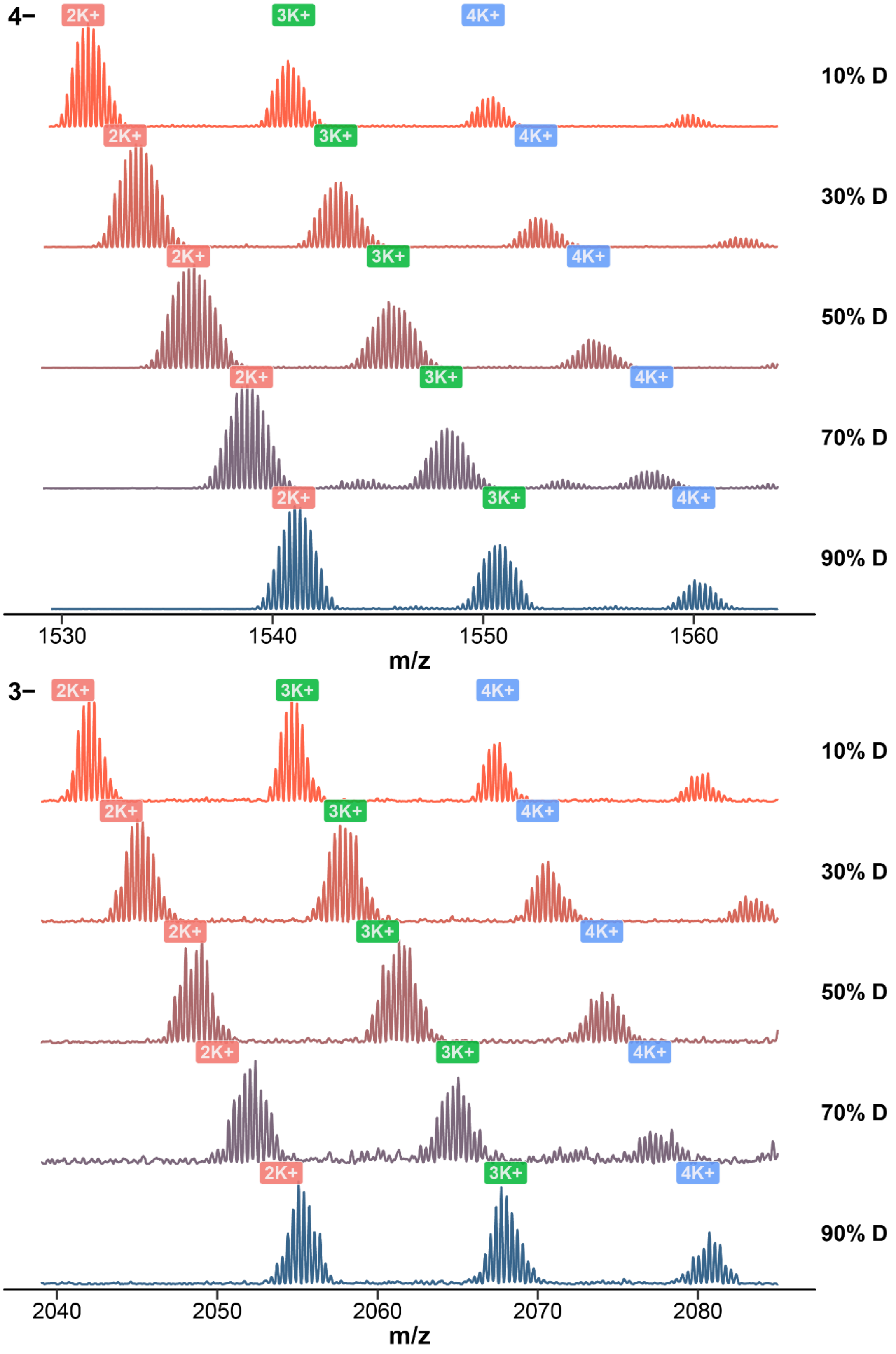


*Figure S9. Native mass spectra of N-myc zoomed at the 4- (top) and 3- (bottom) charge states, for solutions containing 10 to 90% deuterium in 100 mM TMAA, 1 mM KCl, using the reference tune. The number of bound potassium is labeled over the peaks.*

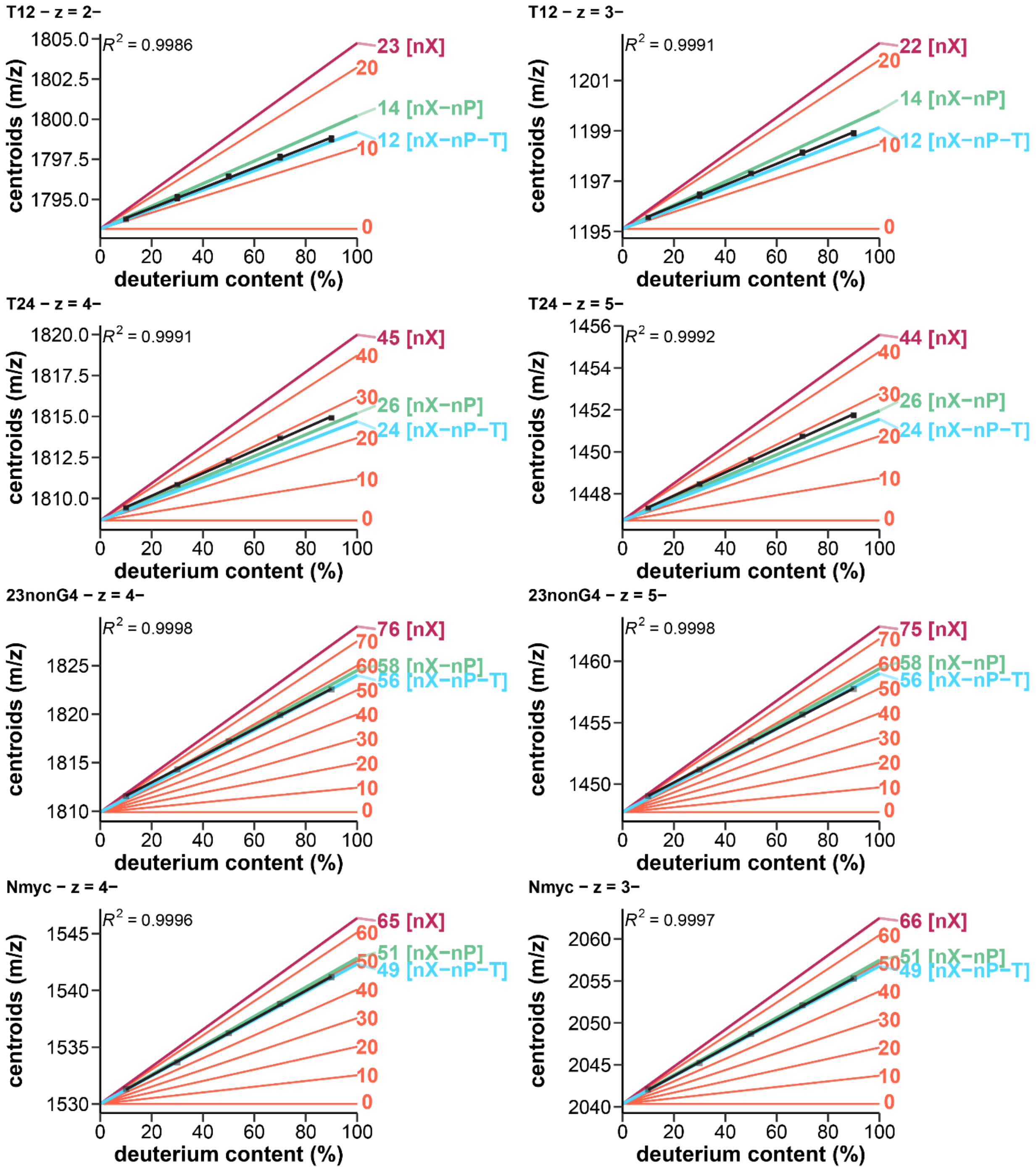


*Figure S10. Exchangeable sites in T12, T24, 23nonG4, and Nmyc for two different charge states, in 100 mM TMAA, 1 mM KCl, using the reference and softer tunes (average values, standard deviation shown as error bars): Measured centroid masses against the solution deuterium content (black square, linear fit: blue line), compared to notable theoretical masses: maximum number of potentially exchangeable sites (nX; purple), no phosphate exchanged (nX–nP; green), no phosphate and termini exchanged (nX–nP–T; cyan), and every ten sites (coral).*

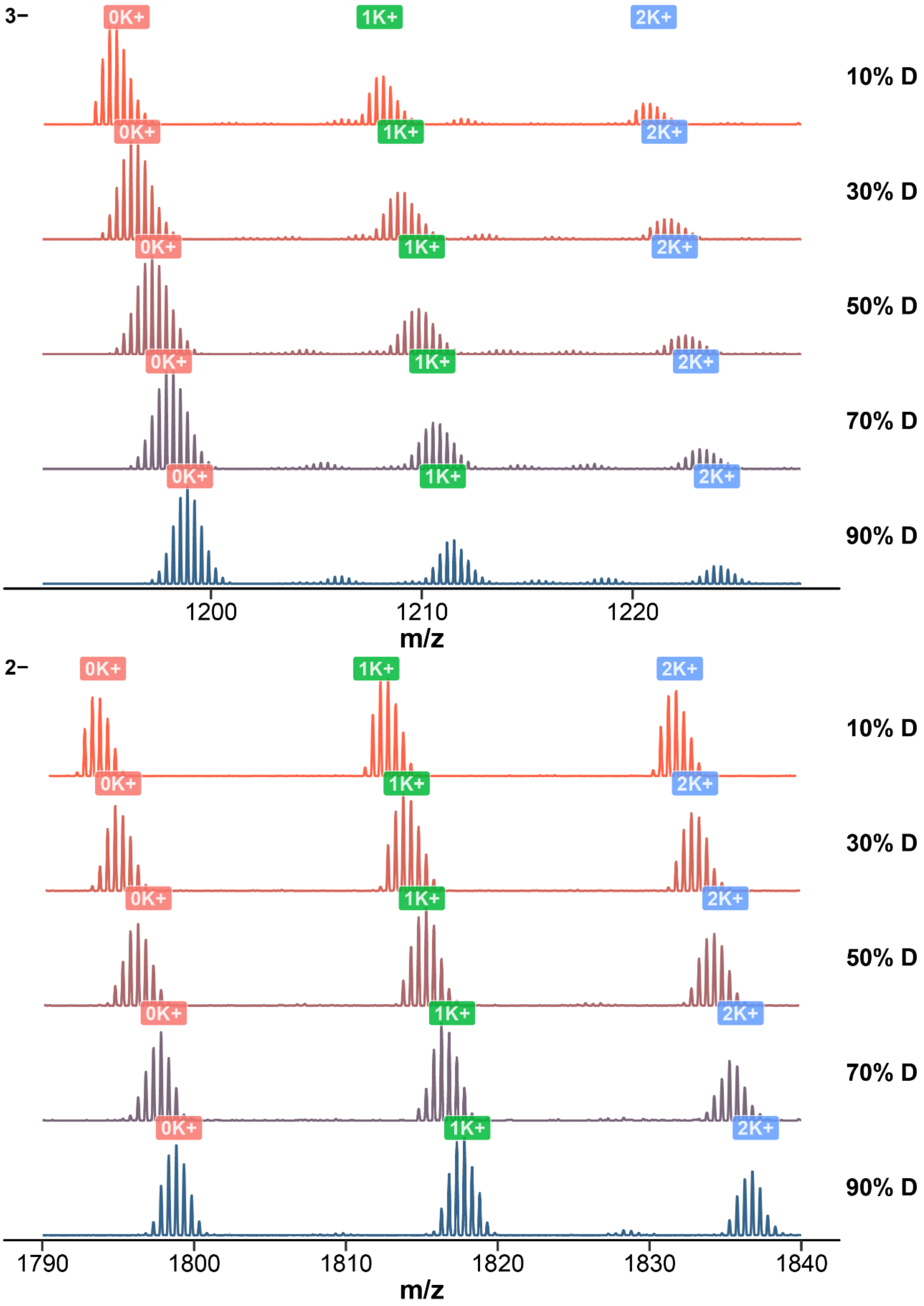


*Figure S11. Native mass spectra of T12 zoomed at the 3- (top) and 2- (bottom) charge states, for solutions containing 10 to 90% deuterium in 100 mM TMAA, 1 mM KCl, infused in an $H_2O$-doped source atmosphere. The number of bound potassium is labeled over the peaks.*

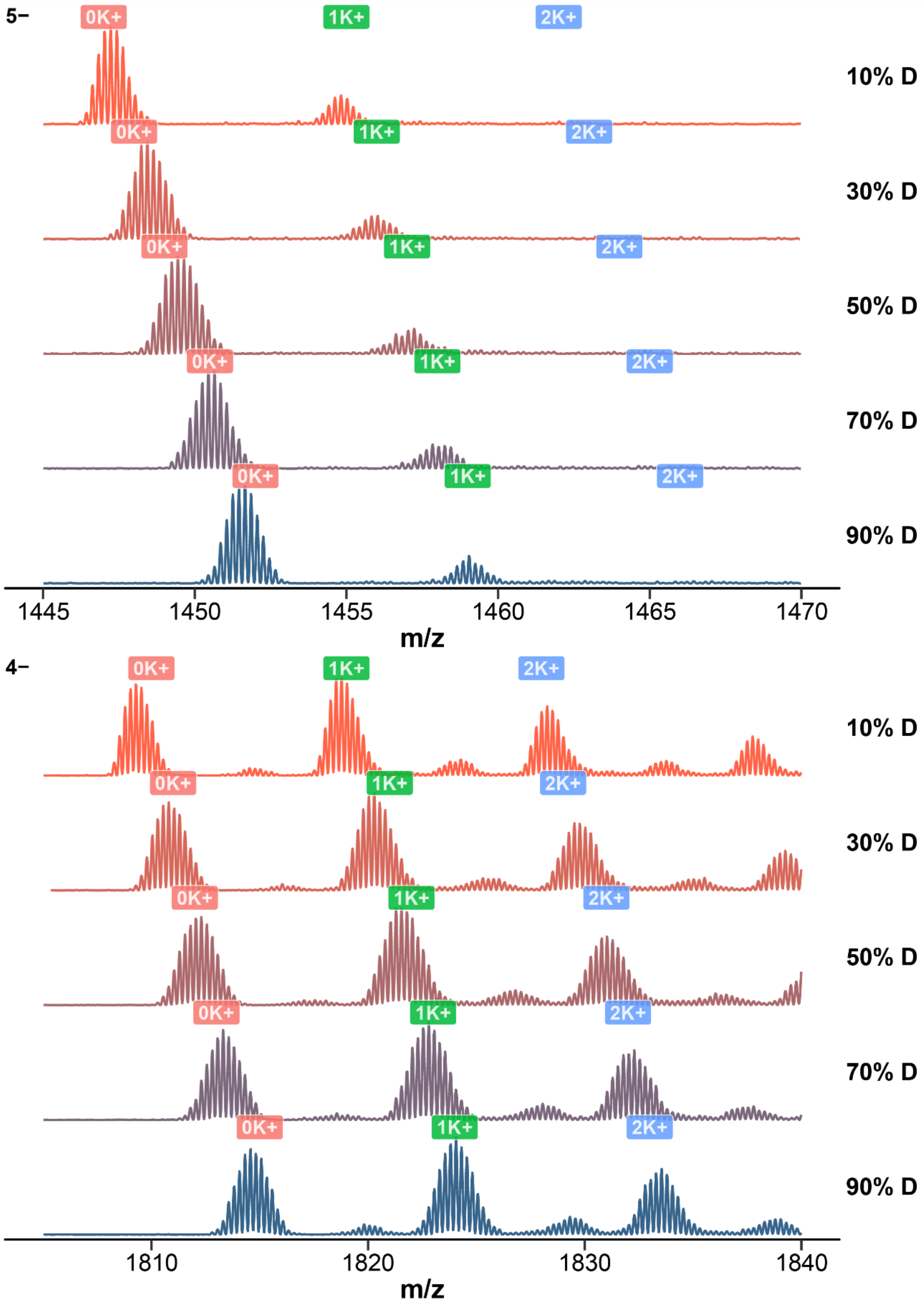


*Figure S12. Native mass spectra of T24 zoomed at the 5- (top) and 4- (bottom) charge states, for solutions containing 10 to 90% deuterium in 100 mM TMAA, 1 mM KCl, infused in an $H_2O$-doped source atmosphere. The number of bound potassium is labeled over the peaks.*

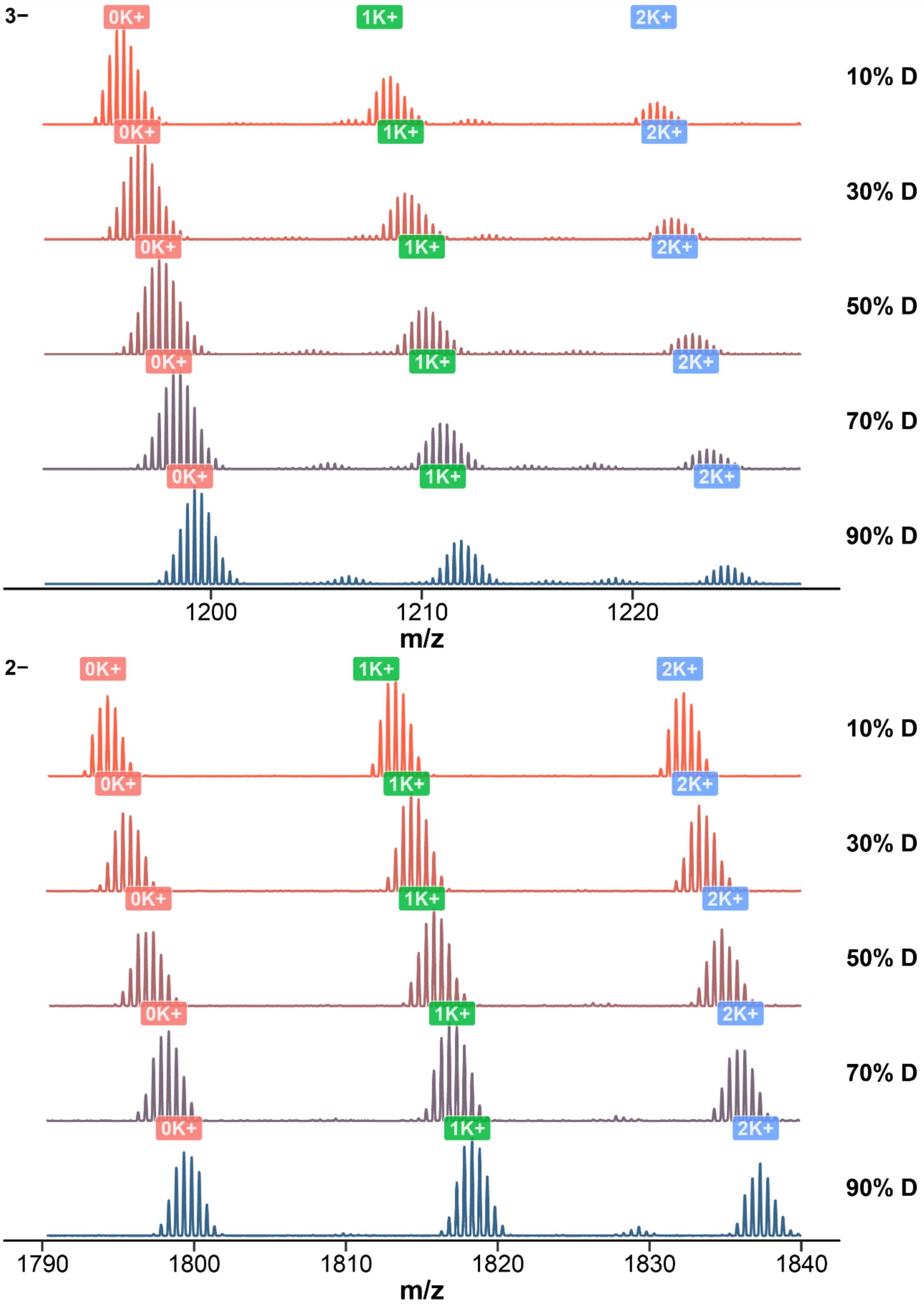


*Figure S13. Native mass spectra of T12 zoomed at the 3- (top) and 2- (bottom) charge states, for solutions containing 10 to 90% deuterium in 100 mM TMAA, 1 mM KCl, infused in a $D_2O$-doped source atmosphere. The number of bound potassium is labeled over the peaks.*

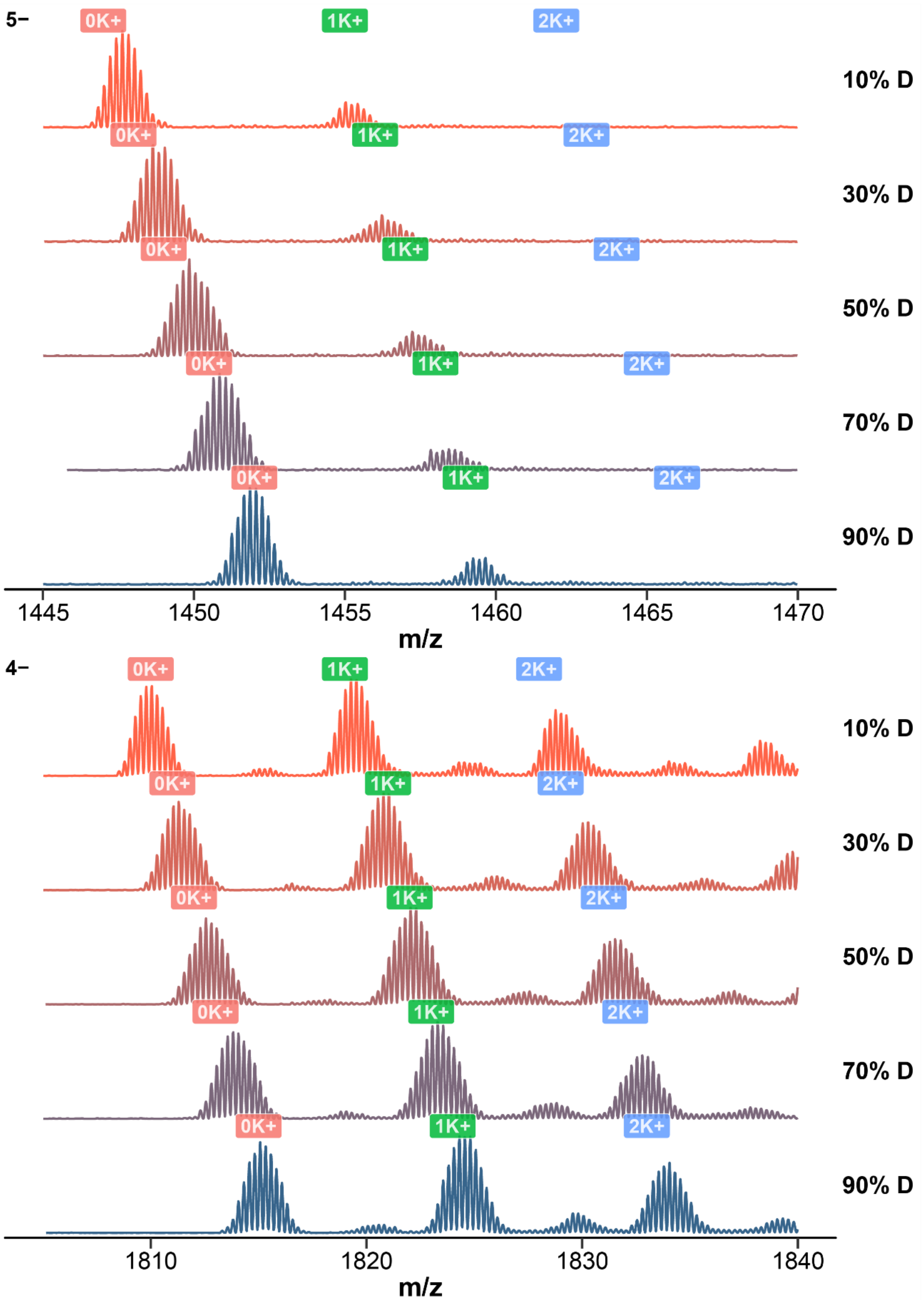


*Figure S14. Native mass spectra of T24 zoomed at the 5- (top) and 4- (bottom) charge states, for solutions containing 10 to 90% deuterium in 100 mM TMAA, 1 mM KCl, infused in an $D_2O$-doped source atmosphere. The number of bound potassium is labeled over the peaks.*

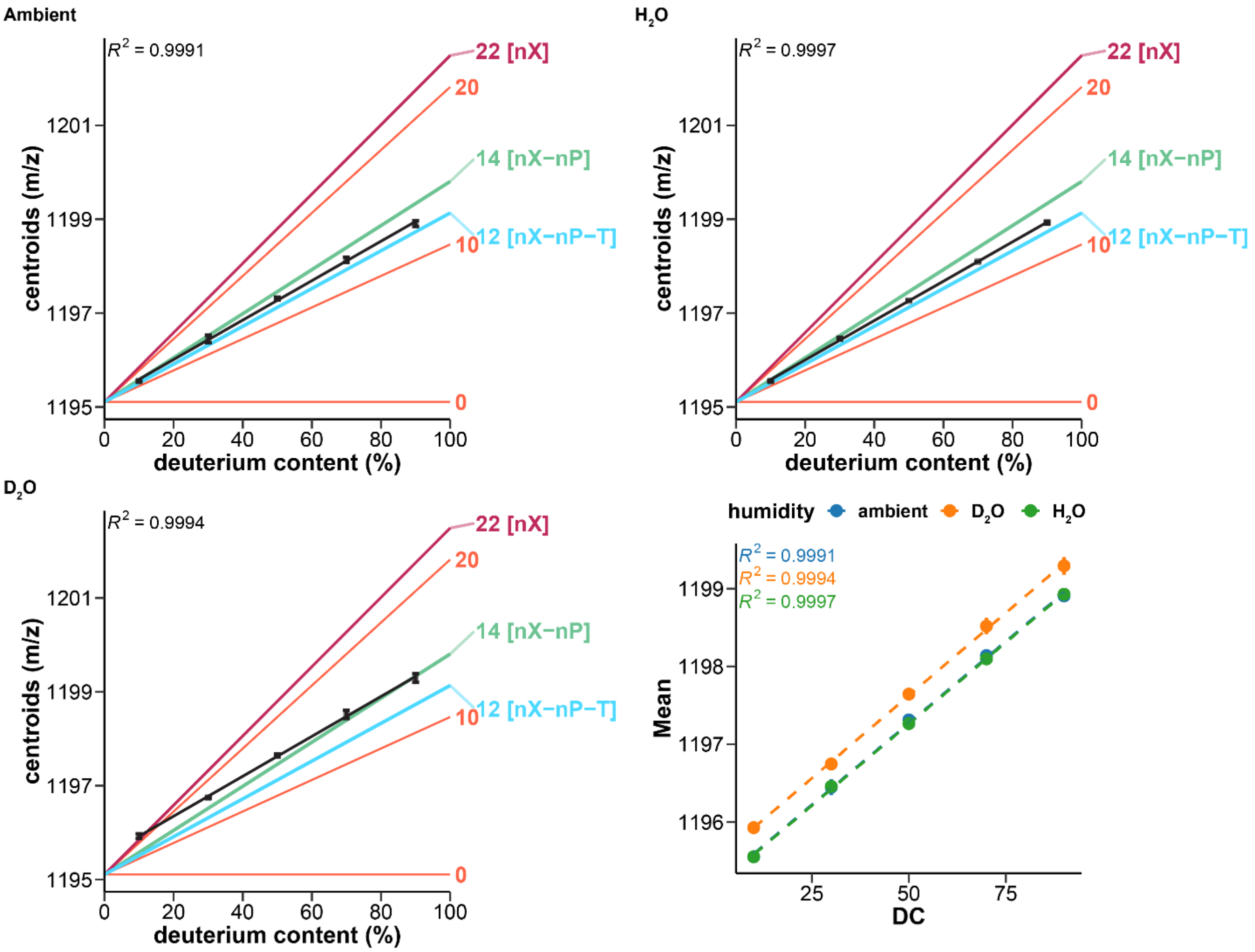


*Figure S15. Exchangeable sites in T12 at z = 3-, in 100 mM TMAA, 1 mM KCl, in different atmosphere (top left: ambient humidity, top right: $H_2O$-doped, bottom left: $D_2O$-doped, bottom right: overlaid data). Measured centroid masses against the solution deuterium content (standard deviation on duplicate measurements as error bars, linear fit as blacklines), compared to notable theoretical masses: maximum number of potentially exchangeable sites (*nX*; purple), no phosphate exchanged (*nX–nP*; green), no phosphate and termini exchanged (*nX–nP–T*; cyan), and every ten sites (coral).*

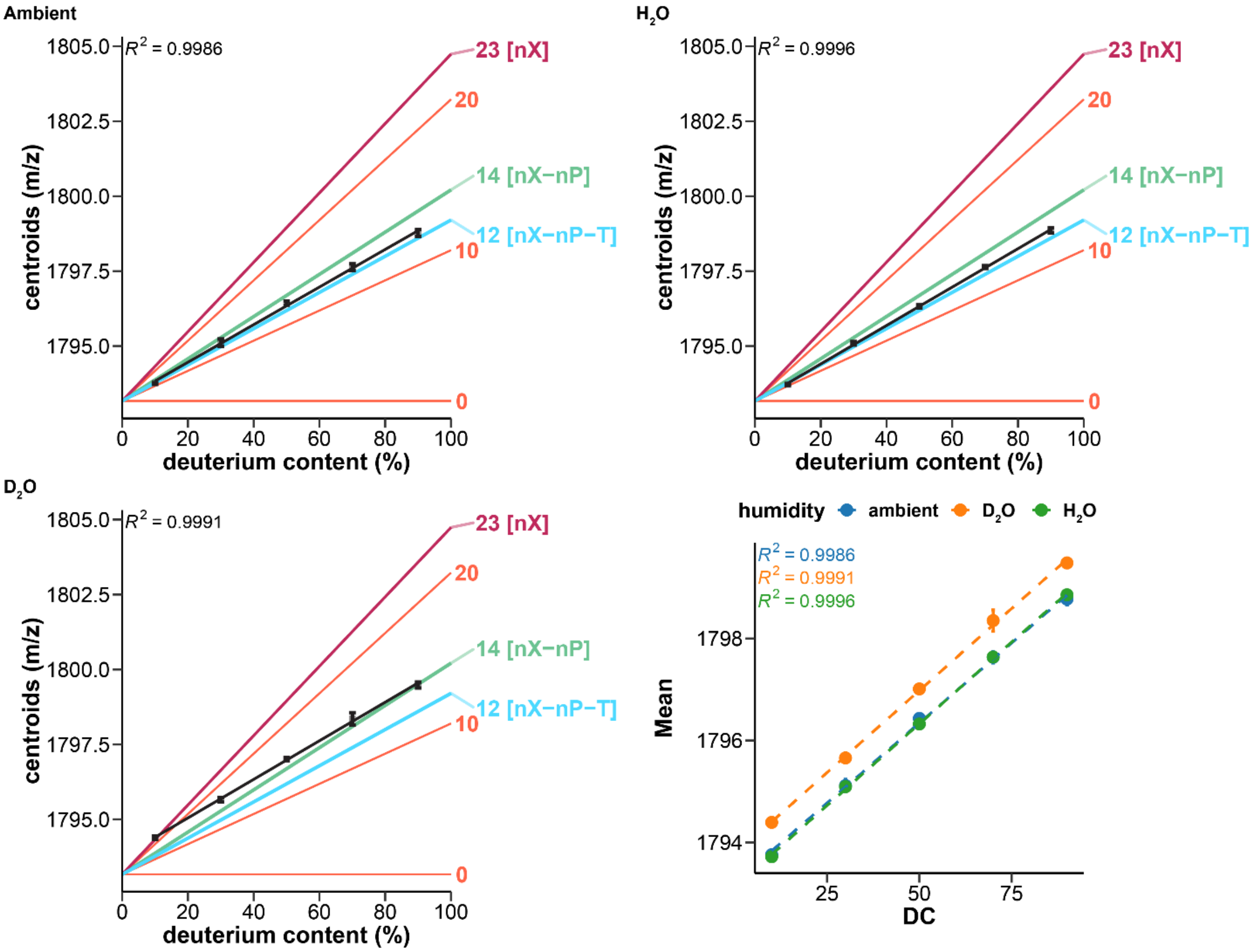


*Figure S16. Exchangeable sites in T12 at z = 2-, in 100 mM TMAA, 1 mM KCl, in different atmosphere (top left: ambient humidity, top right: $H_2O$-doped, bottom left: $D_2O$-doped, bottom right: overlaid data). Measured centroid masses against the solution deuterium content (standard deviation on duplicate measurements as error bars, linear fit as black lines), compared to notable theoretical masses: maximum number of potentially exchangeable sites (*nX*; purple), no phosphate exchanged (*nX–nP*; green), no phosphate and termini exchanged (*nX–nP–T*; cyan), and every ten sites (coral).*

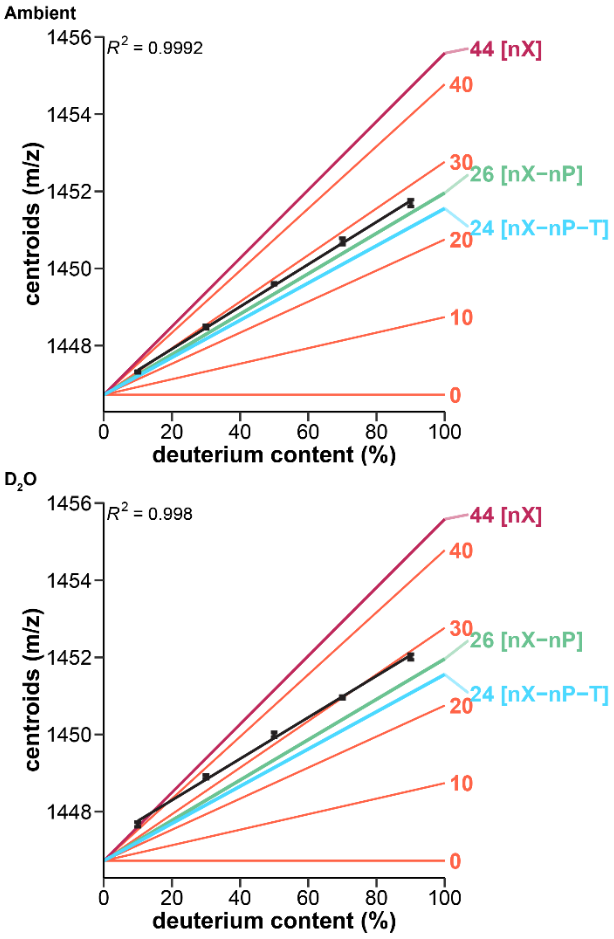


*Figure S17. Exchangeable sites in T24 at z = 5-, in 100 mM TMAA, 1 mM KCl, in different atmosphere (top left: ambient humidity, top right: $H_2O$-doped, bottom left: $D_2O$-doped, bottom right: overlaid data). Measured centroid masses against the solution deuterium content (standard deviation on duplicate measurements as error bars, linear fit as black lines), compared to notable theoretical masses: maximum number of potentially exchangeable sites (*nX*; purple), no phosphate exchanged (*nX–nP*; green), no phosphate and termini exchanged (*nX–nP–T*; cyan), and every ten sites (coral).*

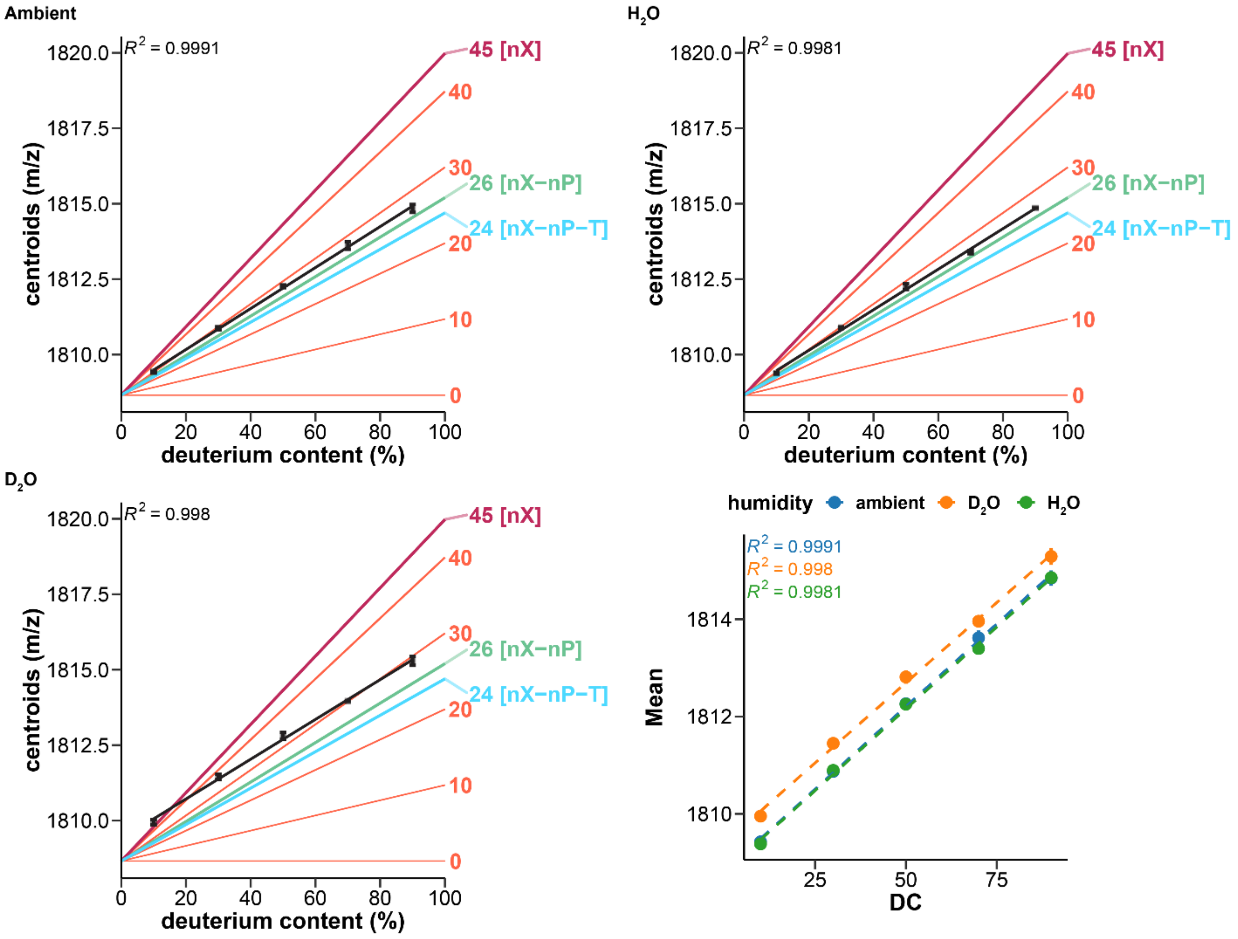


*Figure S18. Exchangeable sites in T24 at z = 4-, in 100 mM TMAA, 1 mM KCl, in different atmosphere (top left: ambient humidity, top right: $H_2O$-doped, bottom left: $D_2O$-doped, bottom right: overlaid data). Measured centroid masses against the solution deuterium content (standard deviation on duplicate measurements as error bars, linear fit as black lines), compared to notable theoretical masses: maximum number of potentially exchangeable sites (*nX*; purple), no phosphate exchanged (*nX–nP*; green), no phosphate and termini exchanged (*nX–nP–T*; cyan), and every ten sites (coral).*

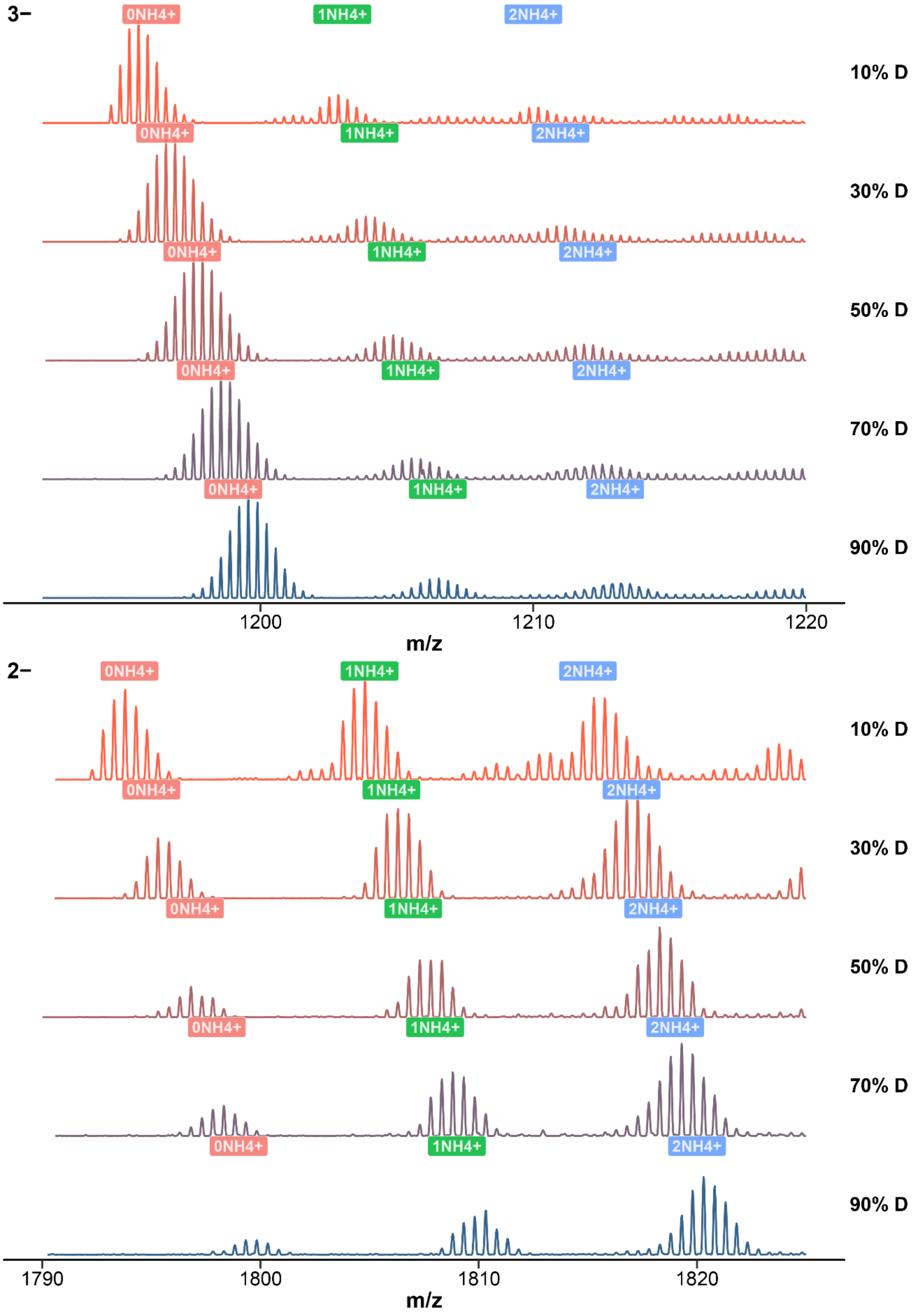


*Figure S19. Native mass spectra of T12 zoomed at the 3- (top) and 2- (bottom) charge states, for solutions containing 10 to 90% deuterium in 100 mM ammonium acetate. The number of bound ammoniums is labeled over the peaks.*

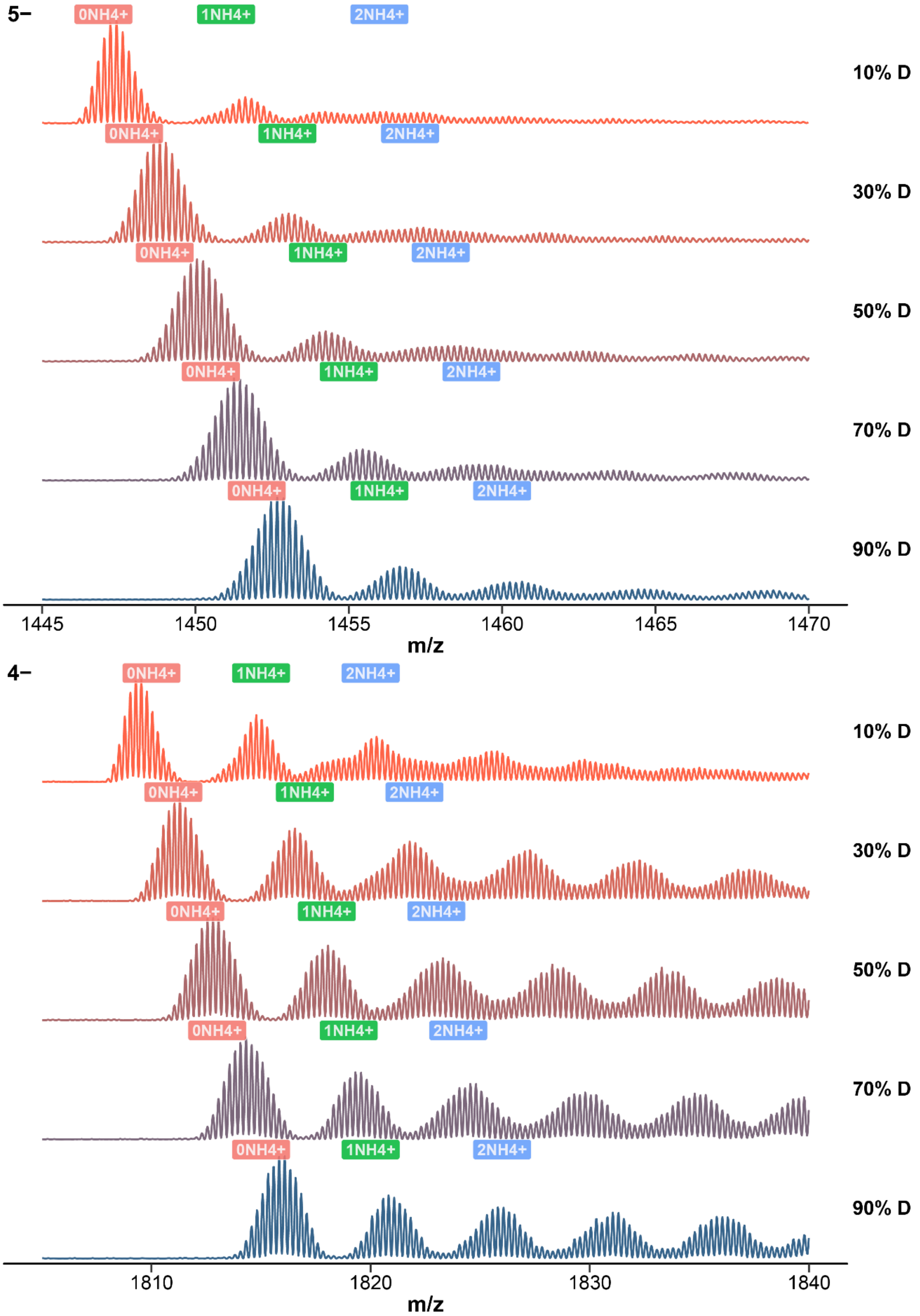


*Figure S20. Native mass spectra of T24 zoomed at the 5- (top) and 4- (bottom) charge states, for solutions containing 10 to 90% deuterium in 100 mM ammonium acetate. The number of bound ammoniums is labeled over the peaks.*

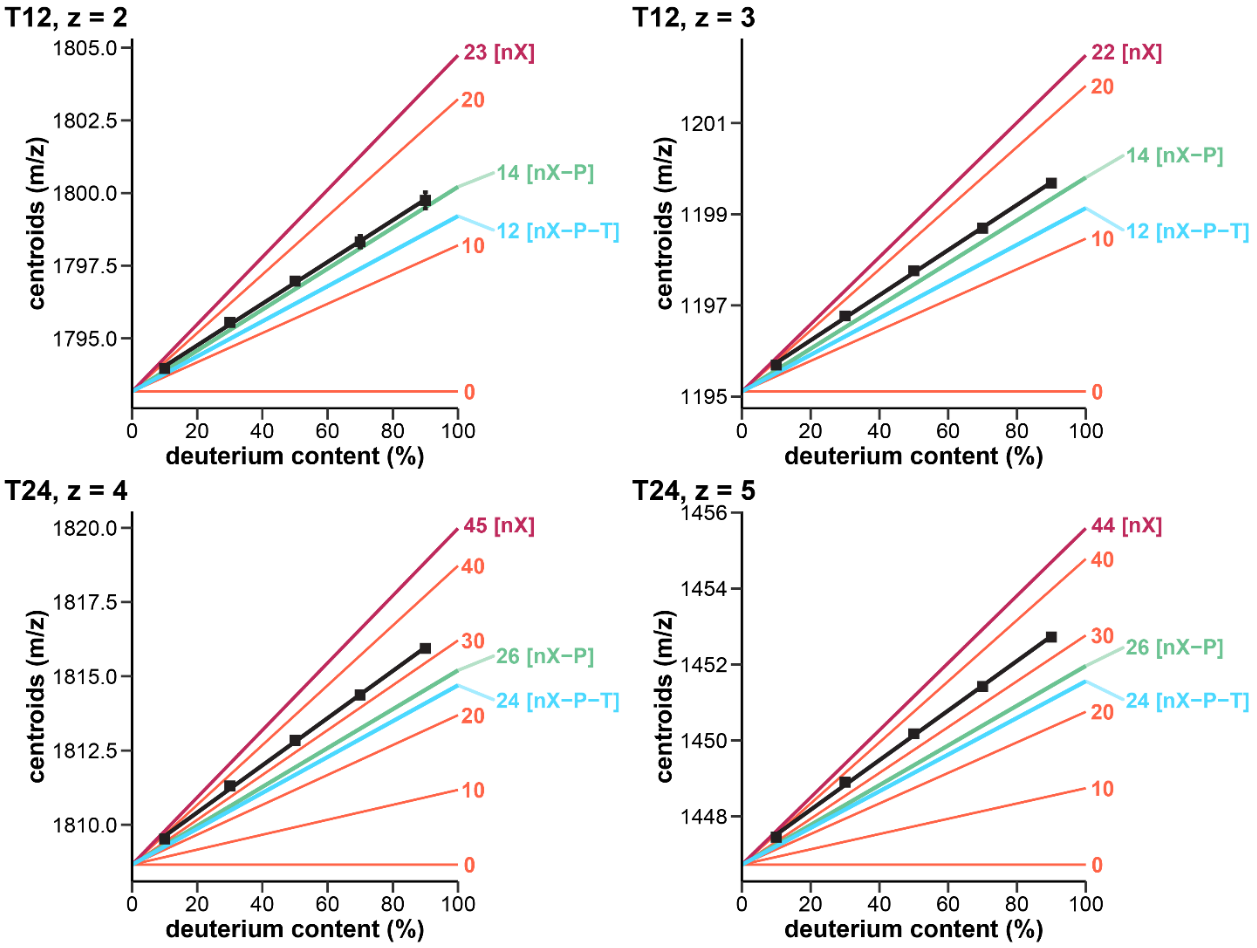


*Figure S21. Exchangeable sites in T12 and T24 in 100 mM ammonium acetate, at two different charge states. Measured centroid masses against the solution deuterium content (standard deviation on duplicate measurements as error bars, linear fit as black lines), compared to notable theoretical masses: maximum number of potentially exchangeable sites (*nX*; purple), no phosphate exchanged (*nX–nP*; green), no phosphate and termini exchanged (*nX–nP–T*; cyan), and every ten sites (coral).*

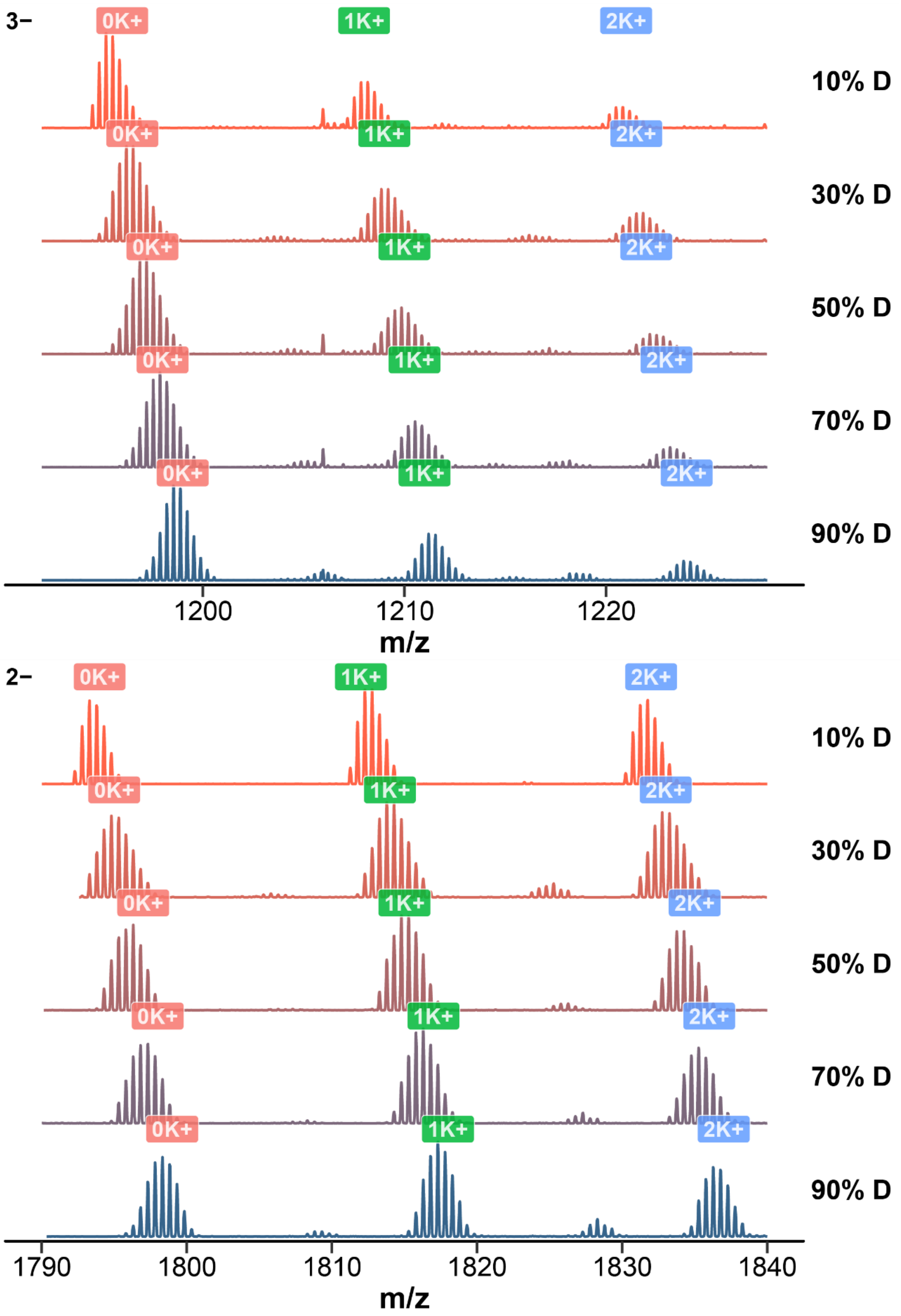


*Figure S22. Native mass spectra of T12 zoomed at the 3- (top) and 2- (bottom) charge states, for solutions containing 10 to 90% deuterium in 100 mM TMAA, 1 mM KCl, using the denaturing tune. The number of bound potassium is labeled over the peaks.*

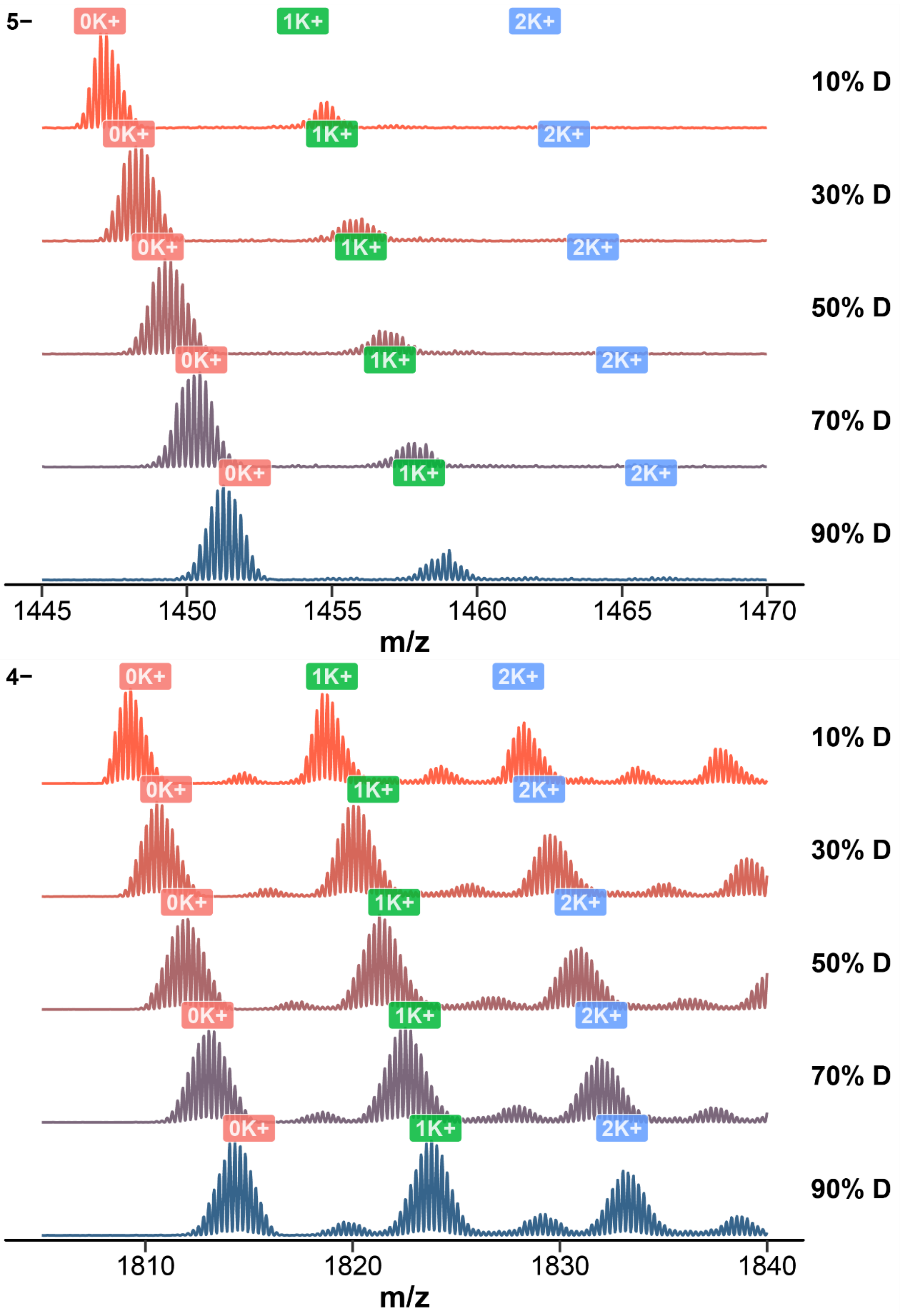


*Figure S23. Native mass spectra of T24 zoomed at the 5- (top) and 4- (bottom) charge states, for solutions containing 10 to 90% deuterium in 100 mM TMAA, 1 mM KCl, using the denaturing tune. The number of bound potassium is labeled over the peaks.*

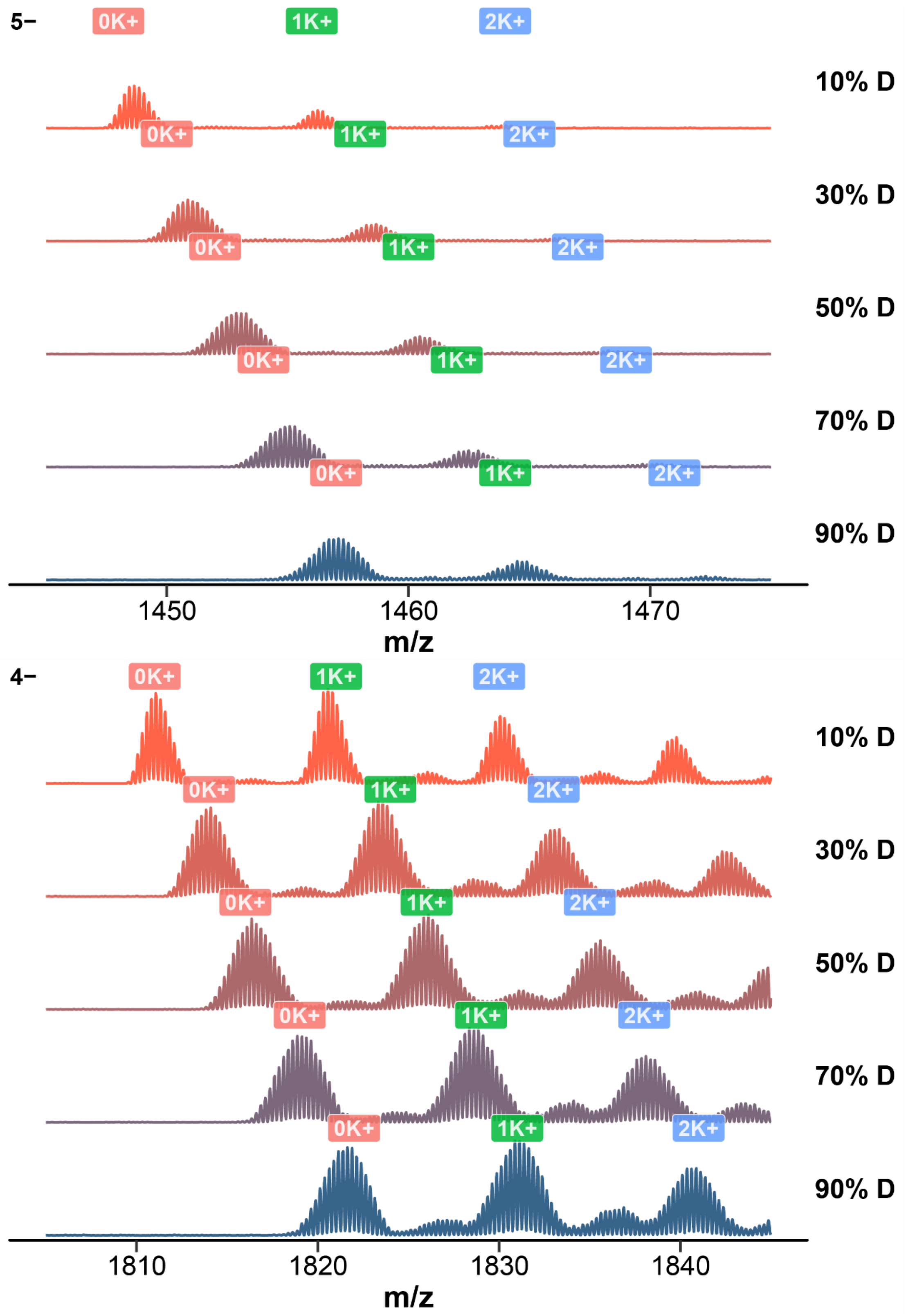


*Figure S24. Native mass spectra of 23nonG4 zoomed at the 5- (top) and 4- (bottom) charge states, for solutions containing 10 to 90% deuterium in 100 mM TMAA, 1 mM KCl, using the denaturing tune. The number of bound potassium is labeled over the peaks.*

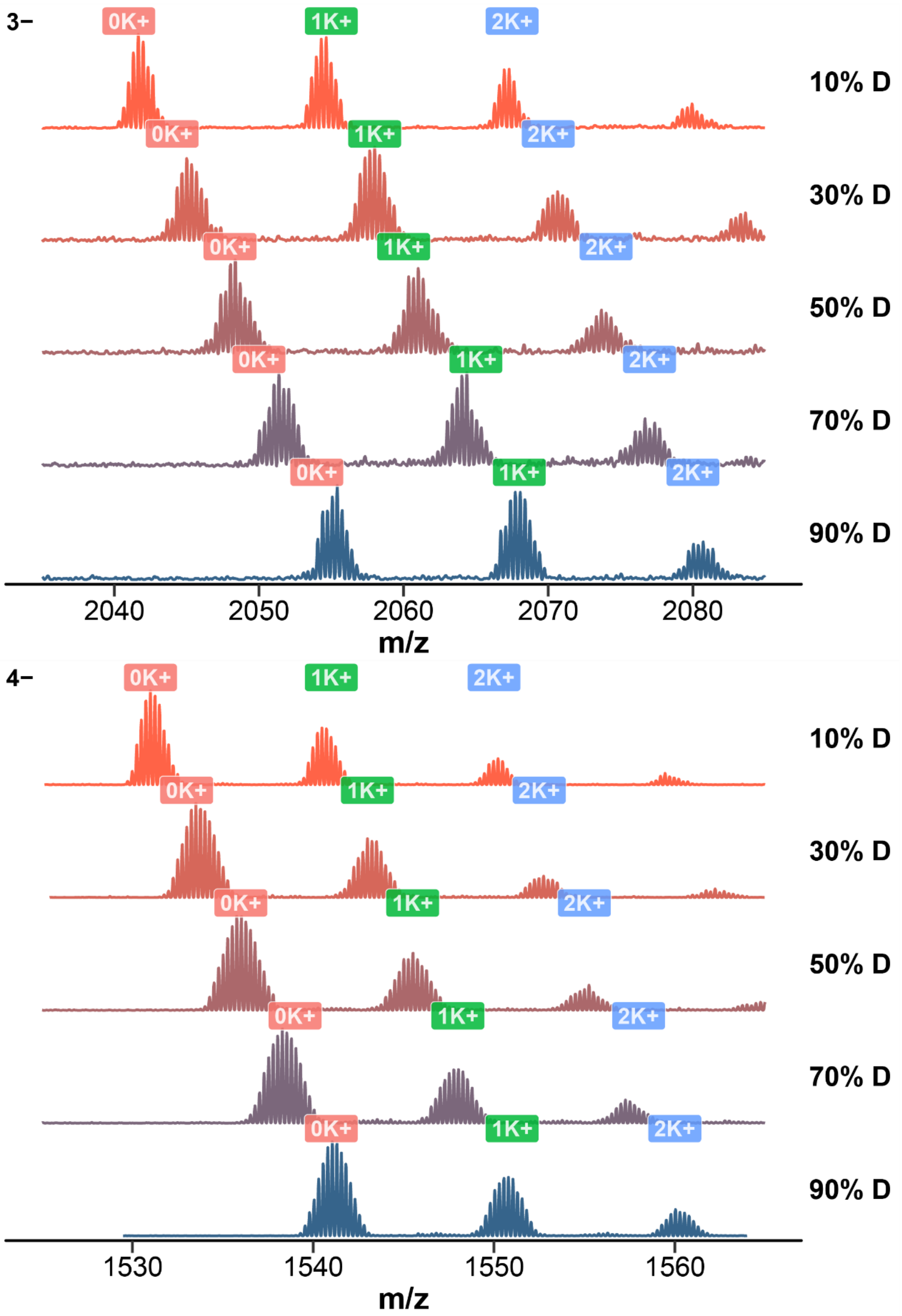


*Figure S25. Native mass spectra of Nmyc zoomed at the 4- (top) and 3- (bottom) charge states, for solutions containing 10 to 90% deuterium in 100 mM TMAA, 1 mM KCl, using the denaturing tune. The number of bound potassium is labeled over the peaks.*

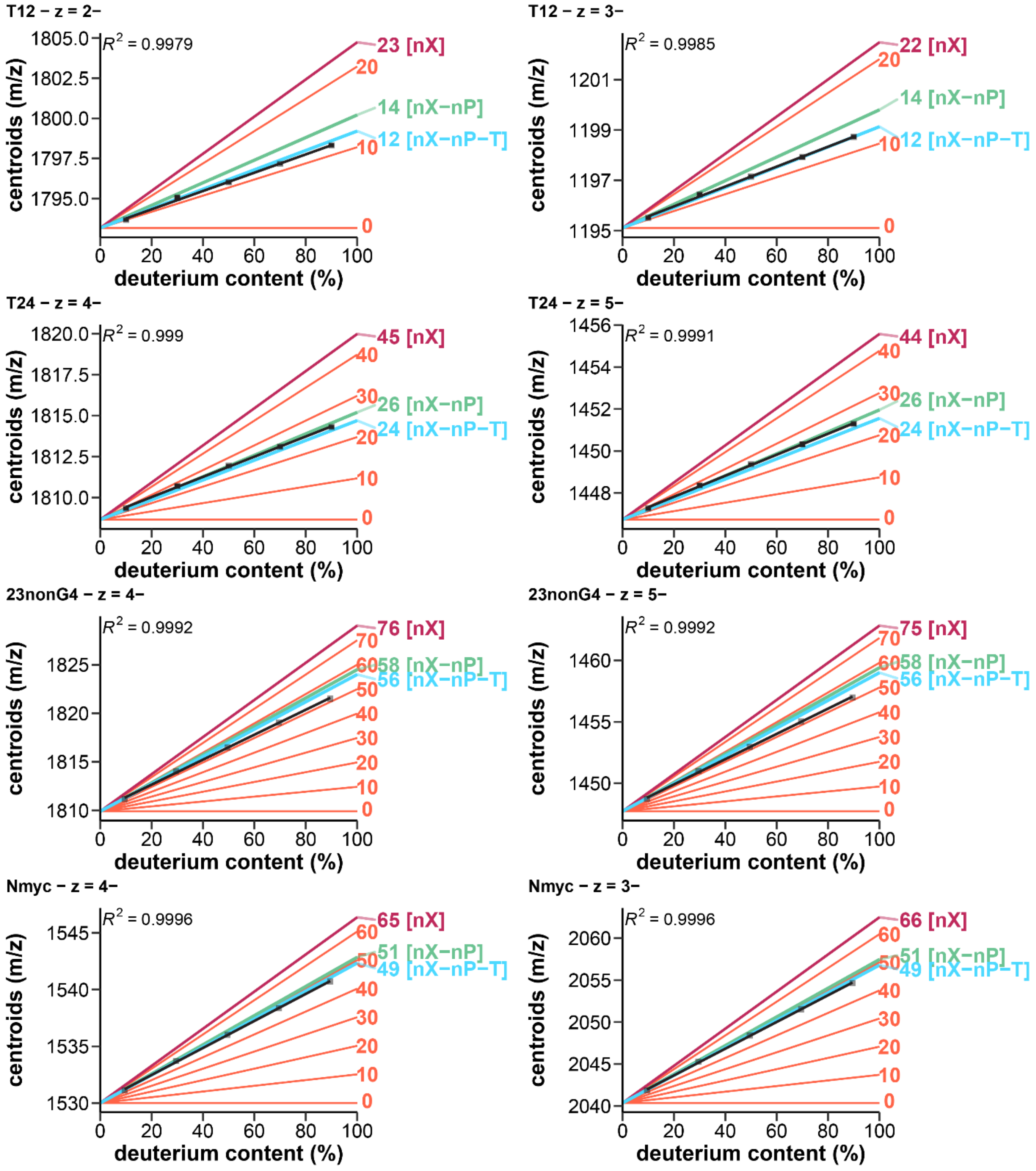


*Figure S26. Exchangeable sites in T12, T24, 23nonG4 and N-Myc in 100 mM TMAA, 1 mM KCl, using the denaturing tune (average of two measurements, standard deviation shown as error bars): Measured centroid masses against the solution deuterium content (black squares, linear fit: black lines), compared to notable theoretical masses: maximum number of potentially exchangeable sites (nX; purple), no phosphate exchanged (nX–nP; green), no phosphate and termini exchanged (nX–nP–T; cyan), and every ten sites (coral).*

#### Implications for converting the *m/z* distribution into unexchanged sites equivalent and interpreting NUS values.

##### Calculation of theoretical isotopic distribution of oligonucleotides

The theoretical distributions were computed in R. The code is given and briefly commented below. When the output is explicitly printed for illustrating purpose, it is preceded by ##, and is therefore not part of the code to run.

##### Libraries

The code can be executed using base R[21] complemented by the Tidyverse package collection.[22]

```
library(tidyverse)
```

##### User input

The user must input the charge state `z`, number of potassium adduct `K`, sequence `seq`, the molar percentage of solution deuterium `DC`, and number of exchanging sites `nX`. The code described below is exemplified for the oligonucleotide T95-2T with 2 potassium adducts, at the 4- charge state, from a 9%D solution. The number of exchanging sites was set at 44.

```
z <- 4 #charge

K <- 2 #potassium adduct

seq <- "TTGGGTGGGTGGGTGGGT"

nX <- 44

DC <- 9 #%D
```

##### Sequence, phosphates and chemical formula

The numbers of each nucleotide are extracted from the sequence, then the total number of nucleotides `nb_nt` is calculated, which gives access to the number of phosphates `nb_PO`.

```
seq2 <- data.frame(number=1:1, string=seq, stringsAsFactors = F)

nbA <- str_count(seq2$string, "A")

nbT <- str_count(seq2$string, "T")

nbG <- str_count(seq2$string, "G")

nbC <- str_count(seq2$string, "C")

#Number of nucleotides
nb_nt <- nbA + nbT + nbG + nbC

#Number of phosphates
nP <- nb_nt - 1

print(paste('the number of phosphates is', nP))

## [1] "the number of phosphates is 17"
```

The number of each atom is readily calculated from the elemental composition of each nucleotide, having already determined the base composition from the sequence. Note that the number of hydrogens takes into account the charge and number of potassium adducts, and therefore the chemical formula is the one of the ion in the mass spectrometer.

```
nC <- nbA*10 + nbG*10 + nbC*9 + nbT*10

nH <- nH <- nbA*12 + nbG*12 + nbC*12 + nbT*13 + 1 - z - K

nO <- nbA*5 + nbG*6 + nbC*6 + nbT*7 - 2

nN <- nbA*5 + nbG*5 + nbC*3 + nbT*2

nK <- K

chem.formula <- paste('Chemical formula: ',
                      "C", nC, "H", nH, "O", nO,
                      "N",nN, "P", nP, "K", nK,
                      sep = '')
chem.formula

## [1] "Chemical formula: C180H217O112N72P17K2"
```

#### Mass calculations

The masses and abundances were obtained on the website of the CIAWW. The abundance of deuterium is calculated from the content given by the user `DC`.

```
#Isotope abundances (averaged).
#Source: Isotopic compositions of the elements 2017. Available online at www.ciaaw.org.
  listIso <- list(
      H = c(0.999855, 0.000145), #Natural 1H ``and 2H isotope abundances
      C = c(0.9894, 0.0106),
      N = c(0.996205, 0.003795),
      O = c(0.99757, 0.0003835, 0.002045),
      D = c(100-DC, DC)/100,
      #computed 1H and 2H isotope abundances for exchangeable
      #sites depending on the deuterium content DC
      P = c(1),
      K = c(0.93258144, 0.0001171, 0.06730244)
    )

#Atomic masses.
#Current atomic masses available online at www.ciaaw.org
#Based on Wang,M., Audi,G., Kondev,F.G., Huang,W.J., Naimi,S., et al. (2017)
#Chinese Phys. C, 41, 030003.
  listMass <- list(
    H = c(1.0078250322, 2.0141017781),
    C = c(12, 13.003354835),
    N = c(14.003074004, 15.000108899),
    O = c(15.994914619, 16.999131757, 17.999159613),
    D = c(1.0078250322, 2.0141017781),
    P = c(30.973761998),
    K = c(38.96370649, 39.9639982, 40.96182526)
  )
```

The monoisotopic mass `MonoMW` is calculated from the chemical formula and the lightest isotope mass for each atom. The corresponding m/z `Monomz` is obtained by only dividing by the charge `z` (the *z* protons are already subtracted in the `Chem.formula`)

```
MonoMW <- nC*listMass$C[1] + nH*listMass$H[1] +
  nN*listMass$N[1] + nO*listMass$O[1] +
  nP*listMass$P[1] + nK*listMass$K[1]

print(paste('the monoisotopic mass is', MonoMW))

## [1] "the monoisotopic mass is 5782.8311645494"

Monomz <- if (z>0) {
  MonoMW/z
} else {
  MonoMW
}

print(paste('the monoisotopic m/z is', Monomz))

## [1] "the monoisotopic m/z is 1445.70779113735"
```

Similarly, the average mass `AveMW` and m/z `Avemz` are calculated, this time by adding all isotopes weighed by their abundances. Note that the number of exchanging sites `nX` is subtracted from `nH` because `nH` is the total number of H, including the exchangeable ones.

```
 #Average mass calculation from number of atoms, isotopic masses and abundances.
AveMW <- nC*(listIso$C[1]*listMass$C[1] + listIso$C[2]*listMass$C[2]) +
         (nH-nX)*(listIso$H[1]*listMass$H[1] + listIso$H[2]*listMass$H[2]) +
         nN*(listIso$N[1]*listMass$N[1] + listIso$N[2]*listMass$N[2]) +
         nO*(listIso$O[1]*listMass$O[1] + listIso$O[2]*listMass$O[2] +
         listIso$O[3]*listMass$O[3]) +
         nX*(listIso$D[1]*listMass$D[1] + listIso$D[2]*listMass$D[2]) +
         nP*(listIso$P[1]*listMass$P[1]) +
         nK*(listIso$K[1]*listMass$K[1] + listIso$K[2]*listMass$K[2] +
         listIso$K[3]*listMass$K[3])

print(paste('the average mass is', AveMW))

## [1] "the average mass is 5789.79685919195"

  Avemz <- if (z>0) {
    AveMW/z
  } else {
    AveMW/z
  }

print(paste('the average m/z is', Avemz))

## [1] "the average m/z is 1447.44921479799"
```

#### Isotopic distribution

The number of peaks `nrPeaks` to compute is defined by the user. Here, 32 are enough to cover the full distribution.

```
nrPeaks <- 32
```

The isotope abundances are redefined to make the use of FFT easier. It takes into account `DC` and `nrPeaks`.

```
Iso_Pattern <- c12 <- rep(0, nrPeaks); h1 <- rep(0, nrPeaks); n14 <- rep(0, nrPeaks);
o16 <- rep(0, nrPeaks); p31 <- rep(0, nrPeaks); k39 <- rep(0,nrPeaks);
h2 <- rep(0, nrPeaks);
```

```
#Isotope abundances
h1[1] = 0.999855; h1[2] = 0.000145 ;#Natural abundances of H isotopes
h2[1] = (1-DC/100); h2[2] = DC/100; #Calculated abundances for exchangeable sites
c12[1] = listIso$C[1]; c12[2] = listIso$C[2];
n14[1] = listIso$N[1]; n14[2] = listIso$N[2];
o16[1] = listIso$O[1]; o16[2] = listIso$O[2]; o16[3] = listIso$O[3];
p31[1] = 1.0;
k39[1] = listIso$K[1]; k39[2] = listIso$K[2]; k39[3] = listIso$K[3];
```

The abundances of the isotopic peaks are computed from the atomic abundances and the `Chem.formula`, by Fast Fourier Transform (FFT). This approach is based on reference 23.

```
Iso_Pattern <- Re(fft(fft(c12)^nC*fft(h1)^(nH-nX)*fft(n14)^nN * fft(o16)^nO * fft(p31)^nP * ff
t(k39)^nK * fft(h2)^nX,
                         inverse=TRUE))/length(c12)
Iso_Pattern

##  [1] 1.137651e-03 7.534397e-03 2.511751e-02 5.620804e-02 9.499299e-02
##  [6] 1.293224e-01 1.477177e-01 1.455920e-01 1.263744e-01 9.811595e-02
## [11] 6.897117e-02 4.432989e-02 2.626140e-02 1.443595e-02 7.405374e-03
## [16] 3.562366e-03 1.613817e-03 6.910346e-04 2.806029e-04 1.083659e-04
## [21] 3.990526e-05 1.404506e-05 4.734737e-06 1.531752e-06 4.763994e-07
## [26] 1.426765e-07 4.120838e-08 1.149411e-08 3.100163e-09 8.095359e-10
## [31] 2.048892e-10 5.031431e-11
```

The corresponding x-axis, i.e. the m/z axis, is finally generated, following the number of peaks wanted by the user `nrPeaks`. The data frame `peaks` contains the final distribution.

```
nbPeaks <- 1:nrPeaks

peaks <- data.frame("nbPeaks1" = unlist(nbPeaks), 'MonoMW1' = MonoMW) %>%
      mutate('mass.th' = MonoMW1 + (nbPeaks1-1)*1.0078250321) %>%
      mutate('mz.th' = mass.th/z) %>%
      select(mz.th) %>%
      cbind('Iso.Pattern' = Iso_Pattern) %>%
      mutate(Iso.Pattern = 1 - (max(Iso.Pattern)-Iso.Pattern)/(max(Iso.Pattern)-min(Iso.Patter
n)))

p.peaks <- ggplot(data = peaks, aes(x = mz.th, y = Iso.Pattern)) +
      geom_line(color = 'tomato', size = 1) +
      geom_point(color = 'tomato', size = 3) +
      geom_vline(xintercept = Avemz, linetype = 'dashed', color = 'steelblue', size = 1) +
      xlab("m/z") +
      ylab('normalized abundance') +
      theme(strip.text.y = element_blank(),
            strip.background = element_blank(),
            panel.border = element_blank(),
            panel.grid.major = element_line(colour = "grey50"),
            panel.grid.minor = element_line(colour = "grey"),
            panel.background = element_blank(),
            plot.background = element_blank(),
            axis.line.x = element_line(colour = "black", size = 0.75),
            axis.line.y = element_line(colour = "black", size = 0.75),
            axis.ticks.length=unit(0.1, "in"),
            axis.ticks.x = element_line(size = 0.75, colour = "black"),
            axis.ticks.y = element_line(size = 0.75, colour = "black"),
            axis.text.y = element_text(size = 16, color = "black", angle = 0),
            axis.text.x = element_text(size = 16, color = "black", angle = 0),
            axis.title.x = element_text(size=18,face="bold", color = 'black'),
            axis.title.y = element_text(size=18,face="bold", color = 'black'),
```

```
            plot.margin = margin(25, 0.5, 0.5, 0.5),
            legend.position="right",
            legend.box = "vertical",
            legend.title = element_text(size=18, face="bold", color = "black"),
            legend.key = element_rect(fill = "white"),
            legend.text = element_text(size=16, face="bold", color = "black"),
      )

p.peaks
```

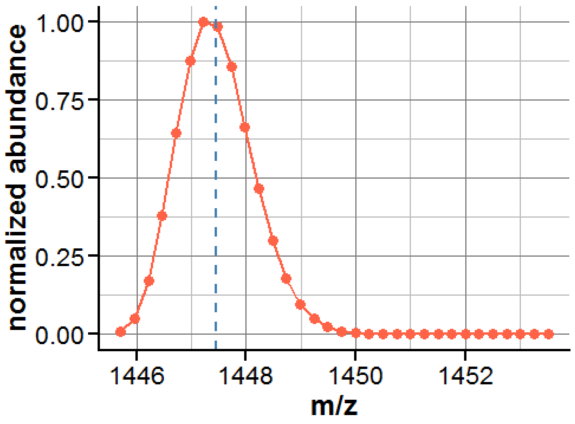

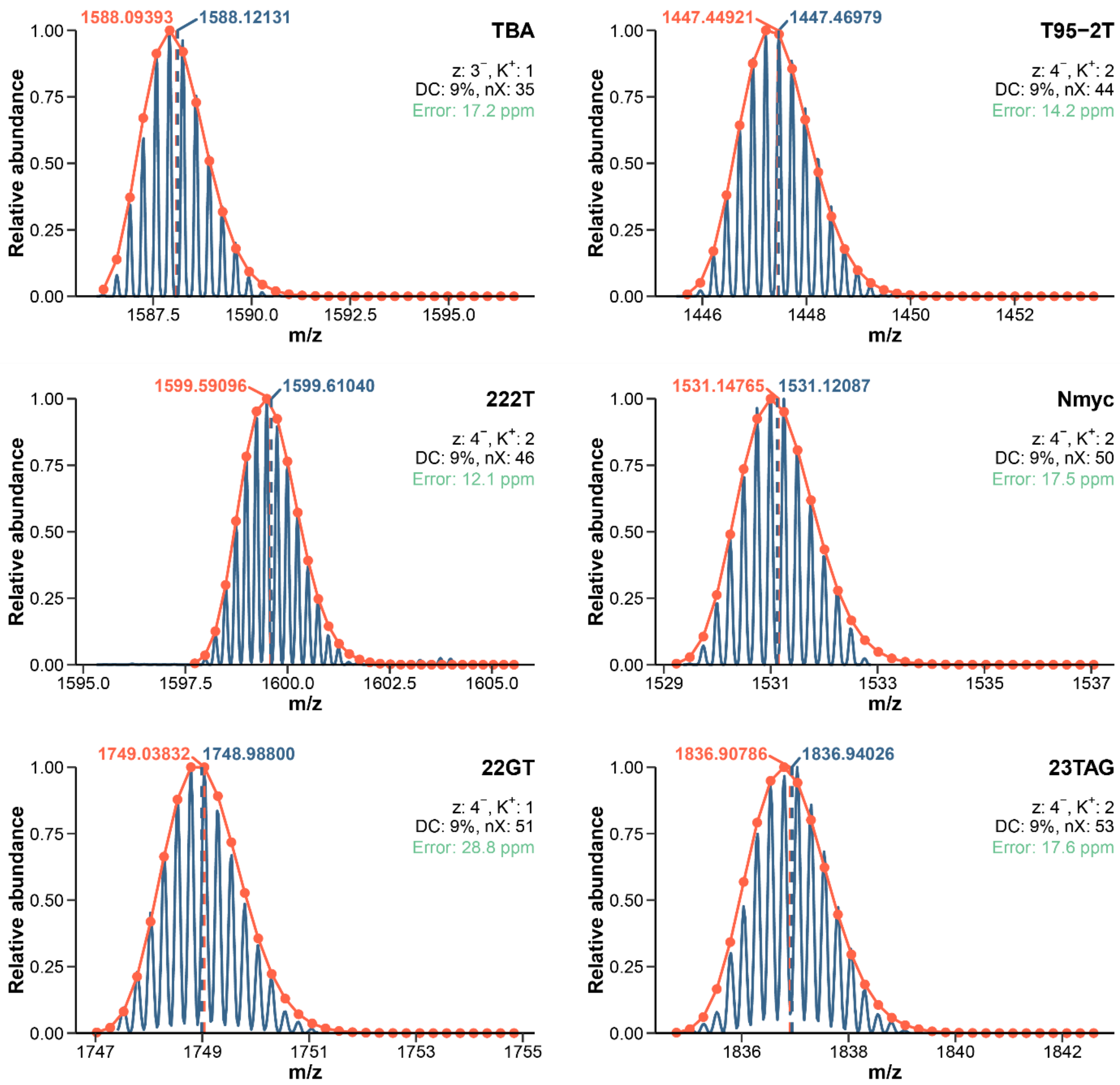


*Figure S27. Experimental isotopic distributions of samples fully exchanged in 9% $D_2O$ (blue) compared to the corresponding theoretical distributions (orange). The centroid masses are indicated with dashed vertical lines of the same colors. DC is the deuterium content, nX the number of exchanged sites.*

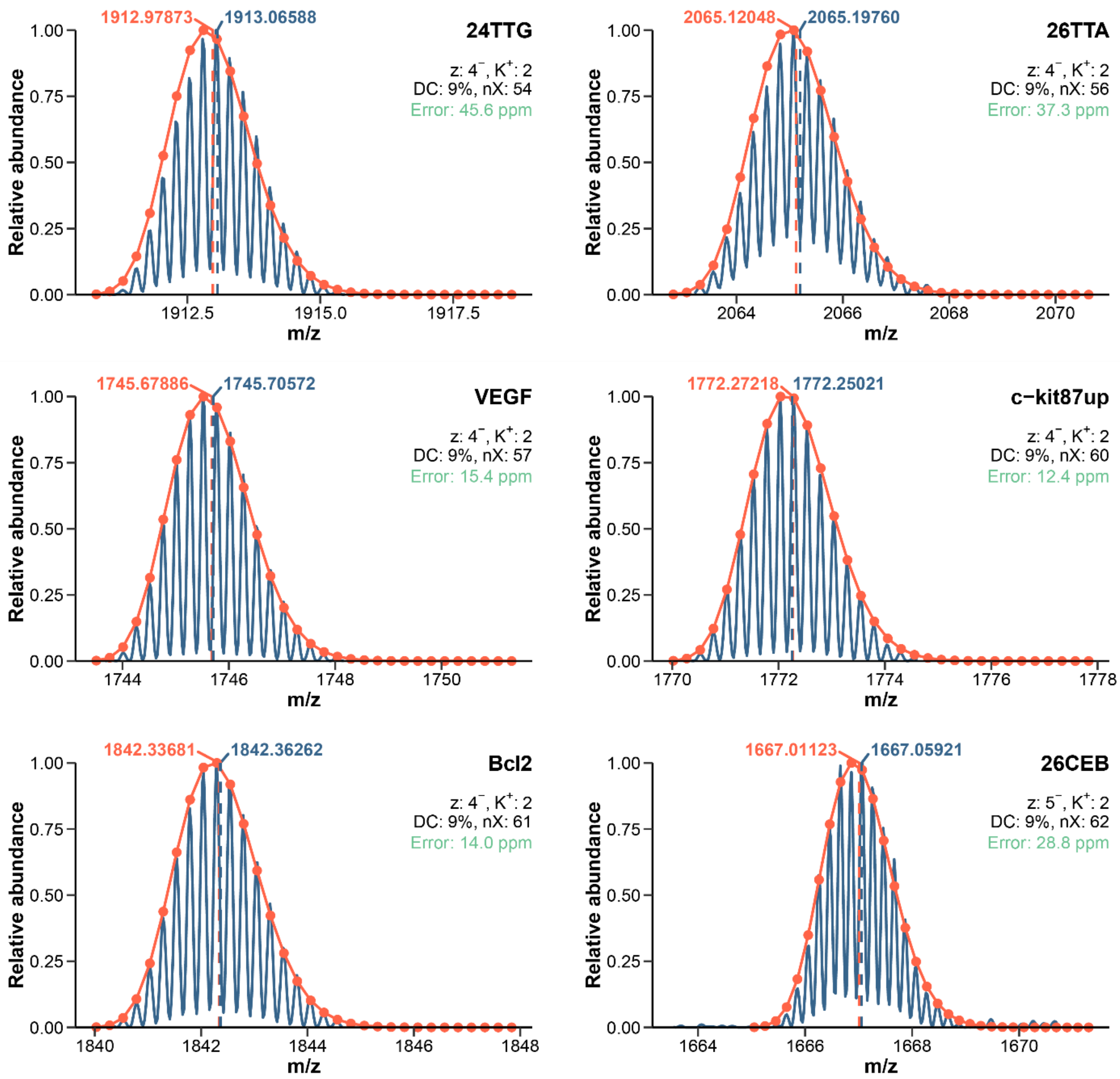


*Figure S28. Experimental isotopic distributions of samples fully exchanged in 9% $D_2O$ (blue) compared to the corresponding theoretical distributions (orange). The centroid masses are indicated with dashed vertical lines of the same colors. DC is the deuterium content, nX the number of exchanged sites.*

#### Exchange rates can be measured precisely on a large time scale, regardless of the charge state and adduct stoichiometry

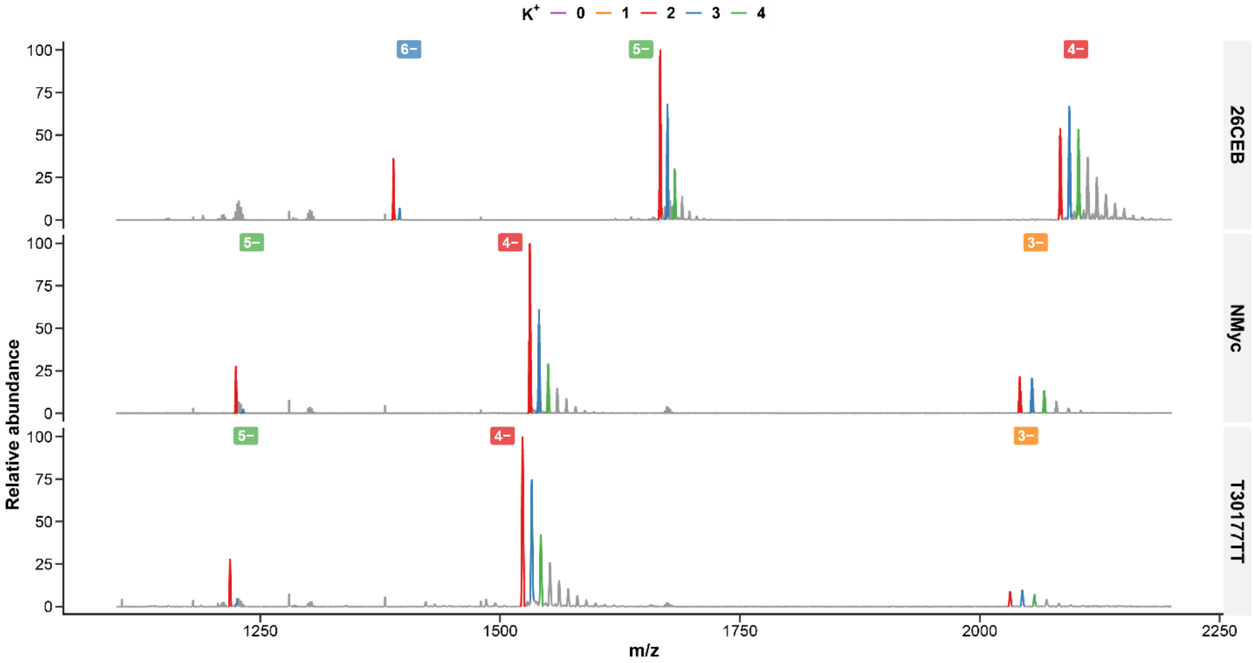


*Figure S29. Native mass spectra of 26CEB, N-myc, and T30177-TT (top to bottom). The charge states are indicated by a label, and for each charge state the 0—4 $K^+$ stoichiometries are colored.*

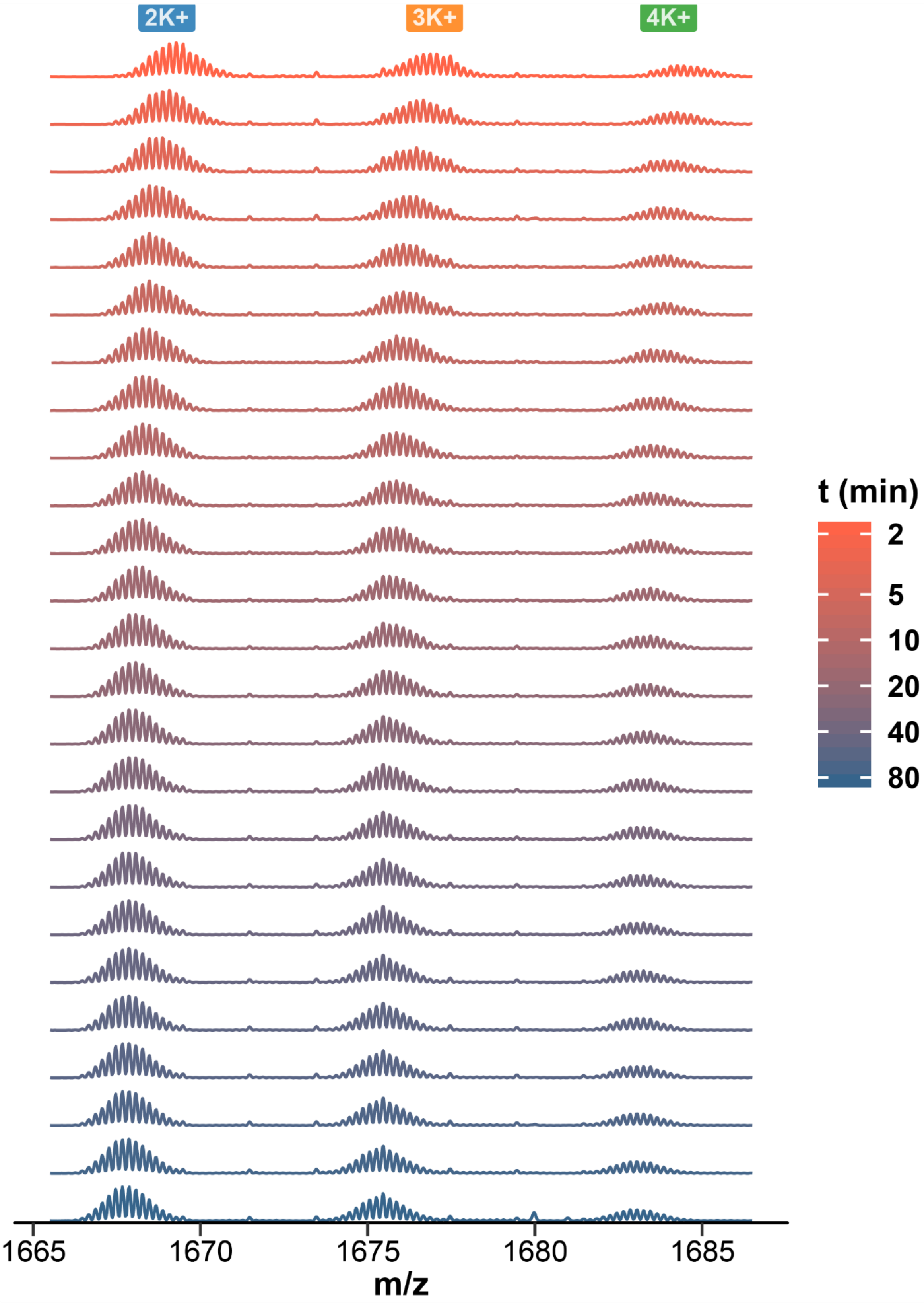


*Figure S30. HDX/native MS spectra of 26CEB, zoomed on the 5- charge state. 25 exchange time points are presented from top to bottom. Replicate 1.*

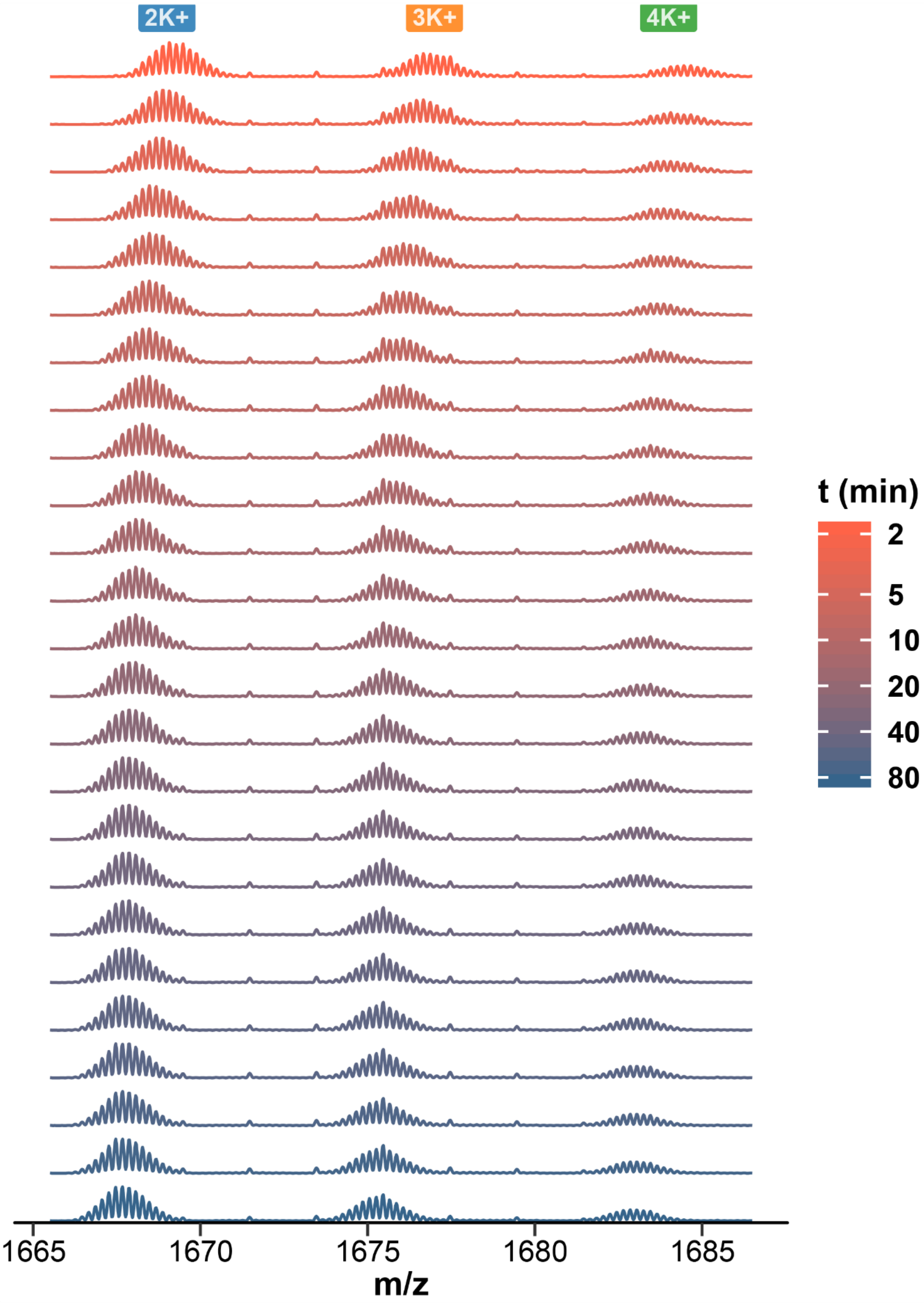


*Figure S31. HDX/native MS spectra of 26CEB, zoomed on the 5- charge state. 25 exchange time points are presented from top to bottom. Replicate 2.*

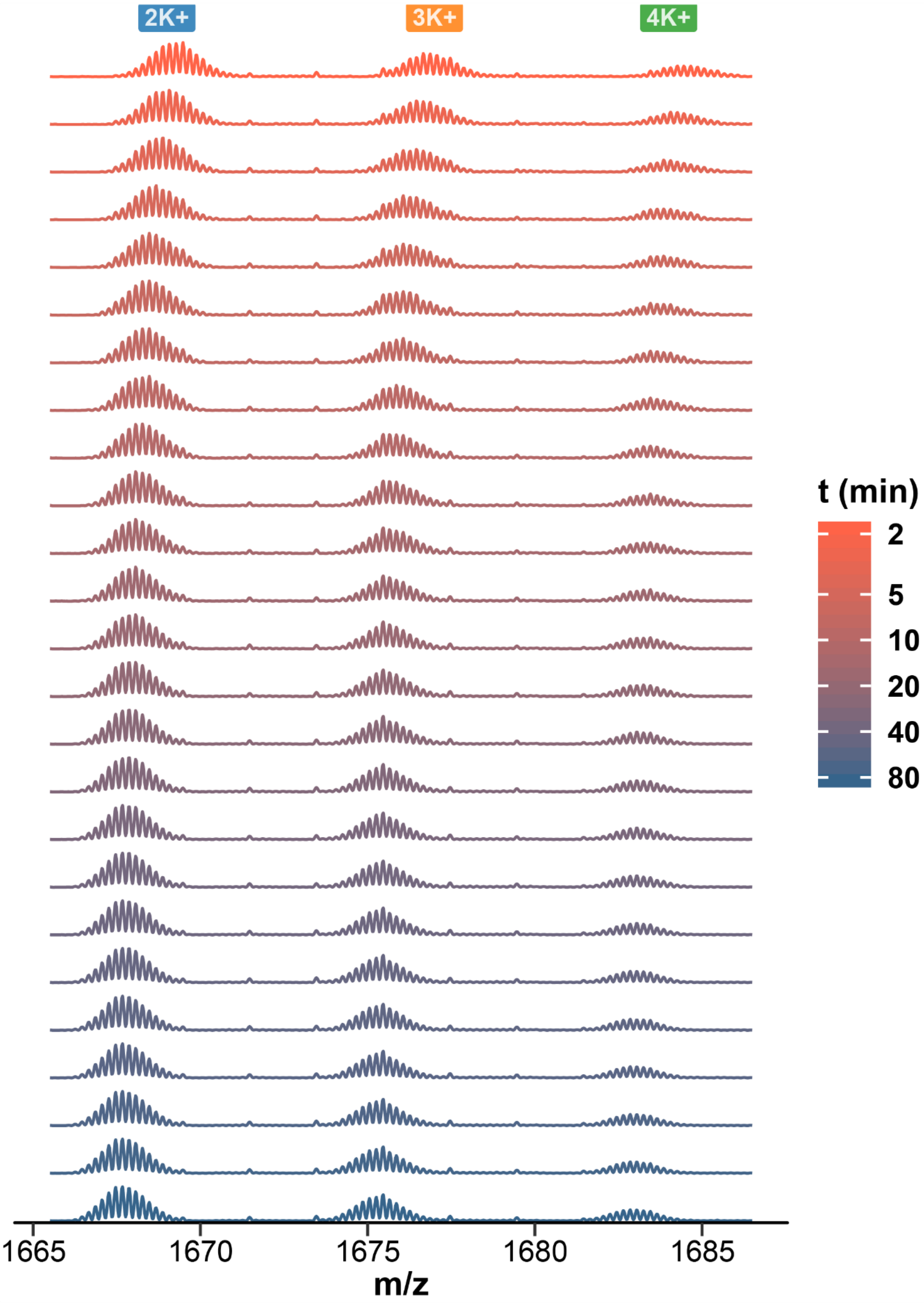


*Figure S32. HDX/native MS spectra of 26CEB, zoomed on the 5- charge state. 25 exchange time points are presented from top to bottom. Replicate 3.*

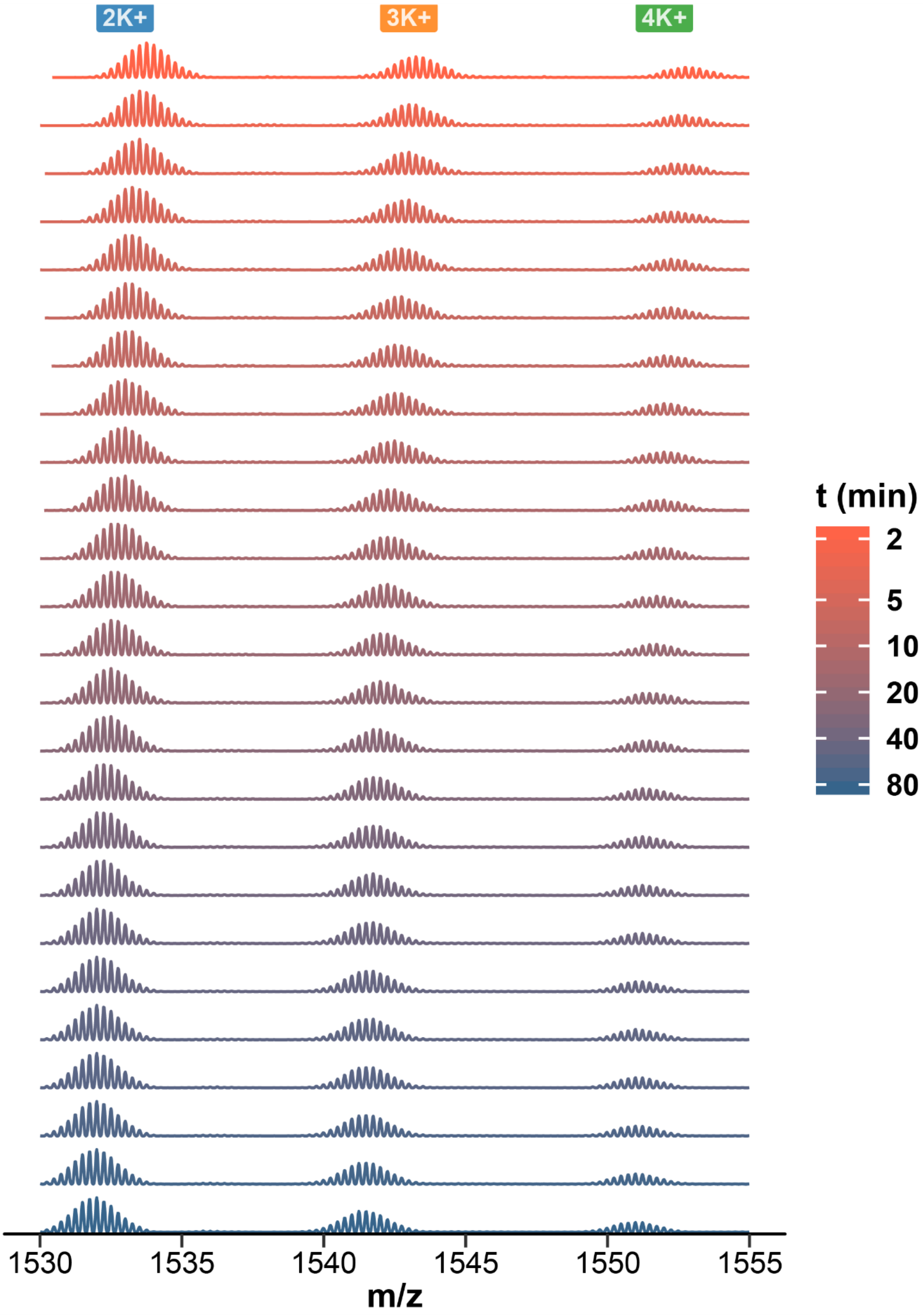


*Figure S33. HDX/native MS spectra of N-myc, zoomed on the 4- charge state. 25 exchange time points are presented from top to bottom. Replicate 1.*

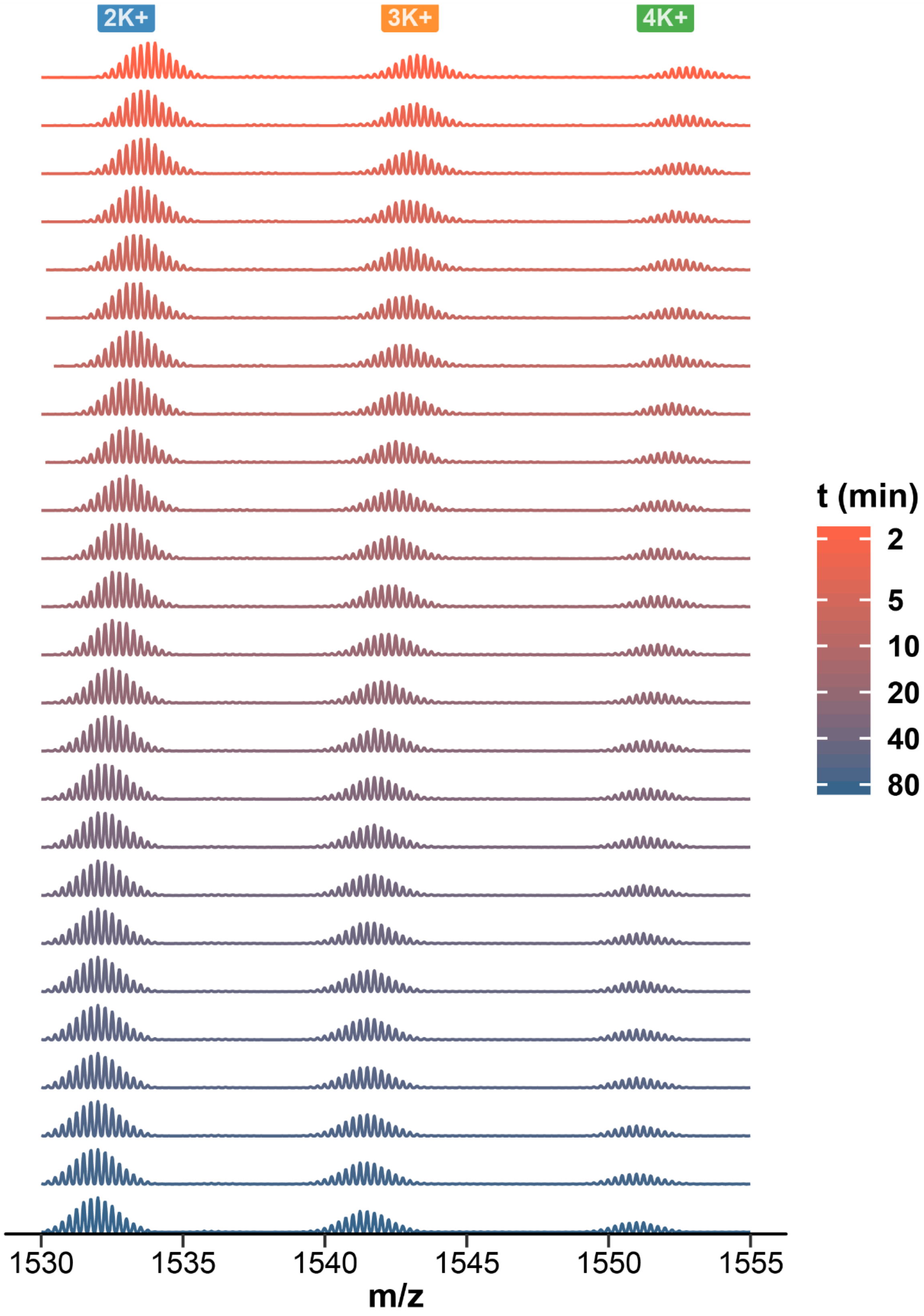


*Figure S34. HDX/native MS spectra of N-myc, zoomed on the 4- charge state. 25 exchange time points are presented from top to bottom. Replicate 2.*

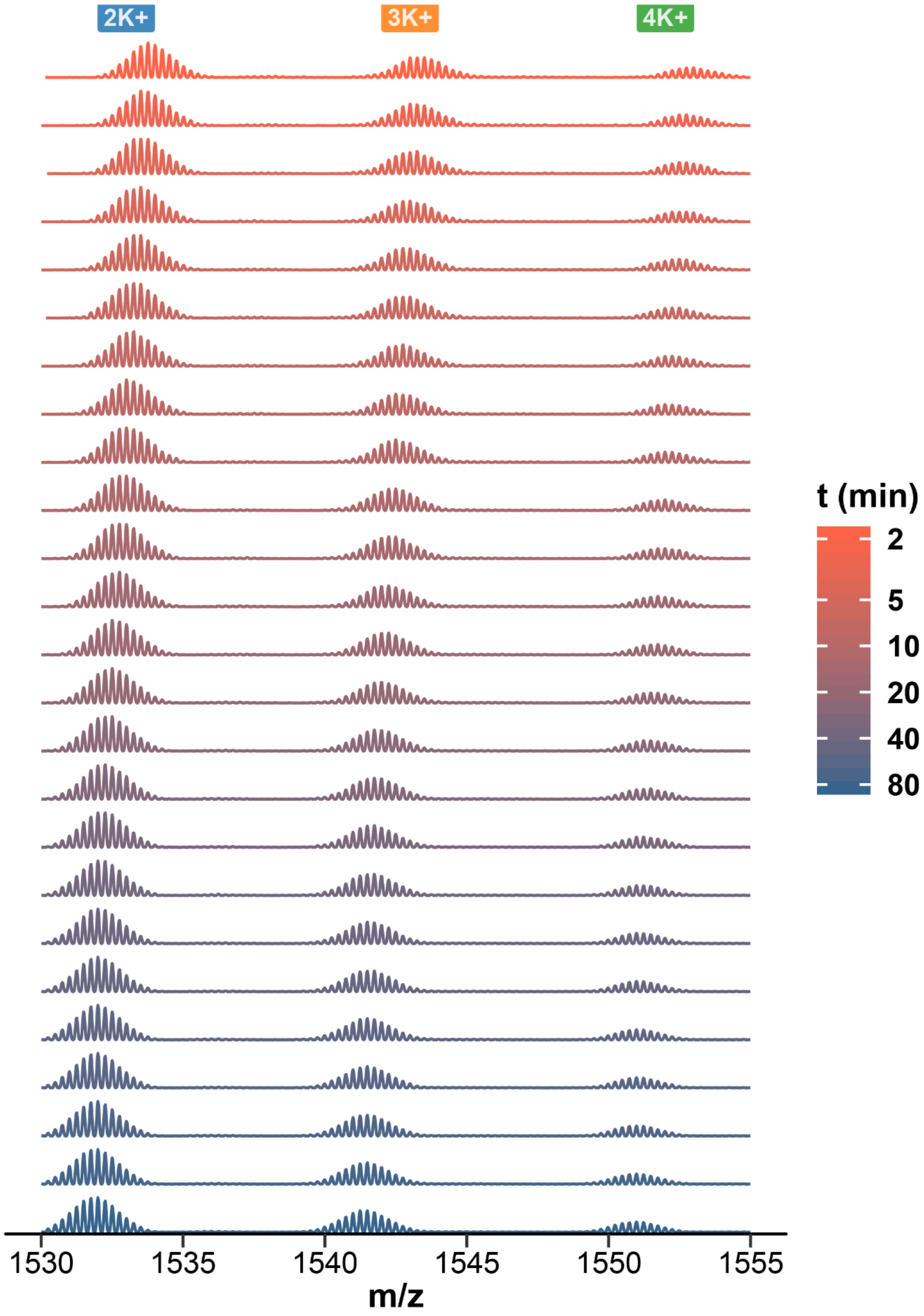


*Figure S35. HDX/native MS spectra of N-myc, zoomed on the 4- charge state. 25 exchange time points are presented from top to bottom. Replicate 3.*

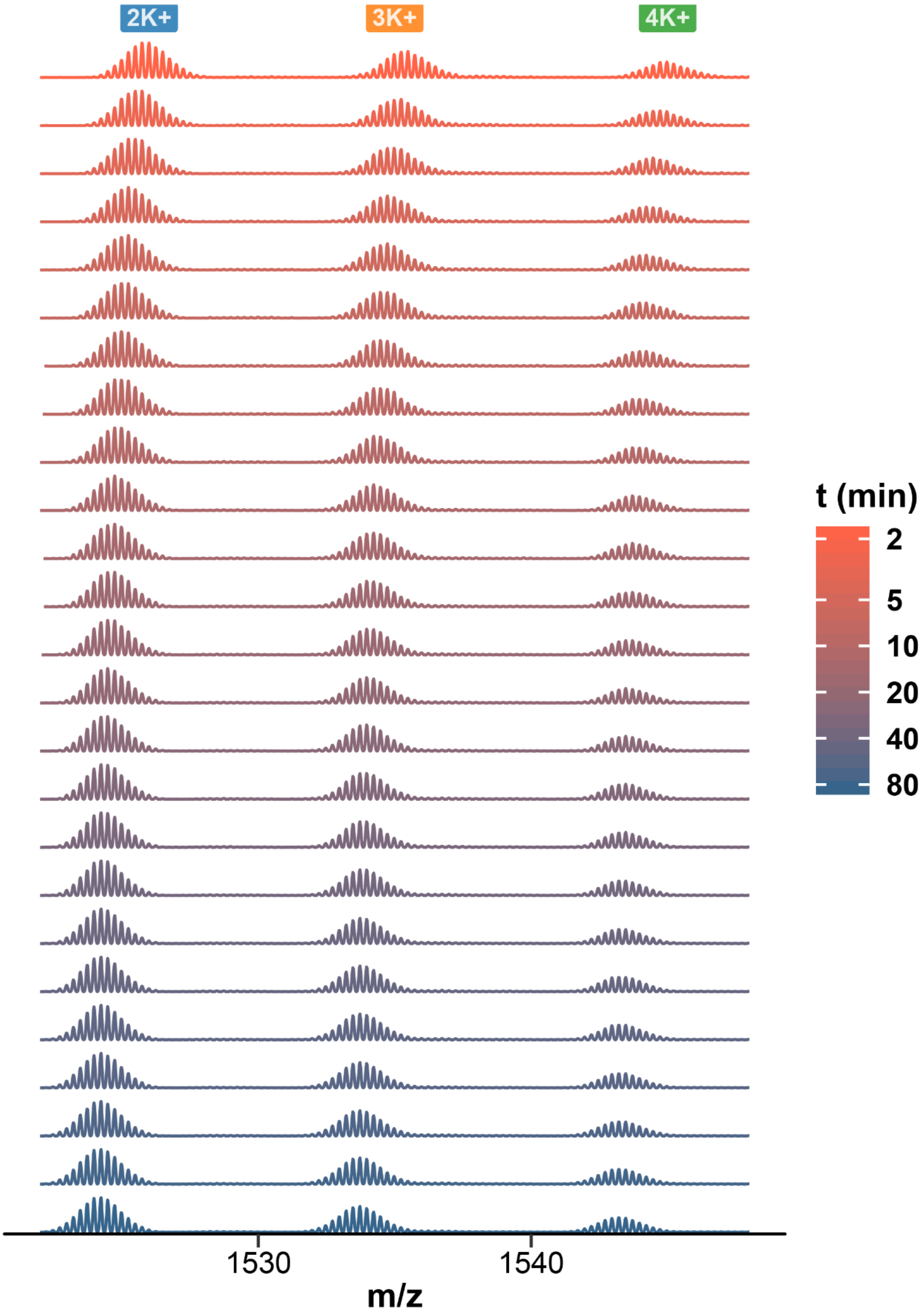


*Figure S36. HDX/native MS spectra of T30177-TT, zoomed on the 4- charge state. 25 exchange time points are presented from top to bottom. Replicate 1.*

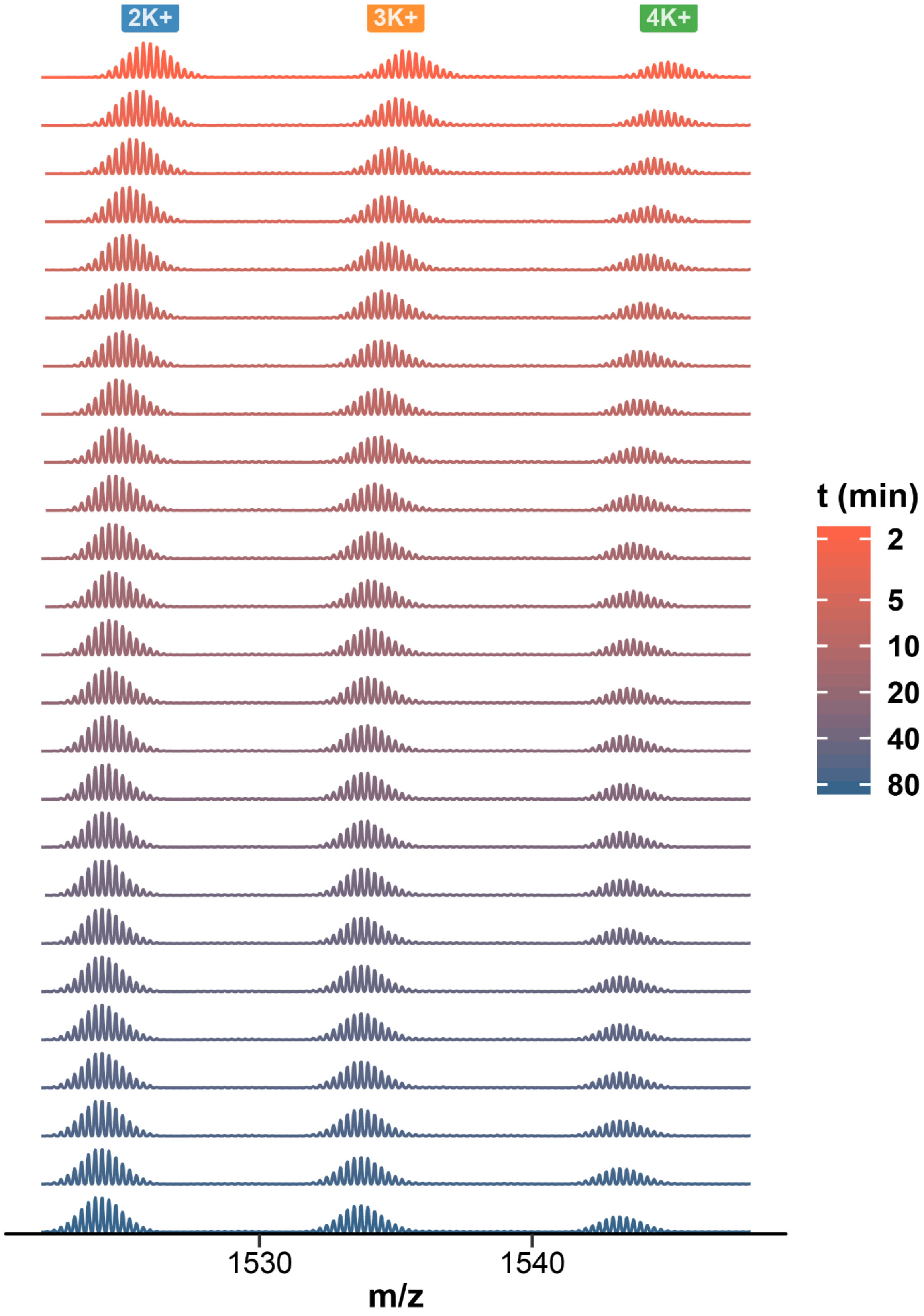


*Figure S37. HDX/native MS spectra of T30177-TT, zoomed on the 4- charge state. 25 exchange time points are presented from top to bottom. Replicate 2.*

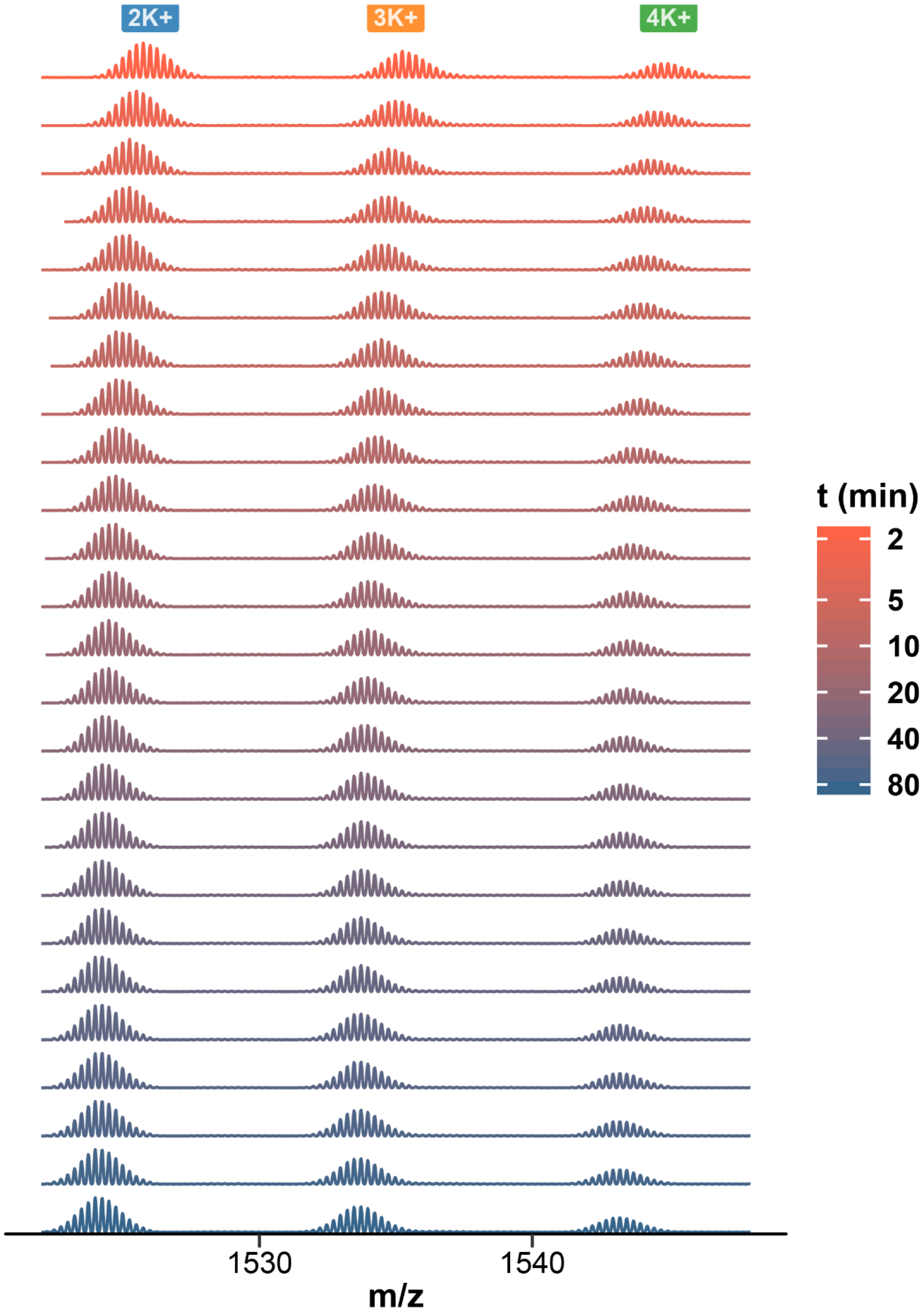


*Figure S38. HDX/native MS spectra of T30177-TT, zoomed on the 4- charge state. 25 exchange time points are presented from top to bottom. Replicate 3.*

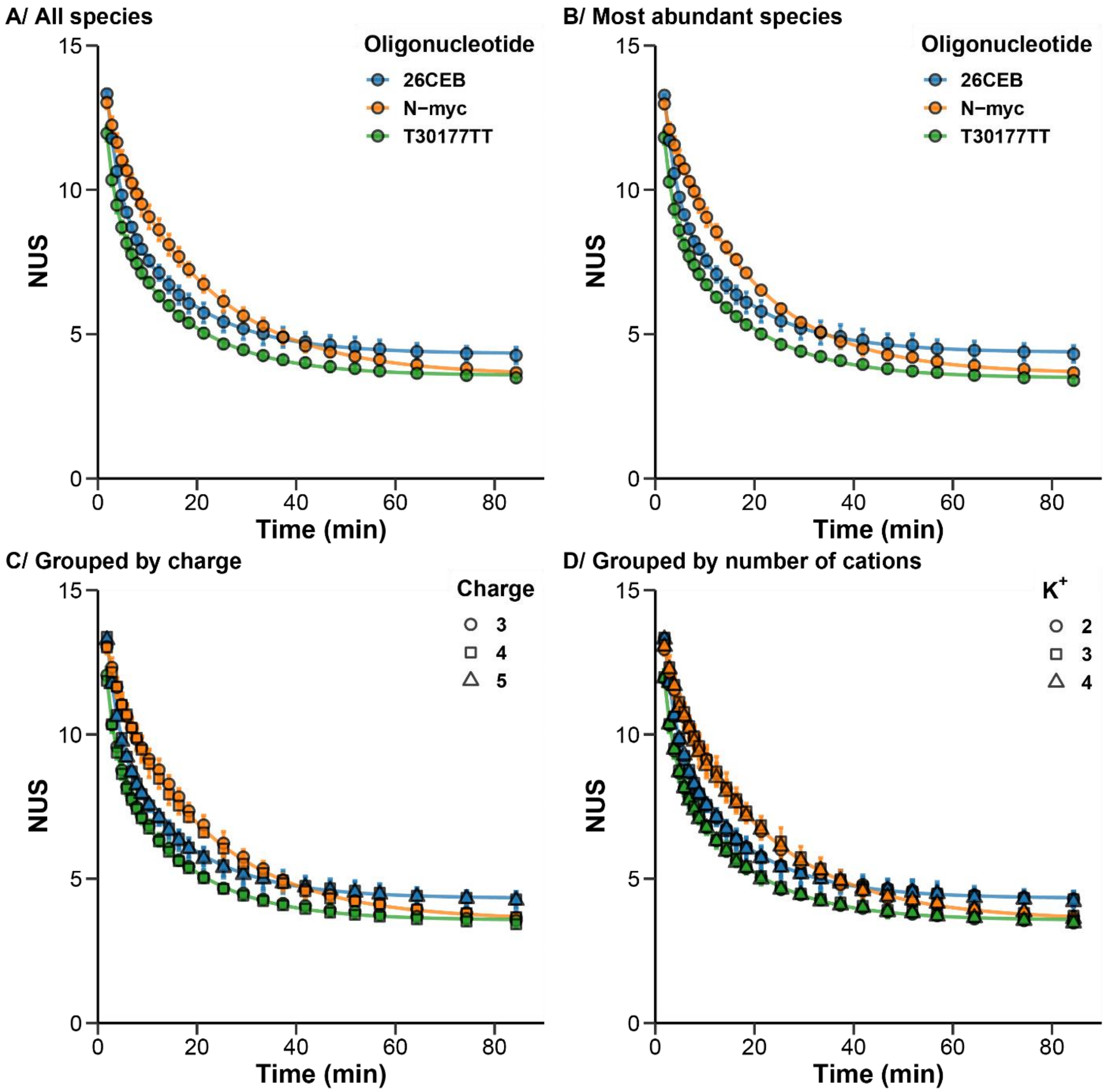


*Figure S39. Triplicated HDX/MS experiments for three oligonucleotides. The error bars show the standard deviation across all species and replicates (A), replicates of the most abundant species (B; $K^+$ = 2, z = 4-, except for 26CEB: 5-), replicates grouped by charge state (C), and replicates grouped by $K^+$ stoichiometry (D). The lines are the results of the non-linear fitting using Equation 4 with n = 2.*

| | Most abundant species | All species | Grouped by $K^+$ stoichiometry | Grouped by charge state |
|---|---|---|---|---|
| **26CEB** | 4.6 | 4.4 | 4.6 | 4.5 |
| **N-myc** | 2.7 | 3.7 | 3.7 | 3.6 |
| **T30177-TT** | 3.0 | 3.1 | 3.1 | 3.0 |

*Table S2. Mean RSD (%) on three replicates.*

| | Most abundant species | All species | Grouped by $K^+$ stoichiometry | Grouped by charge state |
|---|---|---|---|---|
| **26CEB** | 0.20 | 0.20 | 0.21 | 0.20 |
| **N-myc** | 0.15 | 0.22 | 0.22 | 0.21 |
| **T30177-TT** | 0.14 | 0.14 | 0.14 | 0.14 |

*Table S3. Mean SD in Da on three replicates.*

| Pairs | 26CEB/N-myc | 26CEB/T30177-TT | N-myc/T30177-TT |
|---|---|---|---|
| **Paired t-test *t*** | -5.2423 | 30.509 | 14.137 |
| **Paired t-test *p-value*** | $1.447 \times 10^{-6}$ | $< 2.2 \times 10^{-16}$ | $< 2.2 \times 10^{-16}$ |
| **Shapiro-Wilk *W*** | 0.92498 | 0.91285 | 0.92741 |
| **Shapiro-Wilk *p-value*** | 0.0002676 | 0.00007541 | 0.0003486 |
| **Wilcoxon *V*** | 632 | 2850 | 2850 |
| **Wilcoxon *p-value*** | 0.00001527 | $< 2.2 \times 10^{-16}$ | $< 2.2 \times 10^{-16}$ |

*Table S4. Statistical analysis of the significance of differences between pairs of oligonucleotides (most abundant species). The paired Student's t-test suggests that the differences within each pair are significant, but assumes that the differences are normally distributed. The Shapiro-Wilk normality test does not conclusively demonstrate that this is in fact the case. The Wilcoxon signed-rank test with continuity correction can be used instead, as it does not assume normality, yielding the same conclusions.*

| | $k_1$ (min$^{-1}$) | | | $k_2$ (min$^{-1}$) | | |
|---|---|---|---|---|---|---|
| | *2.5%* | *fit value* | *97.5%* | *2.5%* | *fit value* | *97.5%* |
| **26CEB** | 0.32 | 0.41 | 0.50 | 0.056 | 0.066 | 0.075 |
| **N-myc** | 0.10 | 0.85 | 1.61 | 0.050 | 0.053 | 0.056 |
| **T30177-TT** | 0.46 | 0.57 | 0.68 | 0.063 | 0.068 | 0.074 |

*Table S5. Apparent exchange rates and their 95% confidence interval (asymptotic method)*

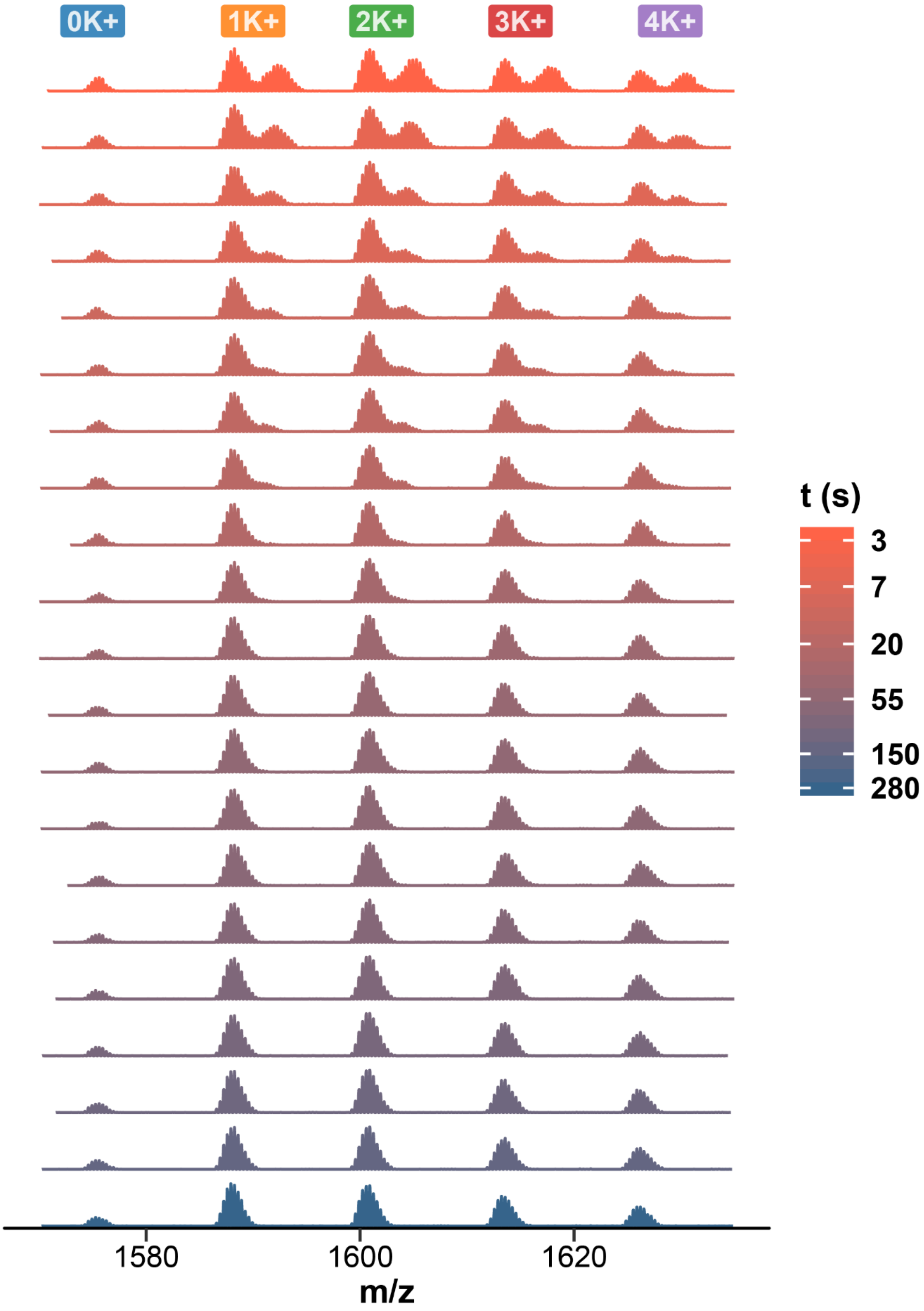


*Figure S40. Continuous-flow HDX/native MS spectra of TBA, zoomed on the 3- charge state. 21 exchange time points are presented from top to bottom.*

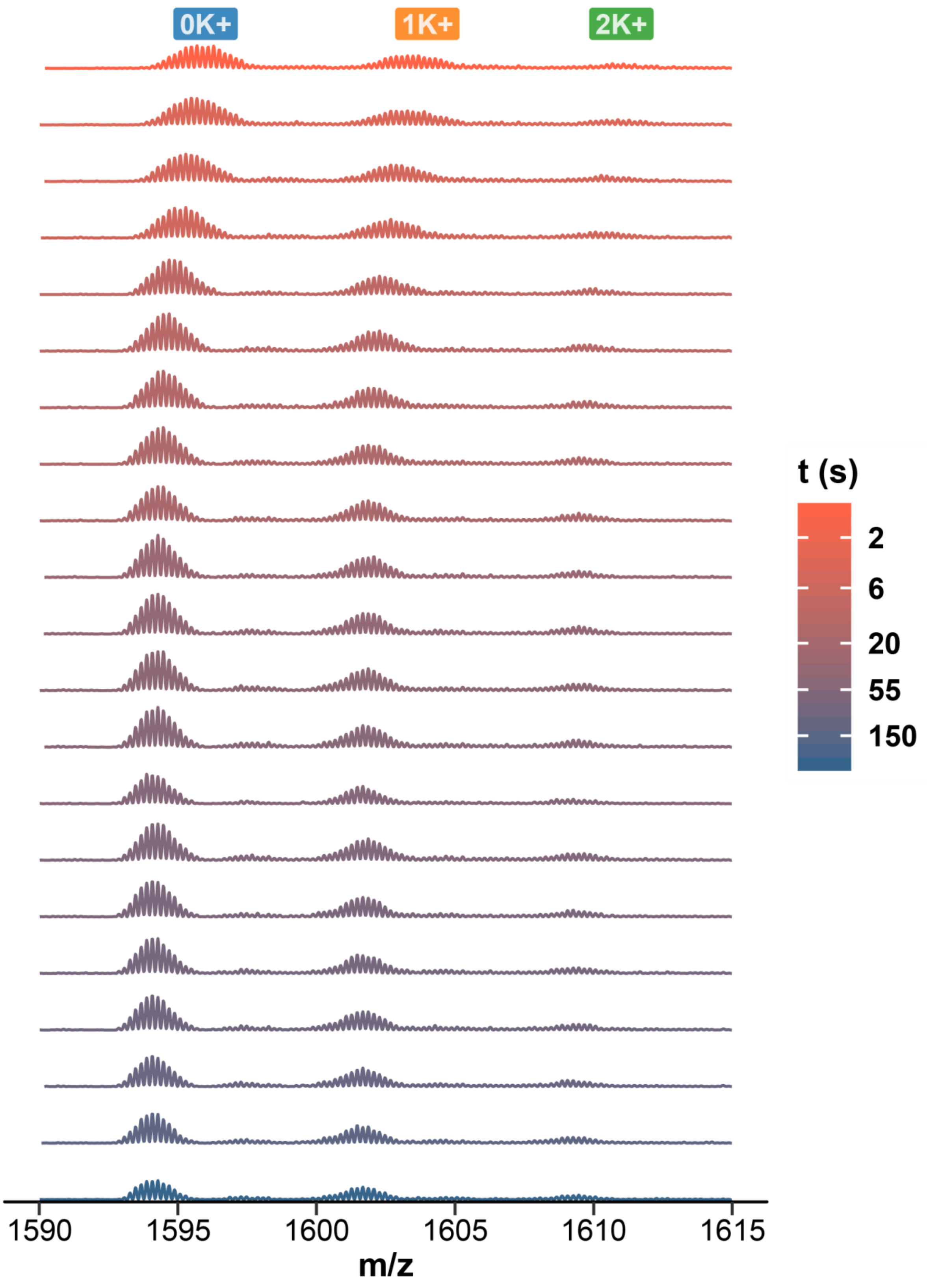


*Figure S41. Continuous-flow HDX/native MS spectra of ds26, zoomed on the 5- charge state. 21 exchange time points are presented from top to bottom.*

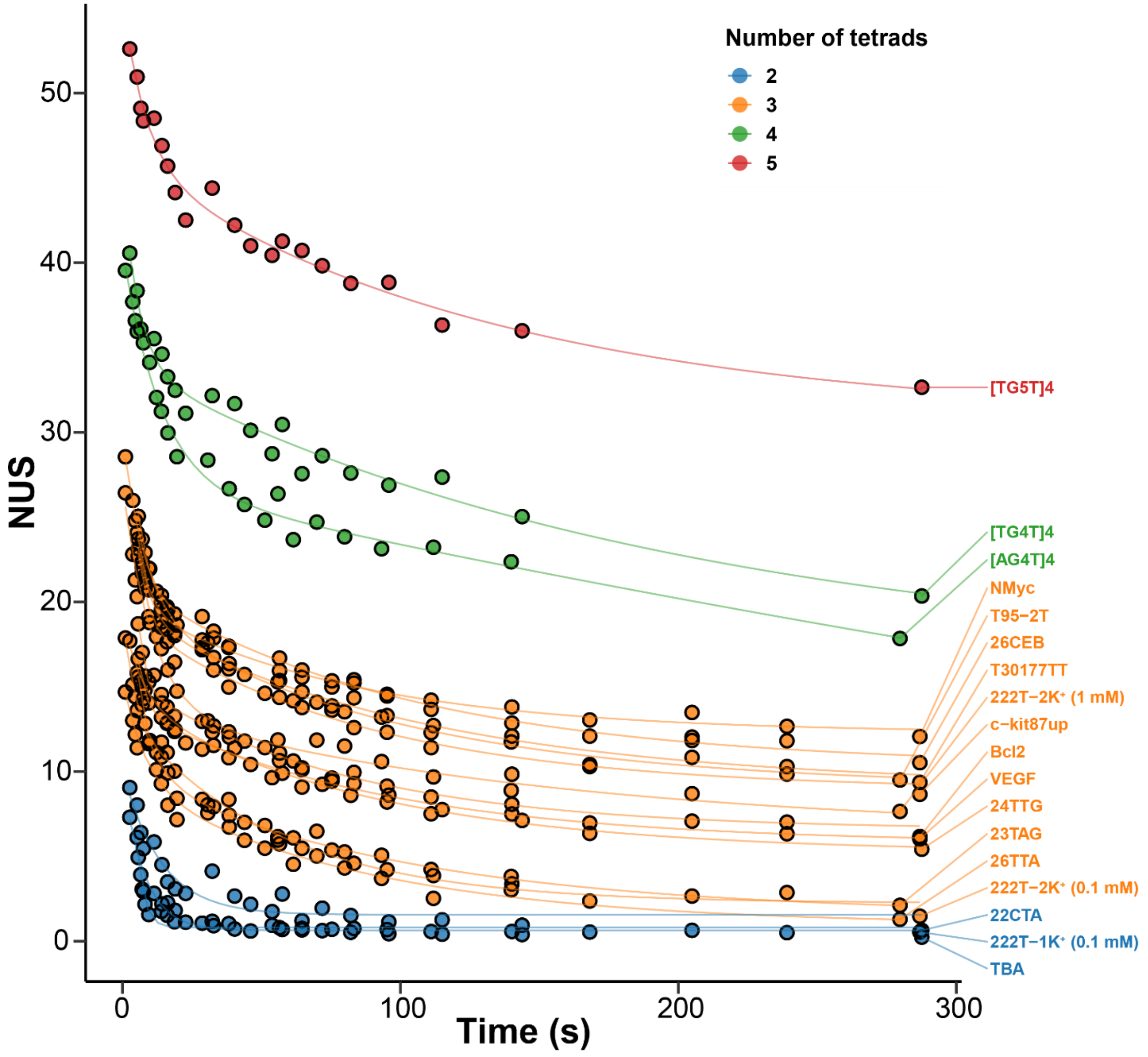


*Figure S42. Exchange kinetics of a panel of G-quadruplex forming oligonucleotides acquired by continuous-flow HDX native MS (90% to 9% D). Oligonucleotides are colored by number of tetrads, thereby indicating similar number of exchangeable sites.*
